# DTractor enhances cell type deconvolution in spatial transcriptomics by integrating deep neural networks, transfer learning, and matrix factorization

**DOI:** 10.1101/2025.04.12.648541

**Authors:** Yong Jin Kweon, Chenyu Liu, Gregory Fonseca, Jun Ding

## Abstract

Spatial transcriptomics (ST) captures gene expression with spatial context but lacks single-cell resolution. Single-cell RNA sequencing (scRNA-seq) offers high-resolution profiles without spatial information. Accurate spot-level decomposition requires effective integration of both. We present DTractor, a deep learning-based framework that improves cell-type deconvolution in ST data through spatial constraints and transfer learning. DTractor achieves dual utilization of scRNA-seq reference data by incorporating both a cell-type-specific gene expression matrix and learned latent embeddings into a unified matrix factorization model. This joint modeling enables accurate estimation of cell-type proportions and cell-type-resolved gene expression within each spatial spot, while preserving biological and spatial coherence. DTractor further applies spatial regularization to maintain local tissue structure. Across multiple ST platforms and tissue types, DTractor demonstrates improved decomposition accuracy, robustness, and interpretability compared to existing methods. The results from DTractor support downstream applications such as spatial domain analysis and the study of spatially organized cellular behaviors.

## Introduction

Understanding gene expression at the cellular level is essential for decoding tissue structure, function, and disease mechanisms. While bulk RNA sequencing provides average transcriptomic profiles across heterogeneous cell populations [1], single-cell RNA sequencing (scRNA-seq) captures cell-specific expression patterns at high resolution [2, 3], revealing diverse cell states and dynamic processes [4, 5]. However, scRNA-seq requires tissue dissociation, disrupting the spatial architecture necessary to study cell-cell interactions and anatomical organization [6–8]. Spatial transcriptomics (ST) technologies aim to address this limitation by capturing gene expression within intact tissue sections. Platforms such as 10x Visium [9], Slide-seq v2 [10], and Spatial Transcriptomics [11] preserve spatial context while profiling gene expression across spatially bar-coded spots. However, these spots typically contain mixtures of multiple cell types, requiring computational deconvolution to infer the underlying cellular composition [12–17].

Several methods have been developed to integrate scRNA-seq references with spatial transcriptomics (ST) data, primarily to estimate cell type proportions in spatial spots. Linear decomposition tools such as RCTD [14], SPOTlight [17], and Stereoscope [15] use average gene expression profiles from the reference and treat each spatial spot as a mixture of fixed signatures. These models often overlook complex gene expression variation and are sensitive to incomplete or imbalanced cell type representation in the reference. Probabilistic approaches such as DestVI [18], Cell2location [19], and Redeconve [20] incorporate latent variable modeling to capture uncertainty and improve robustness, but their inference procedures can be computationally demanding and may not scale efficiently to large tissue sections. Spatially informed methods like CARD [16] and Tangram [21] introduce spatial priors or alignment strategies, yet they often depend on rigid spatial assumptions that may not generalize well across tissue types. In addition, most existing methods rely on a single aspect of the scRNA-seq reference, such as cell type level average expression, without leveraging the full latent structure or aligning representations across data modalities. These limitations can reduce the biological coherence of the results and hinder generalizability across complex tissue architectures.

To address these challenges, we present DTractor, a deep learning based framework that integrates spatial constraints and transfer learning to improve cell type deconvolution in spatial transcriptomics. A key innovation of DTractor is its dual use of scRNA-seq reference data to guide the decomposition of spatial gene expression. First, it incorporates average gene expression profiles for each cell type as one component of a structured matrix factorization model, providing cell type specific transcriptional anchors. Second, it learns low dimensional latent embeddings of cell types from scRNA-seq data using a deep neural network, and constrains the embeddings inferred from spatial data to remain aligned. This alignment enforces biological consistency across modalities and captures both observed expression patterns and latent transcriptional structures. Both components, the explicit expression profiles and the aligned latent embeddings, are integrated into the matrix factorization process to estimate cell type proportions across spatial locations. This design enables DTractor to coherently combine spatial context and reference information, resulting in improved robustness, accuracy, and interpretability in spatial deconvolution.

We evaluate DTractor across multiple spatial transcriptomics datasets and demonstrate improved performance in accuracy, spatial coherence, and computational efficiency compared to existing methods, including CARD [16], Cell2location [19], RCTD [14], SPOTlight [17], Stereoscope [15], DestVI [18], Tangram [21], and Redeconve [20]. By enabling accurate and interpretable spatial deconvolution, DTractor can facilitate down-stream analyses such as tissue microenvironment profiling, spatial domain identification, and the discovery of cell-cell interactions, contributing to a deeper understanding of tissue biology in development and disease.

## Results

### DTractor method overview

DTractor is a deconvolution framework that integrates deep neural networks, transfer learning, and matrix factorization to improve the resolution and accuracy of spatial transcriptomics (ST) decomposition. It takes as input a single-cell or single-nucleus RNA-seq (sc/snRNA-seq) reference and an ST dataset to estimate cell type proportions across spatial locations. A key innovation of DTractor is its dual utilization of the scRNA-seq reference data, which is used to extract both a cell type–specific gene expression matrix and low-dimensional latent embeddings of each cell type. These two components are integrated into a unified framework to guide decomposition at both the gene and representation levels. In the first p hase ( Fig. 1a, b), variational autoencoders (VAEs) with zero-inflated negative binomial (ZINB) likelihoods are trained separately on the scRNA-seq and ST datasets to learn biologically meaningful latent representations, and a cross-modal alignment constraint ensures consistency between the two. In the second phase (Fig. 1c), the expression matrix is constructed by averaging gene expression across annotated cells of each type in the reference and serves as a fixed component during decomposition. In the third phase (Fig. 1 d), DTractor formulates deconvolution as an iterative matrix factorization problem, combining the reference expression matrix and the learned latent spot embeddings to model observed spatial expression. Optimization is performed using the Adam optimizer, with spatial regularization encouraging smoothness across neighboring spots and embedding regularization maintaining cross-modality alignment. The resulting cell type proportion estimates (Fig. 1e) enable downstream analyses such as spatial visualization, marker gene identification, and pathway enrichment, providing biologically interpretable insights into tissue structure and function.

**Fig. 1.**
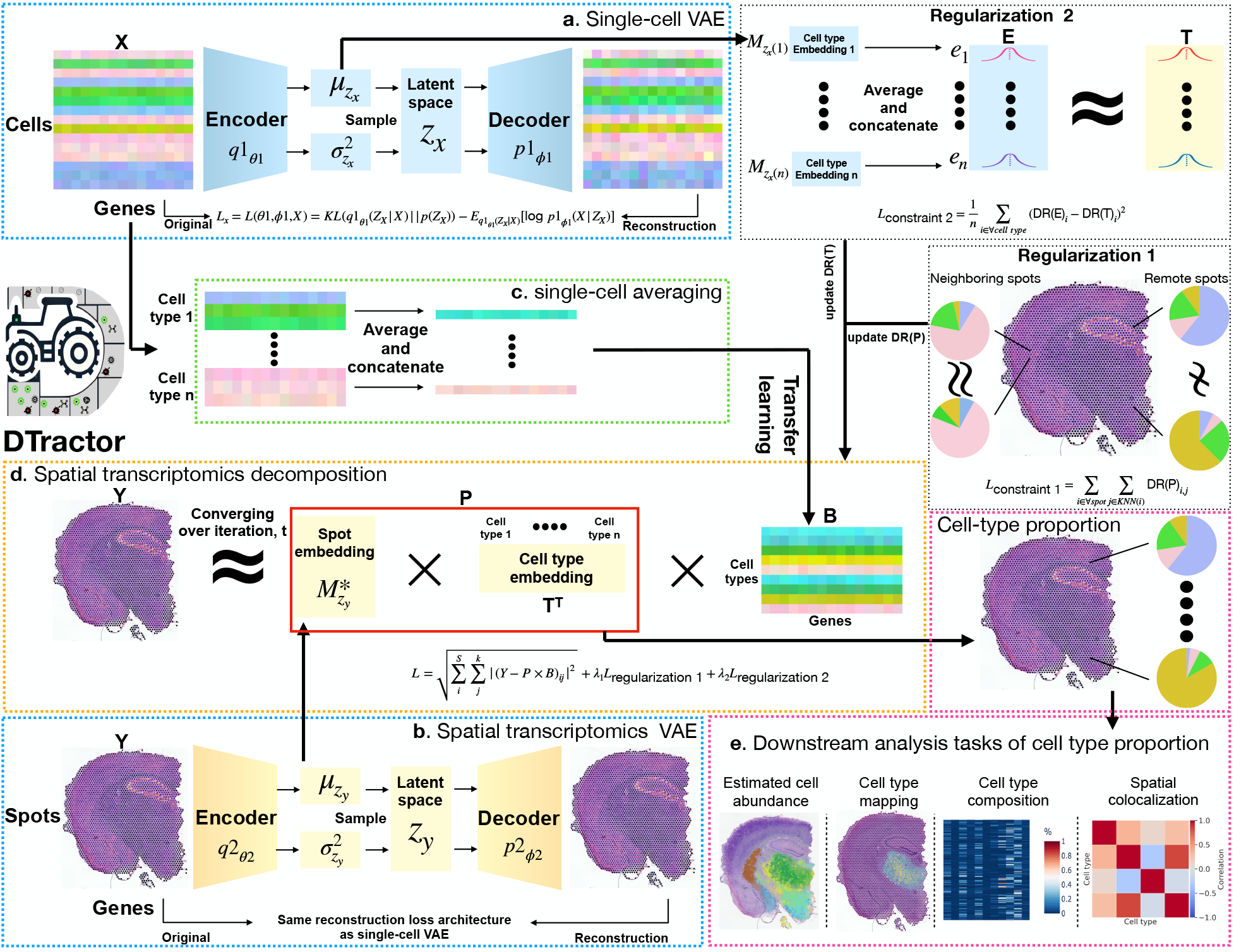
Overview of the DTractor Framework. DTractor integrates sc/snRNA-seq (*X*) and spatial transcriptomics (ST, *Y*) data to estimate cell-type proportions by combining deep representation learning, transfer learning, and structured matrix factorization. **a, b**, Variational autoencoders (VAEs) are independently trained on sc/snRNA-seq and ST data to learn latent representations 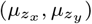, capturing biological variation while modeling sparsity and overdispersion with a zero-inflated negative binomial (ZINB) likelihood. **c**, A cell-type-specific gene expression matrix (*B*) is constructed by averaging expression profiles from annotated scRNA-seq data, providing expression-level guidance for decomposition. **d**, DTractor performs deconvolution through iterative matrix factorization, using both the reference matrix (*B*) and the aligned latent embeddings 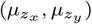 to estimate cell-type proportions (*P*) for each spatial spot. Optimization is conducted via Adam-based gradient descent, with spatial and cross-modal regularization to ensure local smoothness and modality alignment. **e**, The inferred cell-type proportions support downstream analyses, including spatial visualization, marker gene identification, and pathway enrichment.

### DTractor accurately deconvolutes germinal center and T-cell zones within the dynamically evolving microenvironment of the human lymph node

We benchmarked DTractor against eight established methods (CARD, Cell2location, DestVI, RCTD, SPOTlight, Stereoscope, Tangram, and Redeconve) using three published spatial transcriptomics (ST) datasets: a Visium human lymph node dataset (10X Genomics) [19, 22–24], a human pancreatic ductal adenocarcinoma (PDAC) dataset [25], and a mouse olfactory bulb (MOB) dataset [11, 26]. This analysis focuses on the human lymph node dataset, comprising 4,035 spatial locations with diverse microenvironments and intersecting cell populations. The reference dataset consisted of single-cell RNA sequencing (scRNA-seq) data from human secondary lymphoid organs, identifying 34 cell types (e.g., B cells, dendritic cells, macrophages, and T cells) via standard preprocessing, UMAP projection initialized by PAGA, and Leiden clustering (Fig. 2a).

**Fig. 2.**
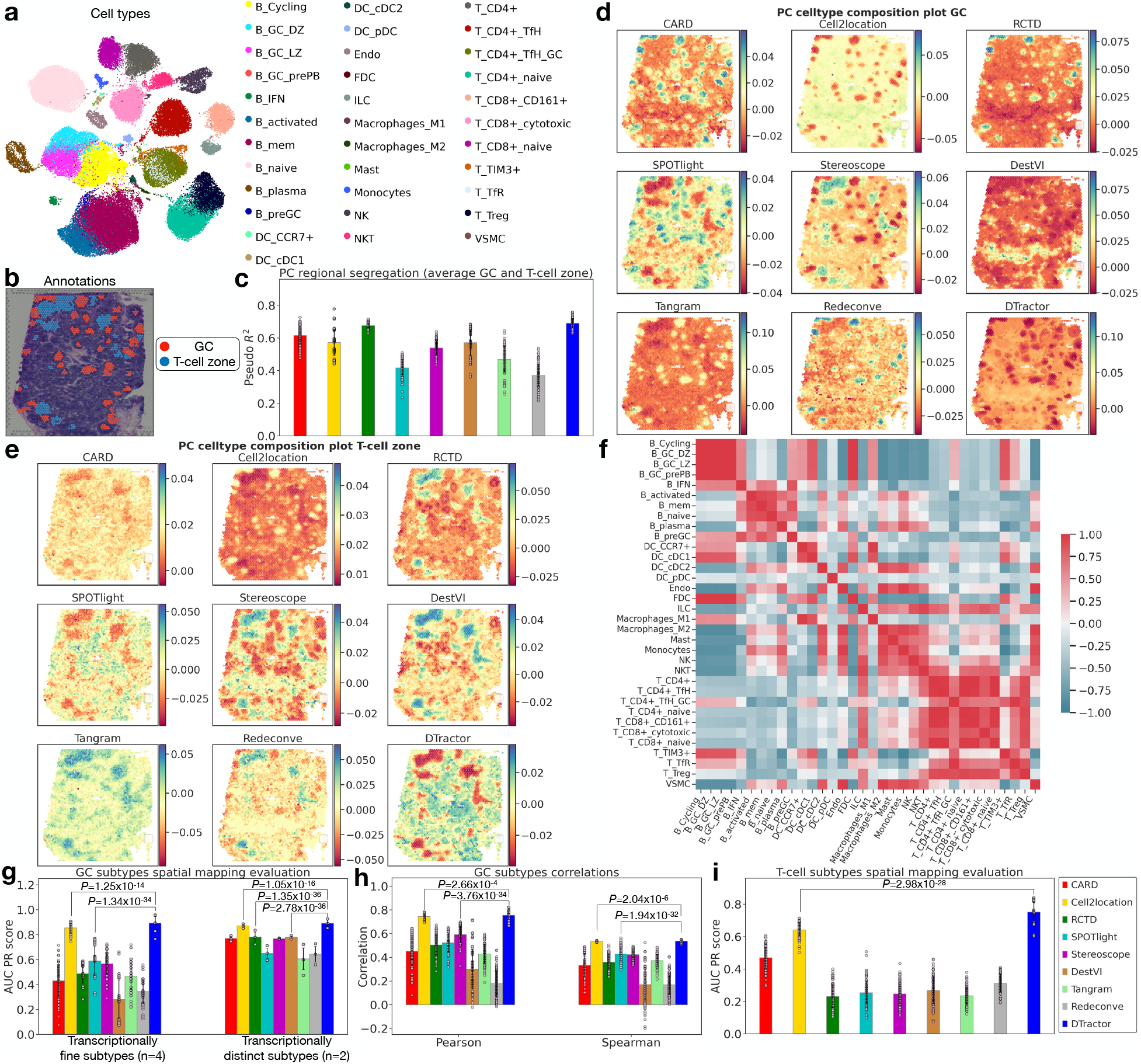
DTractor provides detailed characterization of germinal center (GC) and T-cell zones in the dynamic microenvironment of the human secondary lymph node. **a**, UMAP representation, initialized using a partition-based graph abstraction (PAGA) graph derived from 34 reference cell types in the integrated human secondary lymphoid organ scRNA-seq dataset, with cell type labels informed by Leiden clustering. **b**, Ground truth annotations for GC and T-cell regions were obtained from Hematoxylin and Eosin (H&E) staining, as described in the Redeconve paper, and used for benchmarking. **c**, Quantitative evaluation of the principal components (PCs) of the estimated cell type composition matrix shows distinct segregation between GC and non-GC (**d**) and T-cell vs. non-T-cell zones (**e**). Logistic regression models trained on PC scores yield average *R*^2^ values for GC and T-cell zone classifications. **d, e**, PCs with the highest *R*^2^ for GC (**d**) and T-cell zone (**e**) ground truth annotations were identified, with *R*^2^ calculated for the first three PCs. **f**, DTractor infers correlations in cell type composition across spatial locations, revealing colocalization of GC- and T-cell-related cell types. **g**, Spatial mapping accuracy for six GC-associated cell types is evaluated using macro-average area under the precision-recall curve (AUC-PR) scores, highlighting transcriptionally fine subtypes (B_GC_DZ, B_GC_LZ, B_GC_prePB, T_CD4+_TfH_GC) and distinct subtypes (B_cycling, FDC). **h**, Macro-average correlation coefficients between the estimated cell type composition matrix and GC ground truth annotations are reported for six GC cell types. **i**, Spatial mapping accuracy for nine T-cell types is evaluated using macro-average AUC-PR scores based on T-cell zone ground truth annotations. **c, g-i**, P-values are from one-sided Mann-Whitney U tests without adjustment, based on 100 bootstrap resamples. Bars represent average statistics across resamples with error bars indicating standard errors. Results reflect three preprocessing conditions (**c**), six GC subtypes (**g, h**), and nine T-cell types (**i**).

We first evaluated each method’s capacity to resolve germinal centers (GC) and T-cell zones, validated histologically (Fig. 2b, c). To capture essential cellular spatial relationships while simplifying the high-dimensional data, we employed principal component analysis (PCA) on the inferred cell type composition matrices as detailed in Methods. For each method, rather than assuming the first two PCs would contain the biologically relevant signals for GC and T-cell zones, we selected the component that best discriminated between GC and T-cell zones based on the highest Nagelkerke R^2^ value from logistic regression. This approach identified the most informative component for our biological question, as tissue microenvironment-related variance may distribute across different components depending on the method. The selected components (PC_*GC*_ and PC_*T*_) were then used for subsequent analyses. DTractor uniquely maintained robust segregation of both regions (*R*^2^ > 0.65) across variable preprocessing conditions (e.g., gene filtering stringency yielding 4,423–7,667 shared genes) and bootstrap resampling. While CARD and Cell2location performed well in detecting GC regions (*R*^2^ > 0.7) but struggled with T-cell zones (*R*^2^ < 0.5), DestVI effectively identified T-cell zones (*R*^2^ > 0.65) but underperformed in detecting GC regions (*R*^2^ < 0.5) (Fig. 2c, Supplementary Fig. S1). In PC_*GC*_ plots (Fig. 2d), DTractor distinctly resolved the GC’s layered structure, whereas DestVI failed to differentiate it, and RCTD, SPOTlight, and Redeconve exhibited indistinct boundaries. Likewise, in PC_*T*_ plots (Fig. 2e), DTractor sharply delineated T-cell zones, while CARD, Cell2location, SPOTlight, Stereoscope, Tangram, and Redeconve showed poor resolution or blurred demarcations.

Subsequently, we scrutinized immune cell populations specific to the GC to map transcriptionally fine-grained cell types, considering the dataset’s abundance of nuanced subtypes with specialized immune functions [19, 27–29]. DTractor precisely colocalized six established GC subtypes (B_GC_DZ, B_GC_LZ, B_GC_prePB, T_CD4+_TfH_GC, B_cycling, FDC) (Fig. 2f), outperforming other methods in average precision-recall area under the curve (AUC PR) scores (Fig. 2g). In this context, the classification of six known GC cell types is determined by their transcriptional characteristics, distinguishing between those that are fine-grained (B_GC_DZ, B_GC_LZ, B_GC_prePB, T_CD4+ _TfH_GC) and those that are distinct (B_cycling, FDC) [19]. For fine-grained subtypes, DTractor and Cell2location excelled, whereas DestVI and Redeconve underperformed. For distinct subtypes, DTractor surpassed all methods (P=1.05×10^−16^ compared to the second-best, Cell2location), achieving 89% accuracy, confirmed through precision-recall scores for each GC subtype (Supplementary Figs. S2, S3). By contrast, SPOTlight, Tangram, and Redeconve showed reduced accuracy for distinct subtypes. In cell type proportion estimation (Fig. 2h), DTractor outperformed all other methods in correlation accuracy (Pearson P=2.66×10^−4^ and Spearman P=2.04×10^−6^ compared to the second-best, Cell2location) with strong consistency across bootstrapping iterations (Supplementary Fig. S4), uniquely recapitulating all six GC subtypes spatially (Supplementary Fig. S5), consistent with previous studies [19, 30].

Finally, we evaluated T-cell zone mapping across nine T-cell subtypes (Fig. 2i). DTractor and Cell2location yielded satisfactory AUC PR scores, with DTractor exhibiting superior performance (0.77 vs. 0.64, P=2.98×10^−28^) (Supplementary Fig. S6). Spatial distribution analyses corroborated this, highlighting DTractor’s exceptional resolution of T-cell subtypes (Supplementary Fig. S7). Additionally, T_TIM3+ cells, characterized by the expression of T-cell immunoglobulin and mucin protein 3 (TIM3)—a marker commonly associated with T cell exhaustion [31]—might be less prevalent in T-cell zones due to their impaired functionality [32, 33]. Exclusion of T TIM3+ cells, associated with T-cell exhaustion, enhanced mapping accuracy (Supplementary Figs. S8, S9), though DTractor accurately captured their GC colocalization in contexts of chronic activation [34–36]. Correlation analyses of estimated versus ground-truth T-cell proportions further validated DTractor’s robustness (Supplementary Figs. S10, S11), particularly when excluding T_TIM3+ (Supplementary Fig. S12). Additionally, DTractor effectively captured the distinct spatial distribution of T_TIM3+ cells, demonstrating reduced co-localization with other T cell subtypes (Fig. 2f) and lower enrichment in T cell zones, reflected in their lower AUC PR scores (Supplementary Figs. S7, S13, S14). This biologically relevant pattern was uniquely captured by DTractor, while other methods failed to detect this important cell distribution characteristic. Furthermore, T follicular regulatory cells (T_TfR), which are initially generated in the T-cell zones and later migrate to the GC to regulate the activity of T follicular helper (TfH) cells and B cells [37, 38], were also accurately depicted in the plots. Thus, DTractor detected reduced colocalization of T_TfR and minimal colocalization of T_TIM3+ with other T cell subpopulations, while also identifying colocalizations of T_TfR and T_TIM3+ with GC subtypes, consistent with previous studies.

DTractor’s ability to address spatially heterogeneous cell distributions, such as the contrasting patterns of dispersed T cells versus clustered B cells [39], likely contributes to its enhanced performance. Additionally, it demonstrated exceptional memory efficiency, maintaining consistently low memory usage around 500 MB across all tested gene sets, over 10x reduction compared to other methods consuming 5,000-17,500 MB, while delivering manageable computation times around 3,000 seconds (Supplementary Fig. S15).

### DTractor provides a comprehensive portrayal of the four distinct tissue regions within human PDAC

We evaluated DTractor and eight other methods using a human pancreatic ductal adenocarcinoma (PDAC) spatial transcriptomics (ST) dataset from the original Spatial Transcriptomics platform. The reference dataset, matched single-cell RNA sequencing (scRNA-seq) data from the same sample via inDrop, comprised 1,926 cells across 20 cell types (e.g., acinar, cancer, ductal, fibroblasts, macrophages). Standard preprocessing, Leiden clustering, PAGA initialization, and UMAP visualization identified these cell types (Fig. 3a). The ST dataset included 428 spots spanning four histologically annotated regions—cancer cells with desmoplasia, nonmalignant duct epithelium, stroma, and normal pancreatic tissue—validated by HE staining (Fig. 3b) [20, 25].

**Fig. 3.**
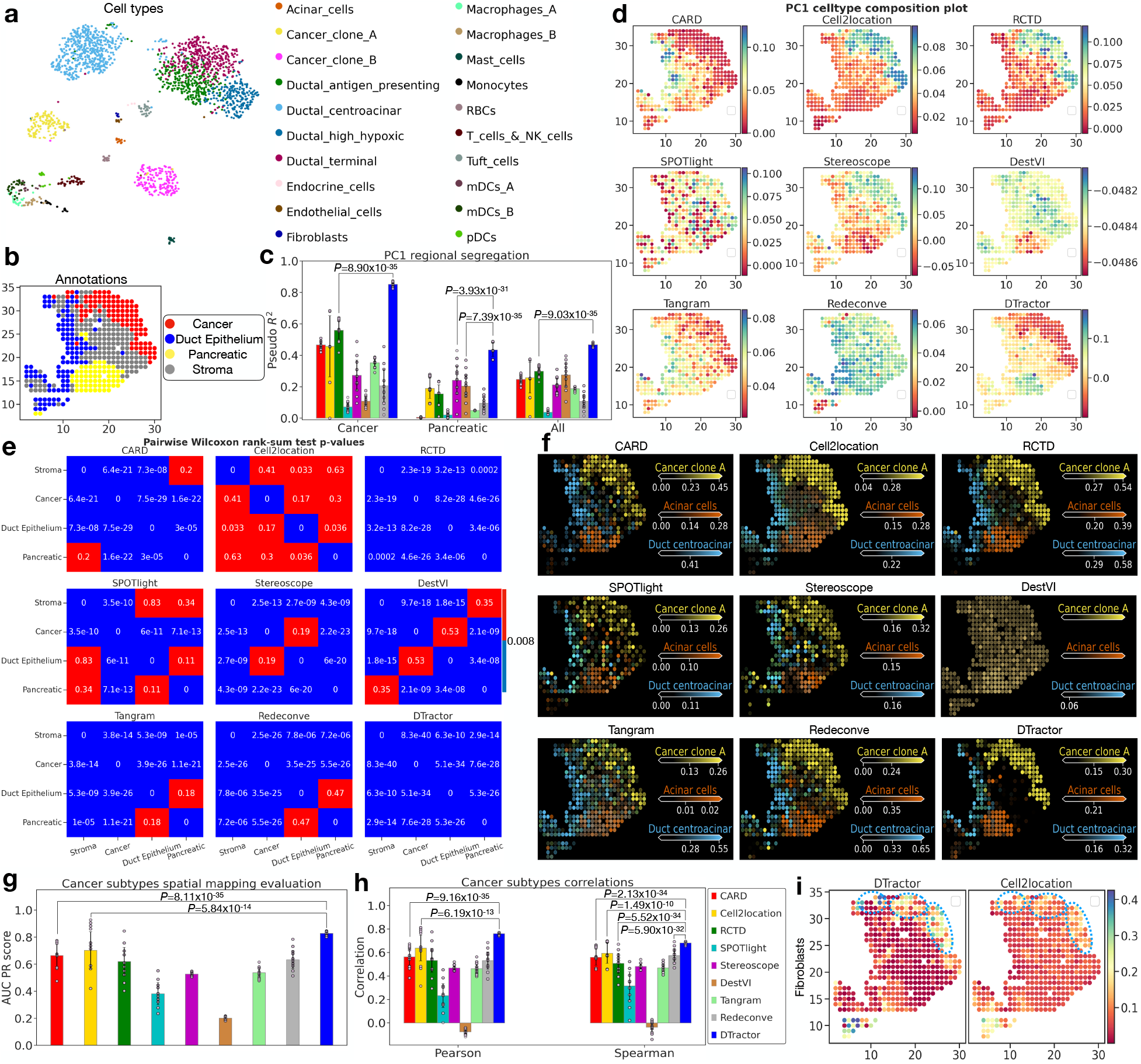
DTractor provides a detailed characterization of four tissue regions in human pancreatic ductal adenocarcinoma (PDAC): cancer, ductal, pancreatic, and stromal regions. **a**, UMAP representation initialized using a partition-based graph abstraction (PAGA) graph of 20 reference cell types derived from an integrated scRNA-seq dataset, with cell types quantified via Leiden clustering. Each model is trained to decompose spatial transcriptomics (ST) spots into these cell types. **b**, Ground truth annotations for the four tissue regions were obtained from Hematoxylin and Eosin (H&E) staining, as described in the original and Redeconve papers. **c**, Quantitative evaluation of the first principal component (PC1) from the estimated cell type composition matrix (**d**) reveals regional segregation between cancer and non-cancer regions, pancreatic and non-pancreatic regions, and all four tissue regions. Logistic regression models were fitted for binary and multinomial classifications using PC1. **d**, PC1 of the estimated cell type composition matrix, highlighting its explanatory power. **e**, Heatmap of pairwise Wilcoxon rank-sum tests using PC1 to assess differentiation between the four tissue regions, with significant p-values (adjusted threshold: 0.008) colored blue. Each box represents the p-value indicating significant differentiation between each pair. **f**, Cell type abundance for major tissue zones (cancer, ductal, and pancreatic regions) estimated using DTractor and visualized via Cell2location. **g**, Spatial mapping accuracy for three cancer-associated cell types (Cancer clone A, Cancer clone B, Fibroblasts) evaluated via macro-average area under the precision-recall curve (AUC-PR) scores. **h**, Macro-average correlation coefficients between the estimated cell type composition matrix and cancer ground truth annotations are reported for three cancer-associated cell types. **i**, Estimated fibroblast abundance demonstrates consistency with prior findings, identifying fibroblasts primarily in the upper cancer region with minimal false positives. Benchmarking highlights DTractor and Cell2location as the top-performing models (**g, h**). **c, g-h**, P-values are derived from one-sided Mann-Whitney U tests without adjustment, using 100 bootstrap resamples. Bars represent average statistics across resamples, with error bars indicating standard errors. Results include three preprocessing conditions (**c**) and three cancer-associated subtypes (**g, h**).

First, we evaluated deconvolution performance by assessing how accurately each method captured known tissue architecture. Using PCA on inferred cell type compositions, we reduced dimensionality while preserving key cellular distribution patterns. This approach converts complex multi-cell-type profiles into principal components that capture major sources of variation, with the first principal component (PC1) typically representing the strongest tissue-specific signal. To quantify how accurately these cellular patterns correspond to known tissue regions, we calculated the *R*^2^ between PC1 scores and annotated regions (cancer, pancreatic, and all four regions), testing robustness across three preprocessing strategies (shared genes: 8,729–13,402). Higher *R*^2^ values indicated better discrimination between tissue regions. DTractor achieved 86% accuracy for cancer regions with low standard error (±0.02), far surpassing RCTD (55%±0.08; 8.90×10^−35^), while SPOTlight, Stereoscope, DestVI, and Redeconve missed less than 30% of cancer areas (Fig. 3c). For pancreatic and all four regions, DTractor consistently outperformed others, with Cell2location, Stereoscope, DestVI, and Redeconve showing higher variability. Bootstrapping (10–500 iterations) reinforced DTractor’s superiority (Supplementary Fig. S16). Subsequently, PC1 visualization (Fig. 3d) revealed DTractor’s distinct segregation of cancer versus non-cancer and pancreatic versus non-pancreatic regions, identifying subtle regional features like cancer tissue at (13, 23) and pancreatic tissue at (14, 25) and (15, 25). It uniquely detected peripheral cancer regions (e.g., (8, 33), (11, 34), (14, 28), (26, 24), and (29, 19)), whereas SPOTlight and Redeconve detected no regions, DestVI failed on cancer areas, and CARD, Cell2location, RCTD, Stereoscope, and Tangram showed incomplete resolution.

Next, we assessed DTractor’s ability to discriminate between distinct tissue regions using pairwise two-sided Wilcoxon rank-sum tests [40] across four known tissue regions: stroma, cancer, duct epithelium, and pancreatic tissue (Fig. 3e). The heatmaps display p-values for each pairwise comparison, with blue cells indicating statistically significant differences (p < 0.008, Bonferroni-corrected threshold) and red cells showing non-significant differences. DTractor and RCTD successfully differentiated all tissue region pairs with high statistical significance, aligning with histological annotations. DTractor yet exhibited superior statistical power, distinguishing pancreatic and stroma regions at P=2.9×10^−14^, compared to RCTD’s P=0.0002 for the same pair. In contrast, Cell2location failed to differentiate all pairs, and SPOTlight missed half of them. These results demonstrate the superior discriminative power of our approach in identifying biologically distinct pancreatic tissue regions. Across varying shared genes, DTractor consistently captured relative regional distances, a pattern absent in most methods (Supplementary Figs. S17, S18, S19). The plots reveal that cancer and pancreatic regions are the farthest apart, with stroma and duct epithelium positioned in between, aligning with the ground truth annotations. Notably, only DTractor consistently maintained this pattern across varying numbers of shared genes.

Transitioning to cell type localization (Fig. 3f), DTractor accurately assigned “Cancer clone A”, “Acinar cells”, and “Ductal centroacinar” to their respective red-labeled cancer, yellow-labeled pancreatic, and blue-labeled duct epithelium regions (Fig. 3b) as reported in the original study [25]. In contrast, DestVI lacked spatial coherence, showing homogeneous Cancer clone A distribution throughout the tissue with minimal distinction between cell types. CARD, SPOTlight, Stereoscope, and Tangram showed moderate accuracy but significant signal bleeding, with ductal or acinar signals extending into stroma. Cell2location, RCTD, and Redeconve mapped cancer effectively but displayed false ductal and acinar signals in stroma and cancer regions. This comparative analysis reveals DTractor’s capacity to maintain tissue architecture fidelity while minimizing cross-region misassignments. Additionally, while correctly identifying the four main annotated regions in the dominant cell type inferred for each location, DTractor successfully captured the slight enrichment of monocyte cells in the stroma and, to a lesser degree, in the duct epithelium regions, as reported in a previous study [25], which none of the other methods achieved (Supplementary Fig. S20, S21, S22). This supports previous findings that monocytes and their derivative macrophages tend to accumulate in stromal regions [41, 42]. Their interaction with epithelial cells may facilitate tumor progression by secreting cytokines and growth factors [43]. This demonstrates DTractor’s unique capability to learn from a small number of cells (monocytes; n = 18) in the reference data.

To further explore cancer cell type mapping in PDAC spatial regions, we mapped three cancer-related cell types previously shown to be enriched in the cancer region [25]: “Cancer clone A,” “Cancer clone B,” and “Fibroblasts” (Fig. 3g). DTractor achieved 83% AUC PR score (P=5.84×10^−14^ compared to the second-best Cell2location) with notably low variability (SD = ±2.1%), significantly outperforming all other methods, with Cell2location reaching only 70%. This superiority was consistent across different bootstrap sample sizes (10, 50, and 500), where DTractor maintained an 81-84% AUC PR score while other methods showed greater performance fluctuations (Supplementary Fig. S23). Furthermore, correlation analyses between inferred cell type proportions and cancer region annotations quantitatively confirmed DTractor’s superiority, achieving the highest correlation coefficients (r > 0.68; Pearson P=6.19×10^−13^ and Spearman P=1.49×10^−10^ compared to the second-best, Cell2location) exceeding the next best method by 12% (Fig. 3h, Supplementary Fig. S24). Specifically, fibroblasts are known to be associated with advanced tumor-node-metastasis stages and abundant in pancreatic cancer [25, 44–46]. Previous studies suggest the upper cancer region is likely a late-stage cancer area with metastatic potential [16, 20]. Notably, fibroblasts are the only cell type specifically localized to the upper cancer region, making this observation particularly intriguing for further investigation [25]. Supporting these findings, our analysis using DTractor uniquely showed fibroblast enrichment in the upper cancer region, consistent with earlier reports (Fig. 3i, Supplementary Fig. S25, S26, S27).

Finally, DTractor revealed spatial correlations among inferred cell types. DTractor and Cell2location uniquely identified high colocalization of two cancer cell types and four ductal cell types, while DTractor alone detected minimal colocalization of high hypoxic ductal cells with other ductal subtypes, aligning with prior findings [25]. This supports hypotheses of cancer origination from ductal epithelium and spread to form tumors under hypoxic conditions [47, 48], with hypoxic ductal cells migrating toward stroma and cancer regions. DTractor also identified colocalization of endothelial cells and fibroblasts, consistent with their roles in pancreatic cancer stroma interaction [16, 49], and of acinar and endocrine cells, reflecting their shared origins and functionally interconnected relationships [25, 50] (Supplementary Fig. S28). Additionally, DTractor maintained exceptionally low memory usage of approximately 90 MB across all tested gene sets, representing a 10-50x reduction compared to other methods that consumed between 800-4000 MB, while delivering manageable computation times around 350 seconds and superior reconstruction accuracy. This remarkable memory efficiency combined with competitive processing speed makes DTractor particularly suitable for analyzing large spatial transcriptomics datasets where both resource optimization and accuracy are essential (Supplementary Fig. S29).

### DTractor demonstrates accurate deconvolution in the well-defined anatomical layers of the mouse olfactory bulb

We assessed DTractor and eight other methods using replicate 12 of a mouse olfactory bulb (MOB) spatial transcriptomics (ST) dataset from the Spatial Transcriptomics platform [11], comprising 282 spots. The reference dataset, scRNA-seq data from the same tissue via 10x Chromium [26], included 21,746 cells across 18 cell types, such as olfactory sensory neurons (OSNs), granule cells (GC), periglomerular cells (PGC), mitral and tufted cells (M/TC), external plexiform layer interneurons (EPL-IN), and immature, transitional, and astrocyte-like subtypes. Unlike Ma et al. [16], we retained all immature, transitional, and astrocyte-like subtypes for a comprehensive assessment. Standard preprocessing, Leiden clustering, PAGA initialization, and UMAP visualization identified these cell types (Fig. 4a). The ST dataset delineated four histologically annotated regions via H&E staining: granule cell layer (GCL), mitral cell and external plexiform layer (MCL/EPL), glomerular and external plexiform layer (GL/EPL), and olfactory nerve layer (ONL) (Fig. 4b). The MOB exhibits diverse neural cell types within well-defined spatial architectures, with the EPL forming a boundary between MCL and GL. The MOB’s intricate layering and the limited resolution of spatial spots complicate the precise delineation of smaller-scale regions, such as the internal plexiform layer (IPL) and rostral migratory system (RMS), which, though not explicitly annotated, may be present [16, 51–53].

**Fig. 4.**
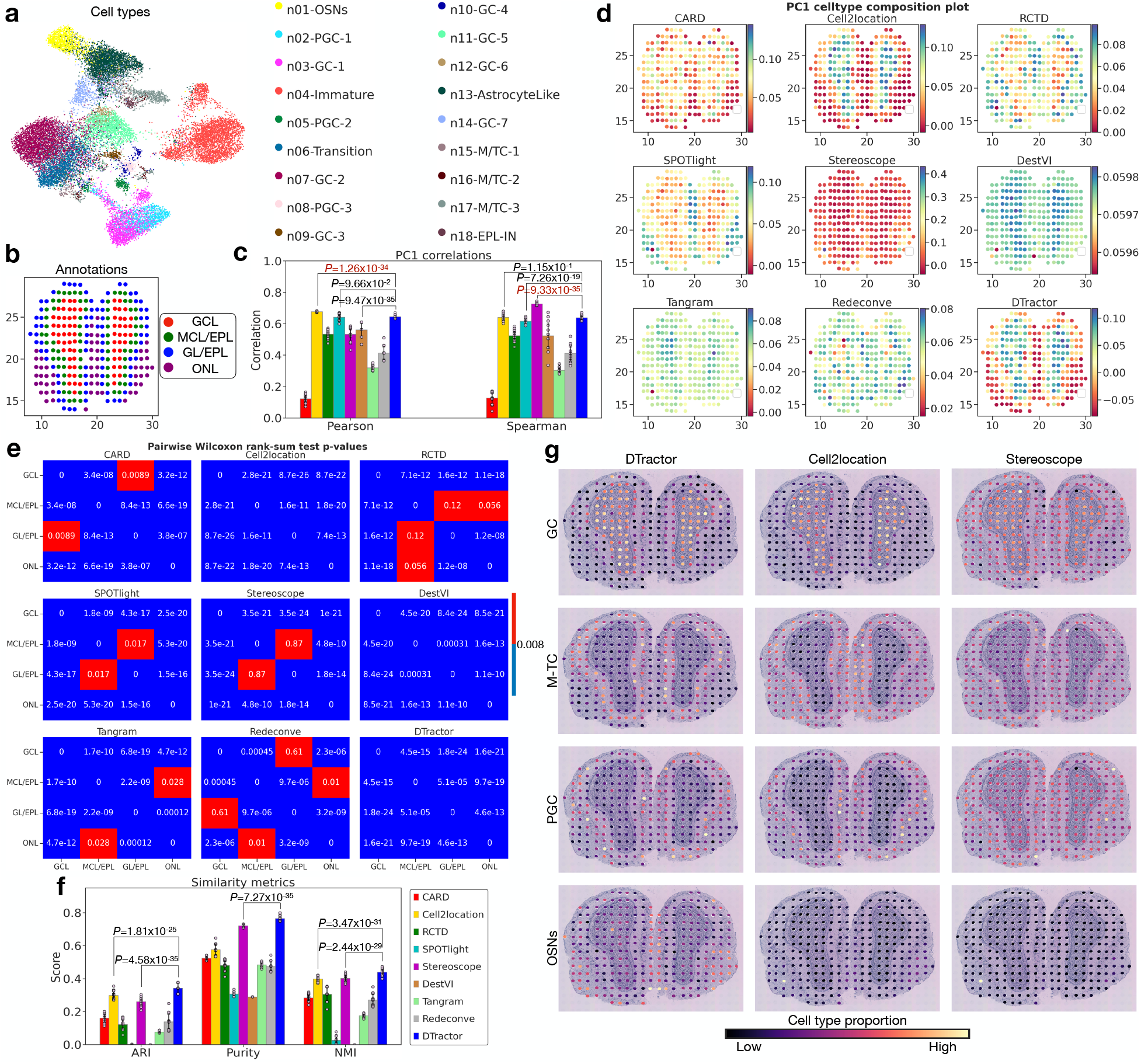
DTractor provides a detailed depiction of the four distinct anatomical regions within a mouse olfactory bulb (MOB): the granule cell layer (GCL), mitral cell layer/external plexiform layer (MCL/EPL), glomerular layer/external plexiform layer (GL/EPL), and olfactory nerve layer (ONL). **a**, The UMAP representation, with the x- and y-axes denoting the low-dimensional embedding, is initialized using the partition-based graph abstraction (PAGA) graph of 18 reference neuronal cell types (labels), forming the basis for training each model. These cell types are quantified based on the connectivity derived from Leiden clustering of the integrated 10x Chromium scRNA-seq reference dataset on the same tissue. **b**, Ground truth annotations for the four different regions in MOB were obtained from Hematoxylin and Eosin (H&E) staining data, as referenced in the original and CARD papers. **c**, Bar plots show Pearson and Spearman correlations between the first principal component (PC1) of the estimated cell type composition matrix (in **d**) and the ground truth annotation (in **b**) for various methods. **d**, The PC1 of the estimated cell type composition matrix. **e**, A heatmap of pairwise two-sided Wilcoxon rank-sum tests on PC1 of the estimated cell type composition matrix shows differentiation between the four regions, aligned with ground truth (in **b**). After applying the Bonferroni correction (p-value threshold of 0.05/6 = 0.008), p-values less than 0.008 are considered significant and are highlighted blue. **f**, Pairwise accuracy between the estimated dominant cell types and the ground truth annotation (in **b**) using adjusted rand index (ARI), normalized mutual information (NMI), and purity score. **g**, The proportion of one randomly selected example from each of the four cell types (GC, M-TC, PGC, and OSNs), utilizing the minimum number of shared genes (4,700), demonstrating DTractor’s robustness in learning the model with a limited gene set for training. We highlighted the three models with the highest similarity metric scores (in **f**): DTractor, Cell2location, and Stereoscope. **c, f**, P-values are from one-sided Mann-Whitney U tests without adjustment, based on 100 bootstrap resamples. Dark red indicates p-values where other models outperformed, meeting the statistical significance threshold of 0.05. Each bar’s height represents the average score from 100 resampled boostraps with three varying preprocessing, while scatter dots represent different resamples, and error bars denote the standard errors.

First, to quantitatively evaluate spatial architecture reconstruction, we applied PCA to DTractor’s inferred cell type composition matrices across three preprocessing conditions (4,700-14,382 shared genes). PCA efficiently transforms complex multi-cell-type profiles into principal components, with PC1 capturing the predominant spatial organization pattern. DTractor, along with Cell2location and SPOTlight, consistently showed strong correlation (r > 0.6) between PC1 values and ground-truth annotations across all conditions, accurately reconstructing the MOB’s layered architecture (Fig. 4c, Supplementary Fig. S30). Next, PC1 visualization (Fig. 4d) revealed DTractor, Cell2location, and SPOTlight effectively segregating GCL from the outer layers (MCL/EPL, GL/EPL, ONL). Consistent with Ma et al. [16], who noted high correlation among MCL/EPL, GL/EPL, and ONL—strongest between MCL/EPL and GL/EPL—DTractor and SPOTlight best reflected this similarity. In contrast, DestVI and CARD resolved only two layers, while other methods exhibited either random patterns with less distinct spatial structures or excessively blurred boundaries that hindered the differentiation of each layer.

To enhance visualization clarity, we simplified cell types by consolidating subtypes into eight major categories following the CARD preprocessing approach [16] (for example, consolidating seven GC subtypes into one GC type, and similarly for PGC and M/TC) for subsequent analyses. We then evaluated each model’s ability to recapitulate known anatomical structures through three complementary analyses. First, we analyzed the dominant simplified cell types inferred at each spatial location (Supplementary Fig. S31), which showed that DTractor accurately reconstructed the MOB’s layered organization as described previously [52]. Second, to quantitatively evaluate regional differentiation, we performed pairwise two-sided Wilcoxon rank-sum tests on PC1 across four anatomically distinct regions (Fig. 4e). The resulting heatmaps display p-values for each pairwise comparison, with blue cells indicating statistically significant differences and red cells showing non-significant differences. DTractor, DestVI, and Cell2location successfully differentiated all region pairs with high statistical significance, aligning with histological annotations, whereas other methods such as Stereoscope showed limited discriminative power between certain regions (MCL/EPL vs GL/EPL). Third, we examined PC1 scores distribution using boxplots (Supplementary Fig. S32), where DTractor accurately reflected the anatomical progression from GCL to MCL/EPL to GL/EPL to ONL, with median scores maintaining correct anatomical order despite some overlap between MCL/EPL and GL/EPL due to their molecular similarity. Collectively, these analyses demonstrated that DTractor was the only model that consistently recapitulated the known anatomical organization of the MOB, while other methods showed only partial success in regional differentiation.

Next, we extended our evaluation beyond regional differentiation to assess each method’s capability for fine-grained cellular decomposition. By comparing the dominant cell types inferred at each spatial location against ground-truth annotations for 18 distinct cell types, we quantified decomposition accuracy using three complementary metrics (Fig. 4f, Supplementary Fig. S33). DTractor consistently outperformed all other methods across adjusted rand index (ARI, 0.36), purity (0.76), and normalized mutual information (NMI, 0.43) with high statistical significance (P=1.81×10^−25^, P=7.27×10^−35^, and P=2.44×10^−29^ compared to the second-best, respectively). These results demonstrate that beyond accurately capturing broad tissue regions, DTractor achieves superior resolution at the individual location level, providing more precise characterization of the complex cellular architecture within the MOB. With simplified cell types, DTractor still remained competitive (Supplementary Fig. S34), though overall *R*^2^ scores were modest likely due to the naive separation of MOB spatial structures into four major layers, as previously mentioned, with DTractor, Cell2location, and RCTD leading with the limited available information (Supplementary Fig. S35).

Finally, DTractor’s key strength lies in its robust resolution of cell type proportions deconvolutions across layers (Fig. 4g), independent of shared gene numbers—a capability unmatched by other methods. It precisely distinguished individual layers, including the highly correlated MCL/EPL and GL/EPL, correctly enriching M/TC in MCL/EPL and PGC in GL/EPL while minimizing false positives, even with only 4,700 genes, unlike the diffuse predictions of competing methods (Supplementary Figs. S36-38). Uniquely, DTractor also accurately positioned EPL-IN cell type between MCL and GL (Supplementary Fig. S31), addressing a challenge noted by Ma et al. [16]. Additionally, DTractor achieved superior memory efficiency (250 MB vs. 2,000-19,500 MB) and competitive computation times (400-500 seconds) across all gene counts tested, without compromising accuracy or robustness, surpassing Redeconve’s downsampling strategy (Supplementary Fig. S39).

### DTractor achieves superior cell count estimation in two simulated datasets with known ground truth

We benchmarked DTractor against eight established deconvolution methods using two simulated spatial transcriptomics (ST) datasets with known ground truth to assess its cell count estimation accuracy. Simulated data is essential for this evaluation as real datasets lack ground truth cell annotations, making reliable performance comparisons challenging. Total cell counts per spot, indicative of biological activity or density such as in tissue structures or pathological regions [54, 55], served as a key regional metric.

The first dataset, adapted from Cell2location [19], comprised 2,500 spatial locations derived from single-nucleus RNA sequencing (snRNA-seq) data of mouse brain sections, integrating 49 neural cell types (Supplementary Table S1). The reference snRNA-seq data, preprocessed conventionally and visualized via UMAP [56] initialized by PAGA [57] with Leiden clustering, included astrocytes, excitatory and inhibitory neurons, oligodendrocyte precursor cells, oligodendrocytes, and microglia. The simulated spatial dataset featured diverse abundance patterns—ubiquitous high and low, and spatially restricted high and low—mirroring Cell2location’s validation approach. Unlike Cell2location, we evaluated overall deconvolution performance across all 2,500 locations for a comprehensive assessment.

To evaluate DTractor’s accuracy in inferring spatial cellular composition, we first examined how well it captures the underlying spatial architecture of tissue. Principal component analysis (PCA) [58, 59] of DTractor’s estimated cell type composition matrix revealed that its first principal component (PC1) faithfully reproduced the anatomical structure observed in ground-truth spatial transcriptomics data across 2,500 locations (Fig. 5a). This dimensional reduction approach allowed us to visualize how DTractor preserves the dominant spatial patterns of cellular organization, outperforming other deconvolution methods (Supplementary Figs. S40, S41). Importantly, DTractor’s PC1 precisely mirrored the total cell count distribution per spot, providing an accurate representation of local cellular density—a critical feature for understanding tissue architecture. Next, we quantified DTractor’s performance in estimating absolute cell type counts at each spatial location (Fig. 5b). To assess reproducibility, we generated three distinct spatial transcriptomics datasets (each with 2,500 locations) using different random seeds. DTractor consistently achieved higher accuracy than competing methods across all datasets (Pearson improvement from 0.4 to 0.52; P=3.02×10^−29^, Spearman improvement from 0.42 to 0.57; P=1.66×10^−25^), demonstrating robust performance regardless of initialization conditions. Multiple bootstrapping experiments further validated these findings (Supplementary Fig. S42). Additionally, cosine similarity analysis confirmed DTractor’s strong directional alignment with ground truth values, with most methods showing directional relationships, while Tangram and Redeconve performed poorly in this metric (Supplementary Fig. S43).

**Fig. 5.**
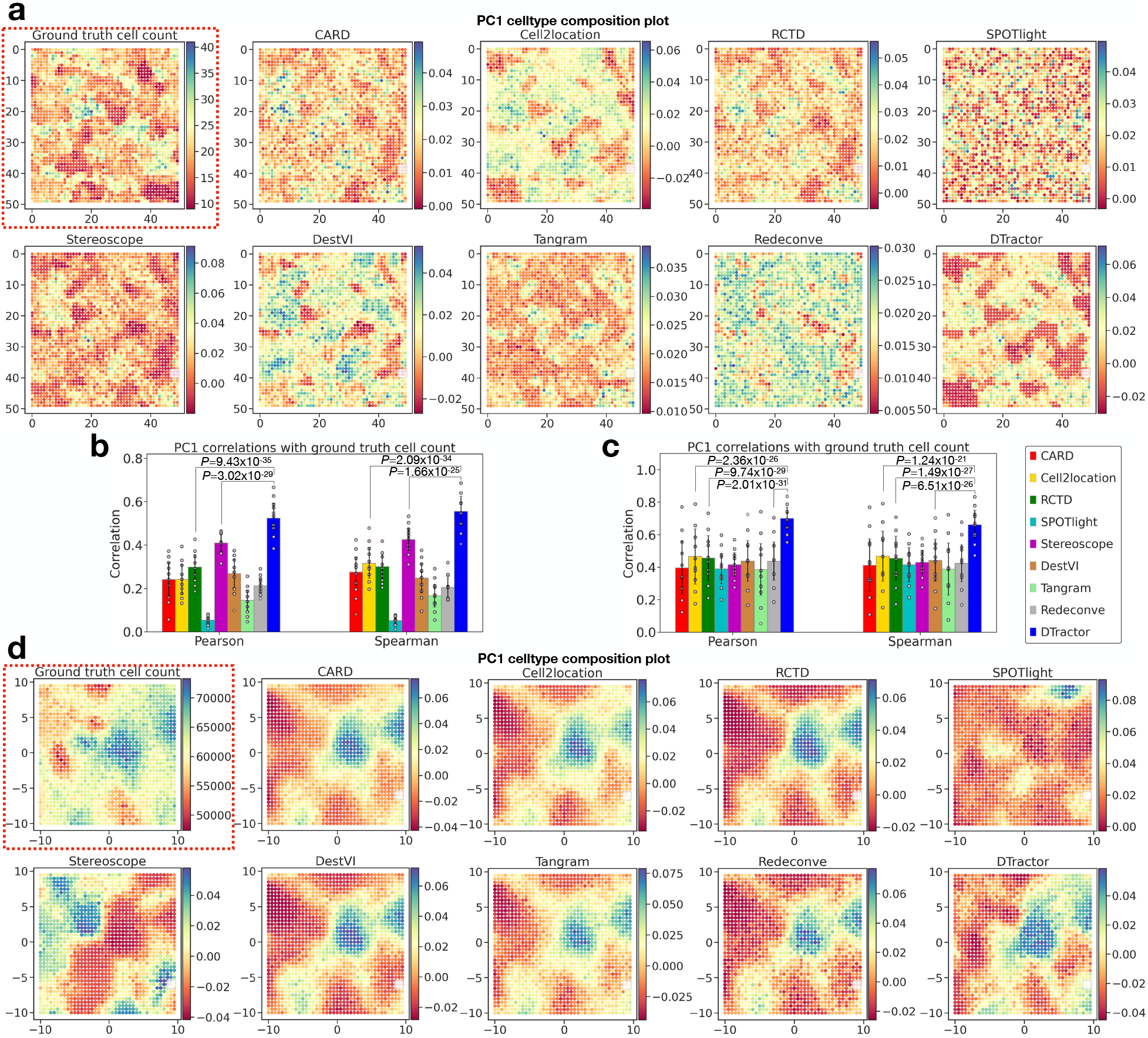
DTractor also exhibits superior accuracy in deconvoluting cell counts across two distinct simulated datasets. **a**, Top left (red dotted box): Ground truth total cell counts for all 49 cell types across all 2500 locations (single seed). The remaining plots: The first principal component (PC1) of the estimated cell type composition matrix (single seed). **b, c**, Correlations between the PC1 of the estimated cell type composition matrix and the ground truth total cell counts (with **b** corresponding to **a** and **c** corresponding to **d**), depicted through bar plots showing Pearson and Spearman correlation coefficients among nine methods. P-values are derived from one-sided Mann-Whitney U tests without adjustment. The plots were generated using 100 bootstrap resamples. The height of each bar represents the average statistics from 100 resampled bootstraps across randomly generated datasets with three different seeds while each scatter dot represents a different resample and the error bars indicate the standard errors. **d**, Top left (red dotted box): Ground truth total cell counts for all 5 cell types across all 1600 locations (single seed). Each spot is scaled to 0.5. The remaining plots: The PC1 of the estimated cell type composition matrix (single seed). **a, d**, The total number of cells in a spot can hold biological significance, as spots with notably high or low counts often correspond to key regions of interest. We compared nine methods, including DTractor, to assess the quality of their cell type proportion outputs and determine their effectiveness in capturing these regions.

The second dataset, adapted from DestVI [18], comprised 1,600 spatial locations from murine lymph node scRNA-seq data (32,000 cells, five cell types), with each spot scaled to 0.5 in a 1×1 grid. Following standard preprocessing and visualization, we evaluated inferred cell counts within each spot. Quantitative bar plots showed DTractor’s superior accuracy in absolute cell count estimation across three seeded ST datasets (Pearson improvement from 0.46 to 0.70; P=2.36×10^−26^; Spearman improvement 0.47 to 0.66; P=1.24×10^−21^), validated by bootstrapping (Fig. 5c, Supplementary Fig. S44). Subsequently, DTractor’s PC1 closely matched the ground-truth anatomical structure, uniquely capturing subtle details—for example, moderate-to-high cell type counts near (−5, −10) and (10, −10), and minimal counts at (−7, −1), (−2, 4), and (−2, 10)—where SPOTlight and Stereoscope faltered, and other methods missed these nuances (Fig. 5d, Supplementary Figs. S45, S46). This demonstrates DTractor’s superior accuracy in inferring cell counts with spots.

Finally, DTractor demonstrated remarkable computational efficiency across both simulations. It consistently maintained the lowest peak memory usage (under 400 MB) compared to other methods (1,000-8,500 MB) while achieving competitive computation times, without sacrificing accuracy or robustness (Supplementary Figs. S47, S48). In summary, DTractor excelled in anatomical structure reconstruction (Fig. 5a, d), cell count estimation (Fig. 5b, c), and robustness, underscoring its precision with known ground truth. The results suggest that DTractor provides advantages beyond cell type deconvolution for each spot, demonstrating improved accuracy in estimating cell counts at the spot level.

## Discussion

We present DTractor, a deep learning–based framework for accurate and biologically coherent deconvolution of spatial transcriptomics data. By integrating reference single-cell RNA sequencing data with spatial transcriptomic profiles, DTractor enables the estimation of both cell-type proportions and cell-type-resolved gene expression across tissue sections. Through a unified modeling framework that combines deep generative learning, transfer learning, and spatial regularization, DTractor consistently outperforms existing methods across a range of simulated and real datasets, including human lymph node, pancreatic cancer, and mouse olfactory bulb.

DTractor introduces several key innovations that distinguish it from existing spatial deconvolution frameworks in both methodological design and biological utility. First, a central contribution is its dual utilization of single-cell RNA sequencing reference data. Rather than relying solely on cell-type-specific average expression, DTractor jointly integrates two complementary components: a gene expression matrix serving as transcriptional anchors for each cell type, and latent cell-type embeddings learned through a deep variational autoencoder. These embeddings are aligned with those inferred from spatial transcriptomics data via cross-modal regularization, ensuring consistent representation of cell types across modalities and capturing both observed and latent transcriptional structures for biologically coherent decomposition. Second, DTractor employs a hybrid architecture that combines deep generative modeling with structured matrix factorization, enabling flexible low-dimensional embedding learning while retaining the interpretability of linear decomposition. This design brings together the expressive power of deep learning and the clarity of classical approaches, supporting accurate and explainable spot-level predictions. Third, DTractor demonstrates strong robustness across diverse spatial technologies, tissue types, spot resolutions, and reference qualities. It effectively resolves both fine-grained subtypes and distinct cell types, including those in complex immune or tumor microenvironments, without relying on marker genes or manual feature curation, addressing a major limitation of many current tools. Fourth, DTractor incorporates adaptive spatial regularization that encourages local coherence while preserving sharp region boundaries. Unlike models that ignore spatial context or impose rigid priors, DTractor learns smoothness constraints during training, enhancing spatial fidelity in both homogeneous and heterogeneous tissues. Fifth, DTractor uniquely estimates both cell-type proportions and total cell counts per spatial location—an uncommon capability that enables quantitative reconstruction of cellular density and tissue organization. This facilitates insights into processes such as immune infiltration, developmental gradients, and tumor growth. Finally, DTractor achieves strong performance while maintaining exceptional memory efficiency, owing to its staged training pipeline, compact latent spaces, fixed reference matrix, and lightweight matrix factorization–based optimization. These design choices eliminate the need for large computational graphs, dense spatial kernels, or per-spot neural decoders, resulting in consistently low memory usage (usually less than 1GB) even on large-scale spatial datasets. Together, these advances result in a unified and interpretable framework that improves spatial decomposition fidelity, supports diverse biological analyses, and generalizes across experimental modalities and tissue systems.

A potential limitation of DTractor lies in its reliance on the assumption that the single-cell RNA sequencing reference and the spatial transcriptomics data contain a similar set of cell types. The model performs decomposition based on both the gene expression profiles and the latent representations learned from the reference. Therefore, its accuracy depends on the presence and alignment of comparable cell states in both modalities. If important cell types are absent or underrepresented in the reference, due to sampling bias or technical variability, the decomposition may misassign proportions or fail to detect cell populations that are present only in the spatial data. To address this limitation, future extensions could incorporate mechanisms to detect previously unobserved cell types by including residual expression components or introducing latent factors that are not anchored to the reference. Additionally, adaptive training strategies that allow partial supervision or flexible alignment in ambiguous regions may help DTractor remain robust in the presence of incomplete or non-overlapping references.

DTractor offers a robust, scalable, and accurate solution for spatial spot decomposition, addressing key limitations of existing methods. Its ability to generalize across tissue types and platforms ensures reliable performance in diverse biological settings, while its computational efficiency supports large-scale analysis without sacrificing precision. By accurately resolving both cell-type proportions and total cell counts, DTractor enables detailed, biologically coherent reconstructions of tissue organization. These capabilities position DTractor as a versatile tool for spatial transcriptomics, facilitating deeper insights into cellular architecture across development, disease, and regeneration. DTractor is the only existing framework that combines dual-reference integration, adaptive spatial regularization, total cell count estimation, and memory-efficient scalability, offering a uniquely comprehensive and interpretable solution for spatial transcriptomic deconvolution.

## Methods

### Data Preprocessing

DTractor requires single-cell RNA-seq (scRNA-seq) data annotated with cell types and spatial transcriptomics (ST) data with spatial coordinates. To prepare these datasets for integration, we performed standardized preprocessing steps to remove technical artifacts, normalize expression values, and align features across modalities. For scRNA-seq data, we used `AnnData` objects from the `anndata` Python package [60] to store gene expression matrices and metadata. Mitochondrial genes were removed to reduce technical noise [19]. Using `Scanpy` [61], we filtered out low-quality cells and low-abundance genes, applied library size normalization to account for sequencing depth, and log-transformed the data to stabilize variance. All preprocessing parameters were adopted from the original publications associated with each dataset. If a dataset was already preprocessed, we skipped the filtering and normalization steps. To correct for potential batch effects, we used `scGen` [62], a deep generative model that aligns cells across batches while preserving biological variation. Only genes shared between the scRNA-seq and ST datasets were retained for integration, and the model was trained using default parameters. Spatial transcriptomics data were preprocessed using similar procedures. We constructed `AnnData` objects containing expression matrices and spatial coordinates. Low-quality spots and rarely detected genes were filtered using `Scanpy`, followed by normalization and log-transformation. Only genes shared with the scRNA-seq dataset were retained for integration. This pre-processing yielded a single-cell expression matrix *X* ∈ ℝ^*C*×*n*^ and a spatial transcriptomics matrix *Y* ∈ ℝ^*S*×*n*^, where *C* and *S* denote the number of cells and spatial spots, respectively, and *n* is the number of shared genes between the two datasets.

### Learning Latent Representations for scRNA-seq and Spatial Transcriptomics Data

To obtain compact and informative representations of the reference single-cell (SC) and target spatial transcriptomics (ST) data, **DTractor** employs two separate variational autoencoders (VAEs) to learn latent embeddings of cells and spatial spots, respectively (Fig. 1a,b). VAEs are probabilistic generative models that map high-dimensional inputs into a lower-dimensional latent space via an encoder and reconstruct the input through a decoder, effectively capturing the underlying data manifold [63]. This approach is well-suited for modeling sparse and noisy single-cell and spatial transcriptomics data. Each VAE uses a zero-inflated negative binomial (ZINB) likelihood, which captures both overdispersion and the abundance of zeros caused by technical dropout and biological non-expression [64, 65]. The ZINB combines a negative binomial component for modeling gene count variability with a zero-inflation term for dropout.

For the single-cell dataset **X** ∈ ℝ^*C*×*n*^, the encoder 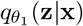 maps each cell *x*_*i*_ ∈ ℝ^*n*^ to a latent distribution 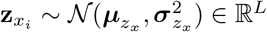, where *L* ≪ *n*, and *θ*_1_ are the encoder parameters. The decoder 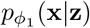 reconstructs gene expression from the latent representation, outputting three parameters of the ZINB distribution: means ***µ***, dispersions ***θ***, and dropout probabilities ***π***. Similarly, for spatial transcriptomics data **Y** ∈ ℝ^*S*×*n*^, the encoder 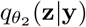and decoder 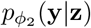 define the VAE for spatial spots, generating latent embeddings 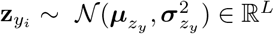. The overall loss function for the VAE with ZINB likelihood is:

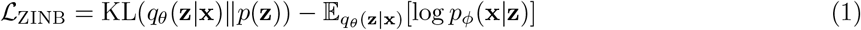

where the KL divergence term regularizes the approximate posterior *q*_*θ*_(**z** | **x**) toward the prior *p*(**z**) ~ 𝒩 (**0, I**), and the reconstruction term encourages accurate modeling of the observed data [66]. The ZINB log-likelihood for a reconstructed gene expression vector **x** is:

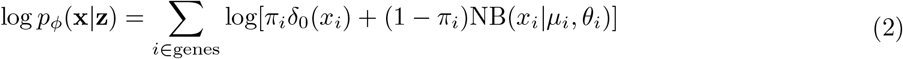

Here, *π*_*i*_ is the dropout (zero-inflation) probability for gene *i*, and *µ*_*i*_, *θ*_*i*_ are the gene-specific mean and dispersion parameters of the negative binomial component, both generated by the decoder network from the latent embedding **z**. In contrast, *π*_*i*_ is modeled as a free gene-specific parameter, independent of **z**. We implemented the models using the `scvi-tools` Python package [67], which supports ZINB-based VAEs with GPU-accelerated training. Any negative expression values introduced by upstream batch correction were clipped to zero before training, as required by the ZINB likelihood. For SC data, batch effects can alternatively be handled directly in the VAE using SCVI [65].

To ensure comparability between modalities, we fixed the latent dimensionality as Dim_SC_ = Dim_ST_ = *L*. Both VAEs were trained in an unsupervised manner without using cell type or region labels. For `VAE`_SC_, we selected the model epoch with the highest adjusted Rand index (ARI) between latent clusters and annotated cell types. For `VAE`_ST_, we applied the elbow method [68] on the reconstruction loss to determine the optimal stopping point. Final latent embeddings are denoted 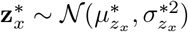 and 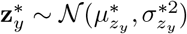. The DTractor framework is flexible and supports alternative generative models; users may substitute the VAE with other architectures as needed. For benchmarking, we optionally evaluated ARI, normalized mutual information (NMI), and average silhouette width (ASW) using cell-type labels.

Both encoder and decoder networks consist of linear layers with ReLU activations [69]. Hidden layer widths were set to 128 or 256 depending on the dataset. Latent dimensions were set to 16 or 32 by default. Training used mini-batches, early stopping, dropout, and a maximum of 50 epochs for `VAE`_SC_ and 70 for `VAE`_ST_. These default settings provided strong and consistent performance without extensive hyperparameter tuning. Architectural choices were adapted from SCVI to ensure stable and efficient model training.

### ST decomposition framework for spatial mapping of cell types

#### Model formulation

DTractor reconstructs spatial gene expression by combining spatial embeddings learned via a variational autoencoder (VAE) with cell-type-specific gene expression profiles from single-cell reference data. Let **Y** ∈ ℝ^*S*×*n*^ denote the spatial transcriptomics (ST) gene expression matrix with *S* spots and *n* genes, 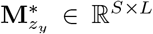 the latent VAE embeddings, **T** ∈ ℝ^*L* × *k*^ the trainable cell-type embedding matrix, and **B** ∈ ℝ^*k*×*n*^ the single-cell reference profiles for *k* cell types. We first compute:

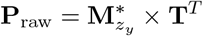

as an unconstrained score matrix indicating the presence of each cell type across spatial spots. However, because we interpret these values as cell-type proportions, they must be non-negative. To enforce this, we apply small-base exponentiation, with the base parameter set to 1.001 as the default value followed by normalization:

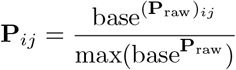

This transformation ensures that all entries in **P** ∈ ℝ^*S*×*k*^ are non-negative, allowing them to be interpreted as cell-type proportions. Additionally, the small-base exponentiation preserves smooth gradients for optimization and helps control the numerical scale of the values, reducing the risk of unstable updates during training. The reconstructed ST matrix is then:

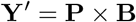

and the model is trained to minimize the Frobenius norm between the observed and reconstructed gene expression:

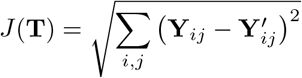

#### Training

We optimize **T**^*T*^ using the Adam optimizer [70] implemented in PyTorch [71], with default hyperparameters. Detailed iteration strategies and tuning recommendations are provided in the GitHub repository. This formulation enables DTractor to infer spatially resolved, non-negative, and biologically interpretable cell-type proportions in complex tissues.

### Matrix Factorization Regularization

As part of DTractor’s matrix factorization framework, we introduce two optional regularizations designed to enhance spatial transcriptomics (ST) decomposition by incorporating spatial smoothness and single-cell (SC) embedding alignment. These regularizations act on the factorized matrices—the cell type proportion matrix *P* and the cell type embedding matrix *T* —and can be flexibly enabled or disabled depending on dataset complexity and biological context.

#### Spatial proximity regularization

This regularization is applied directly to the cell type proportion matrix *P* ∈ ℝ^*S*×*k*^, where *S* is the number of spatial spots and *k* is the number of cell types. It is based on the biological observation that spatially adjacent tissue regions tend to exhibit similar cellular compositions [16, 72, 73]. First, we construct a fixed *K*-nearest neighbors (KNN) graph for the spatial spots based on Euclidean distances between their coordinates (default *K* = 10). Then, we compute a pairwise Euclidean distance matrix *D*_*P*_ ∈ ℝ^*S*×*S*^ between all rows *p*_*i*_ and *p*_*j*_ of *P* :

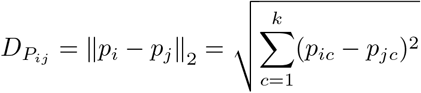

To obtain differentiable soft rankings, we normalize each row:

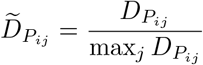

and apply a softmax using an exponent base of 1024:

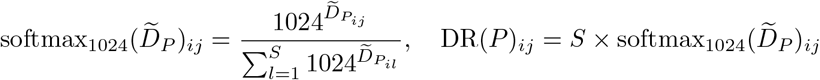

This yields a differentiable proxy for ranking, where DR(*P*)_*ij*_ approximates the soft rank of spot *j* relative to spot *i*. The choice of base 1024 empirically improved rank separation in our model and mimics the effect of a low-temperature softmax, amplifying differences in otherwise close distances while maintaining stability due to max normalization. We then define the spatial regularization loss as:

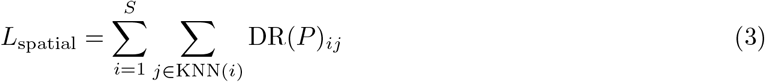

This loss encourages nearby spots to exhibit similar cell type proportions, enforcing local smoothness. Unlike fixed spatial priors in other methods, DTractor allows this regularization to be weighted or scheduled during training to better accommodate heterogeneous datasets. (See Supplementary Fig. S49 for an illustrative example.)

#### SC-ST cell type embedding consistency regularization

This regularization constrains the cell type embedding matrix *T* ∈ ℝ^*k*×*L*^ to match the relational structure of cell types derived from the SC reference data. We construct a reference matrix *E* ∈ ℝ ^*k*×*L*^ by averaging the latent embeddings 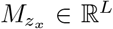 of all SC cells of each type:

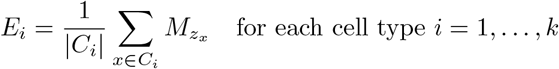

We compute pairwise Euclidean distances between the embeddings:

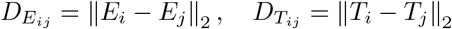

These matrices are normalized and transformed into differentiable rank approximations using the same 1024-based softmax:

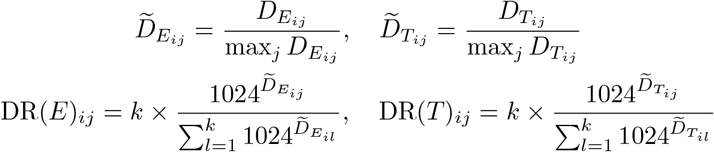

The regularization loss is computed using the mean squared error between the SC and ST distance rank matrices:

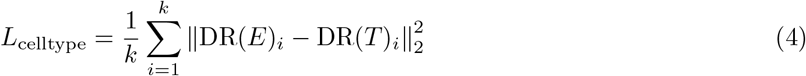

This loss encourages the geometry of cell type embeddings in ST to reflect the biological structure observed in SC data, promoting biologically grounded decomposition.

#### Trade-offs and recommendation

Both regularizations enhance interpretability and robustness but come with additional computational cost—particularly in datasets with large numbers of spots and cell types. Further, some cell types may benefit more than others depending on spatial structure or alignment with SC references. We therefore recommend using the default Frobenius loss alone unless preliminary data analysis or pilot runs indicate that spatial or embedding regularization would substantially improve performance.

### Inference of Cell Type Proportion Matrix

As training proceeds, the matrix *T* ^T^ is iteratively updated to better map latent features to biological cell types. Cell types that consistently contribute to the reconstruction of specific spatial spots gradually receive higher weights in the proportion matrix *P* ∈ ℝ^*S*×*k*^, while irrelevant types are downweighted. This process ensures that dominant cell types in each region are accurately captured over time. At convergence, the final cell type proportion matrix *P* is row-normalized so that the proportions across all *k* cell types sum to 1 for each spot. This normalization step preserves a probabilistic interpretation of cell type composition across the tissue. While increasing the embedding dimension allows for more expressive modeling of latent cell type signals, it also introduces additional computational complexity. Thus, a balance between flexibility and efficiency must be considered.

#### Final loss function

To guide accurate decomposition, we incorporate additional regularization terms into the final objective:

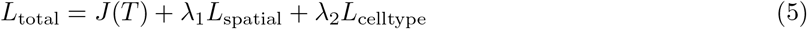

where *J* (*T*) is the reconstruction loss, *L*_spatial_ enforces local spatial coherence in the proportion matrix *P*, and *L*_celltype_ encourages consistency between the cell type embeddings derived from ST and SC data. The model is trained using an iterative matrix decomposition strategy outlined in Algorithm 1.

##### Algorithm 1

DTractor: Iterative Decomposition and Regularized Optimization

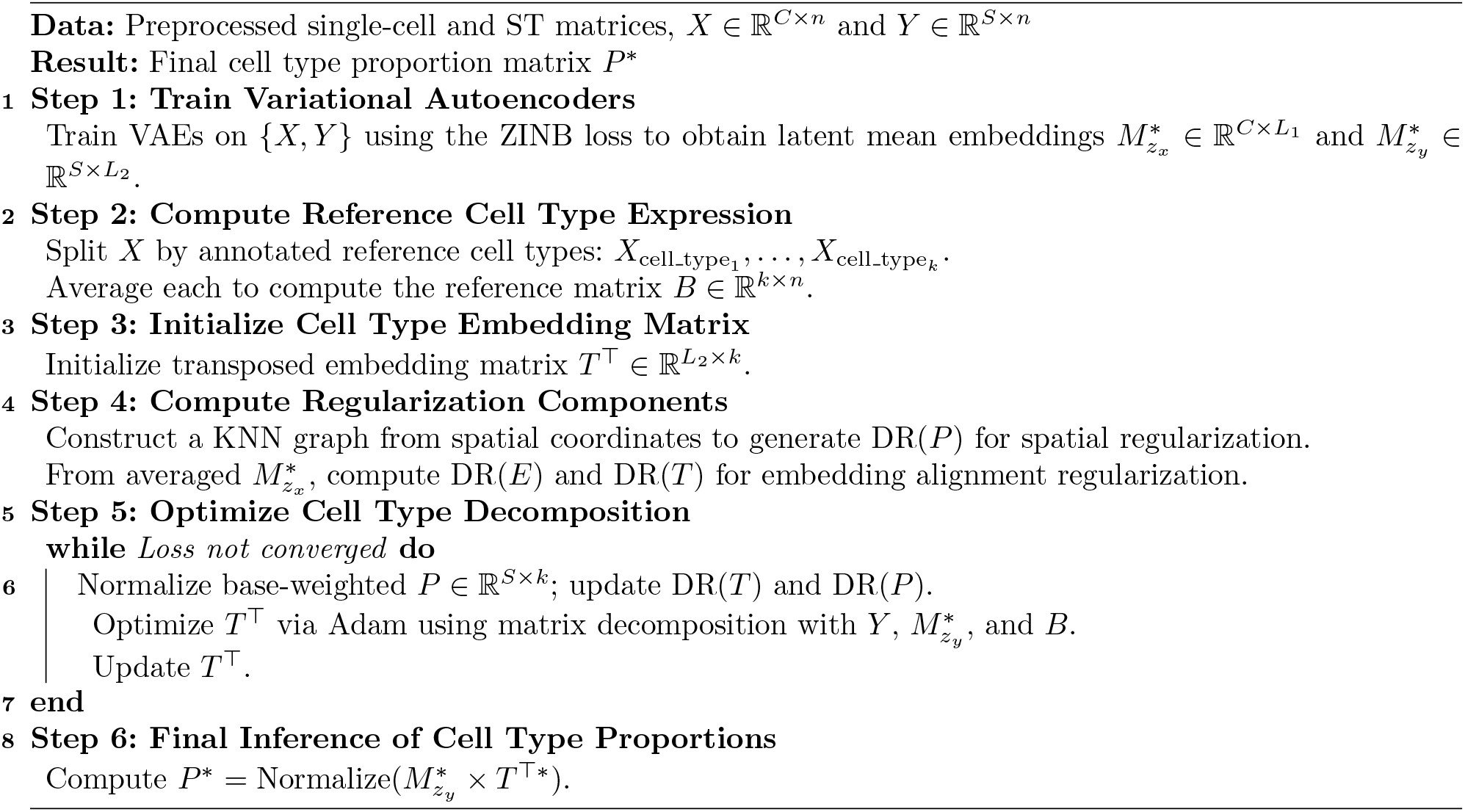

### Performance validation for benchmark evaluations

To evaluate DTractor’s performance in inferring cell type proportions from the integration of single-cell reference and spatial transcriptomics (ST) data, we benchmarked it against eight widely used deconvolution and mapping methods: CARD [16], Cell2location [19], RCTD [14], SPOTlight [17], Stereoscope [15], DestVI [18], Tangram [21], and Redeconve [20]. These methods are all peer-reviewed, publicly available, and explicitly designed to estimate cell or cell type abundances in spatial locations using both ST data and single-cell RNA-seq references.

Each tool represents a distinct modeling philosophy. CARD applies a conditional autoregressive model on top of non-negative matrix factorization to incorporate spatial smoothness. Cell2location uses hierarchical Bayesian modeling with negative binomial likelihood to estimate cell type abundance across spots. RCTD applies Poisson probabilistic modeling and supervised decomposition based on reference profiles. SPOTlight combines seeded non-negative matrix factorization with non-negative least squares regression, incorporating marker gene selection. Stereoscope jointly models single-cell and spatial expression counts using a negative binomial framework. DestVI integrates latent variable models trained on single-cell and spatial data using variational inference. Tangram leverages deep learning and non-convex optimization to spatially align single-cell data. Redeconve addresses collinearity in non-negative least squares regression with regularization, assuming similar single-cell states yield similar spatial abundances.

All methods were evaluated using identical single-cell and spatial transcriptomics inputs and consistent preprocessing. We focused on three main criteria: (1) the accuracy of cell type proportion estimation compared to known tissue annotations, (2) the usability and training stability of the methods, and (3) computational efficiency. To reduce bias and ensure reproducibility, we followed default parameter settings recommended in each tool’s documentation or GitHub repository, only modifying parameters when necessary to ensure convergence or compatibility. For example, we adjusted filtering thresholds in CARD and RCTD to prevent excessive gene or cell exclusion, increased training epochs in Cell2location, Stereoscope, and DestVI based on ELBO or loss convergence, and used cell-level mapping in Tangram for finer resolution despite increased memory demand. DTractor was run with 250,000 training iterations, and the default regularization weights (*λ*_1_ = *λ*_2_ = 0) were maintained for benchmarking unless otherwise noted.

To assess performance quantitatively, we applied principal component analysis (PCA) on inferred proportion matrices to identify dominant axes of variation, and visually inspected these using UMAP projections initialized with PAGA, implemented via Scanpy [61]. We then compared the principal components to ground truth tissue region annotations using Pearson and Spearman correlations, as well as cosine similarity scores. To quantify how well inferred clusters recapitulated annotated tissue zones, we computed adjusted Rand index (ARI), normalized mutual information (NMI), and purity scores. Logistic regression models were trained on the PCA components of inferred proportions to predict tissue regions, using Nagelkerke *R*^2^ for binary classification and McFadden *R*^2^ for multinomial classification, following standard implementation in Statsmodels [74]. To evaluate whether individual cell types were predictive of tissue architecture, we performed additional logistic regression analysis using the dominant cell type at each spot as input.

For region-specific evaluation, we identified key cell types based on prior literature and assessed the spatial accuracy of their distributions using correlation metrics against tissue labels. Differential enrichment across zones was tested using pairwise Wilcoxon rank-sum tests with Bonferroni correction. To assess performance in class-imbalanced settings, we computed precision-recall area under the curve (AUC-PR) scores for each cell type’s ability to identify its associated spatial region. We also visualized spatial patterns using Cell2location’s `plot_spatial` utility and overlaid cell abundances on histological images where available. Finally, we examined co-localization and enrichment patterns by computing pairwise correlations across inferred proportions, highlighting functional tissue organization and microenvironmental structure.

We tested all methods across five benchmark datasets: two simulated and three real. The first simulated dataset was generated following the Cell2location benchmarking protocol, using 2,500 spatial locations and gene expression profiles from 49 cell types derived from mouse brain snRNA-seq data [19]. In this design, 8 cell types were spatially ubiquitous, and 41 were region-specific across 12 tissue zones. Cell types were assigned to 1–3 zones, each containing 2–8 cell types, with abundance profiles stratified by low or high levels to reflect empirical sparsity. The second synthetic dataset followed the DestVI simulation framework [18], consisting of a 40×40 grid of 1,600 spatial spots. Cell type proportions at each spot were generated from a spatial Gaussian process with a covariance kernel ( = 0.1), while expression counts were simulated using a negative binomial distribution based on real murine lymph node scRNA-seq data. Five major immune cell types were retained: B cells, CD4 T cells, CD8 T cells, migratory dendritic cells, and Tregs.

Real tissue datasets included a human lymph node dataset compiled by Kleshchevnikov et al. [19] from three scRNA-seq studies [22–24], with 73,260 cells across 34 cell types and matching 10x Visium ST data (4,035 spots). Batch correction was performed using scGen [62]. We also evaluated a human pancreatic ductal adenocarcinoma (PDAC) dataset [25], where ST data (428 spots) and matched inDrop scRNA-seq data (1,926 cells, 20 cell types) were derived from the same patient. The final dataset was from the mouse olfactory bulb, where ST was performed using the Spatial Transcriptomics protocol [11] on 282 locations, and matched scRNA-seq data (21,746 cells across 18 cell types) was obtained from 10x Chromium [26]. Manual annotations of tissue zones were assigned to each spot by overlaying H&E-stained images and grouping them into GCL, MCL/EPL, GL/EPL, and ONL. All details of spots and cells are provided in Supplementary Tables S1-S3.

To assess the robustness of each method to data preprocessing, we repeated all analyses using three alternative gene filtering schemes per dataset to define the shared gene set between SC and ST data. Many methods depend heavily on gene filtering and shared gene selection, yet robustness to these parameters is rarely assessed systematically. While some tools such as CARD, DestVI, Redeconve, and SPOTlight have evaluated certain robustness aspects, none have comprehensively addressed the influence of preprocessing variability. To fill this gap, we applied each method on three gene filtering configurations and measured the change in performance across all metrics. Since the sample size (n = 3) was insufficient for statistical testing, we applied bootstrapping with 10, 50, 100, and 500 iterations to estimate empirical distributions of performance differences. This enabled rigorous evaluation of whether each method maintained stable accuracy across varying preprocessing conditions—a critical factor for reliable deployment of deconvolution tools in diverse datasets.

For benchmarking running time and memory requirements, we performed all experiments on a high-performance workstation equipped with three NVIDIA GeForce RTX 4090 GPUs and an AMD Ryzen Threadripper PRO 5965WX 24-core processor (48 threads) with a x86 64 architecture. The system was equipped with 512 GB of system memory (RAM).

## Data availability

Our study utilized publicly available datasets. The processed mouse brain snRNA-seq data for the first simulation is available on the ArrayExpress website under accession number E-MTAB-11115 (https://www.ebi.ac.uk/biostudies/arrayexpress/studies/E-MTAB-11115) [19]. Annotated snRNA-seq data for cell types are publicly accessible via the cellxgene portals (https://zenodo.org/records/3710410). The murine lymph node data for the second simulation has raw data deposited in the National Center for Biotechnology Information (NCBI) Gene Expression Omnibus (GEO) repository with accession number GSE173778 and processed data available in the DestVI [18] repository (https://github.com/romain-lopez/DestVI-reproducibility/tree/master/lymph_node). The human lymph nodes Visium data can be accessed at 10X Genomics (https://www.10xgenomics.com/datasets/human-lymph-node-1-standard-1-1-0), and the processed scRNA-seq is available at the Cell2location S3bucket (https://cell2location.cog.sanger.ac.uk/browser.html). The processed Visium spatial expression data can be easily downloaded using the scanpy [61] function ‘scanpy.datasets.visium_sge’ with the sample id ‘V1_Human_Lymph_Node.’ Ground truth annotations of GC zones in Visium data can be found on the Cell2location GitHub page (https://github.com/vitkl/cell2location_paper/tree/master/notebooks) or for both GC and T-cell zones on the Redeconve [20] Code Ocean page (https://codeocean.com/capsule/1351962/tree/v1). The ST and scRNA-seq data used for PDAC is publicly available in the NCBI GEO database under accession number GSE111672 from Moncada et al. [25]. Ground truth annotations of four tissue regions can be found on either the CARD [16] GitHub page (https://github.com/YMa-lab/CARD/tree/master/data) or the Redeconve Code Ocean page (https://codeocean.com/capsule/1351962/tree/v1). The MOB ST data is available at the Spatial Research lab (https://www.spatialresearch.org/resources-published-datasets/doi-10-1126science-aaf2403/), and the scRNA-seq data is available in the NCBI GEO repository under accession number GSE121891.

## Code availability

The DTractor source code and results reproduction are publicly available at our GitHub repository (https://github.com/mcgilldinglab/DTractor).

## Acknowledgements

This study was funded in part by grants awarded to J.D. We gratefully acknowledge the support from the Canadian Institutes of Health Research (CIHR) under Grant Nos. PJT-180505; the Funds de recherche du Québec – Santé (FRQS) under Grant Nos. 295298 and 295299; the Natural Sciences and Engineering Research Council of Canada (NSERC) under Grant No. RGPIN2022-04399; and the Meakins-Christie Chair in Respiratory Research. We also extend our gratitude to Janusz Rak, Gregory Fonseca and Yasser Riazalhossini, the academic advisor and thesis committee members of Y.J.K., for their feedback on the DTractor model.

## Author Information

J.D. conceived and designed the study. Y.J.K. and J.D. both designed the methodology. Y.J.K. collected the data, implemented the methodology, and analyzed the results. C.L. improved the code efficiency. J.D. supervised the production of results. Y.J.K. and J.D. contributed to writing and revising the manuscript. All authors have read and approved the final manuscript.

## Declarations

The authors declare no competing interests in conducting this study.

## Declaration of generative AI and AI-assisted technologies in the writing process

During the preparation of this work the author(s) used ChatGPT in order to enhance language, clarity and readability. After using this tool/service, the author(s) reviewed and edited the content as needed and take(s) full responsibility for the content of the publication.

