## Supplementary information (figures and tables) for "DTractor enhances cell type deconvolution in spatial transcriptomics by integrating deep neural networks, transfer learning, and matrix factorization"

### Contents

List of Supplementary Figures 3

List of Supplementary Tables 5

Supplementary Figures 6

Supplementary Tables 62

#### List of Supplementary Figures


**Figure S1** Bootstrap resampling to assess regional segregation in the human lymph node.
**Figure S2** Spatial mapping accuracy for six germinal center (GC) cell types in the human lymph node.
**Figure S3** Bootstrap resampling to assess spatial mapping accuracy of germinal center (GC) subtypes in the human lymph node.
**Figure S4** Bootstrap resampling to assess macro-average correlation coefficients of germinal center (GC) subtypes in the human lymph node.
**Figure S5** Estimated cell abundance of selected six known germinal center (GC) cell types along with Naive B and pre-GC B cell types in the human lymph node.
**Figure S6** Bootstrap resampling to assess spatial mapping accuracy of T-cell subtypes in the human lymph node.
**Figure S7** Estimated cell abundance of selected T cell types in the human lymph node.
**Figure S8** Spatial mapping accuracy for eight T cell types excluding the T TIM3+ subtype in the human lymph node.
**Figure S9** Bootstrap resampling to assess spatial mapping accuracy of T-cell subtypes excluding the T TIM3+ subtype in the human lymph node.
**Figure S10** Macro average correlation coefficients for subtypes in the T-cell zones of the human lymph node.
**Figure S11** Bootstrap resampling to assess macro-average correlation coefficients of T-cell subtypes in the human lymph node.
**Figure S12** Bootstrap resampling to assess macro-average correlation coefficients of T-cell subtypes excluding the T TIM3+ subtype in the human lymph node.
**Figure S13** Spatial mapping accuracy for nine T cell types in the human lymph node.
**Figure S14** Correlations in the estimated cell type composition matrix across spatial locations between pairs of cell types in the human lymph node.
**Figure S15** Computation time and peak memory usage for the human lymph node.
**Figure S16** Bootstrap resampling to assess regional segregation in PDAC.
**Figure S17** Boxplot and heatmap of pairwise two-sided Wilcoxon-rank sum test between the four tissue regions (13402 shared genes) in PDAC.
**Figure S18** Boxplot and heatmap of pairwise two-sided Wilcoxon-rank sum test between the four tissue regions (9818 shared genes) in PDAC.
**Figure S19** Boxplot between the four tissue regions (8729 shared genes) in PDAC.
**Figure S20** Inferred dominant cell type (13402 shared genes) in PDAC.
**Figure S21** Inferred dominant cell type (9818 shared genes) in PDAC.
**Figure S22** Inferred dominant cell type (8729 shared genes) in PDAC.
**Figure S23** Bootstrap resampling to assess spatial mapping accuracy of cancer subtypes in PDAC.
**Figure S24** Bootstrap resampling to assess macro-average correlation coefficients of cancer subtypes in PDAC.
**Figure S25** Estimated fibroblasts cell abundance (13402 shared genes) in PDAC.
**Figure S26** Estimated fibroblasts cell abundance (9818 shared genes) in PDAC.
**Figure S27** Estimated fibroblasts cell abundance (8729 shared genes) in PDAC.
**Figure S28** Correlations in the estimated cell type composition matrix across spatial locations between pairs of cell types in PDAC.
**Figure S29** Computation time and peak memory usage for PDAC.
**Figure S30** Bootstrap resampling to assess first principal component (PC1) correlation coefficients of entire tissue regions in the mouse olfactory bulb.
**Figure S31** Inferred dominant simplified cell type in the mouse olfactory bulb.
**Figure S32** Boxplot between the four tissue regions in the mouse olfactory bulb.
**Figure S33** Bootstrap resampling to assess similarity metrics in the mouse olfactory bulb.
**Figure S34** Bootstrap resampling to assess similarity metrics with the simplified cell types in the mouse olfactory bulb.
**Figure S35** Bootstrap resampling to assess regional segregation with the simplified cell types in the mouse olfactory bulb.
**Figure S36** The proportion from each of the four cell types of the mouse olfactory bulb.
**Figure S37** Estimated cell abundance of major cell types from the key tissue zones of the mouse olfactory bulb, from each of the GC, M/TC, and PGC subtypes.

**Figure S38** Estimated cell abundance of major cell types from the key tissue zones of the mouse olfactory bulb, from each of the GC, M/TC, and OSNs subtypes.
**Figure S39** Computation time and peak memory usage for the mouse olfactory bulb.
**Figure S40** The ground truth total cell counts and the first principal component (PC1) of the estimated cell type composition matrix for the seed v.1 in the mouse brain simulation.
**Figure S41** The ground truth total cell counts and the first principal component (PC1) of the estimated cell type composition matrix for the other seed v.2 in the mouse brain simulation.
**Figure S42** Bootstrap resampling to assess the correlation between the first principal component (PC1) and the ground truth cell count in the mouse brain simulation.
**Figure S43** Bootstrap resampling to assess the cosine similarities between the simulated cell counts and the estimated cell type composition at individual locations in the mouse brain simulation.
**Figure S44** Bootstrap resampling to assess the correlation between the first principal component (PC1) and the ground truth cell count in the murine lymph node simulation.
**Figure S45** The ground truth total cell counts and the first principal component (PC1) of the estimated cell type composition matrix for the seed v.1 in the murine lymph node simulation.
**Figure S46** The ground truth total cell counts and the first principal component (PC1) of the estimated cell type composition matrix for the other seed v.2 in the murine lymph node simulation.
**Figure S47** Computation time and peak memory usage for the mouse brain simulation.
**Figure S48** Computation time and peak memory usage for the murine lymph node simulation.
**Figure S49** Examples of two optional regularizations in ST decomposition.

#### List of Supplementary Tables

**Table S1** Information on cell types in sc/snRNA-seq datasets.
**Table S2** List of spatial transcriptomics datasets and sc/snRNA-seq datasets in our analysis.
**Table S3** Information on tissue regions in spatial transcriptomics datasets.

### Supplementary Figures

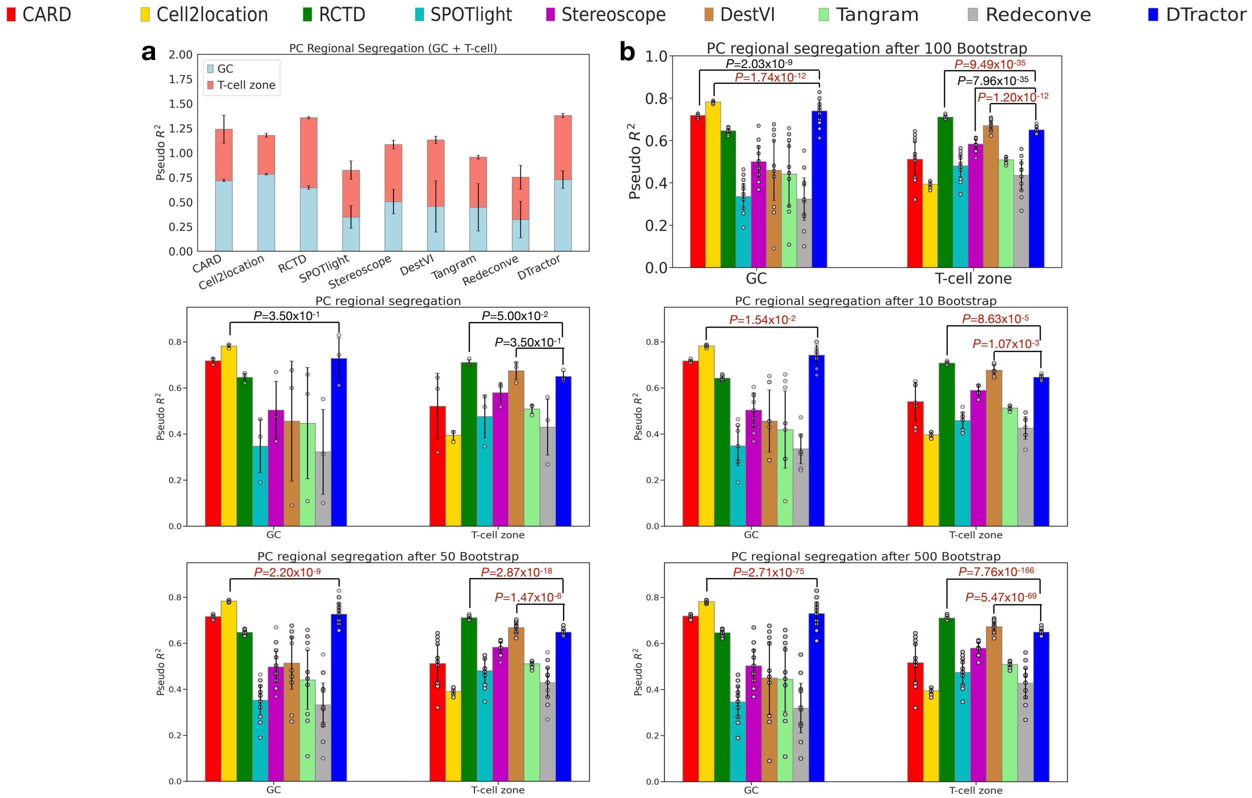


**Fig. S1 a,b, The height of each bar represents the average statistics, while the error bars indicate the standard errors. a, We calculated the mean of GC and T-cell zone values for each method, then, concatenated the two contributions into one bar per method, with different colors for GC and T-cell zone. Next, we used a stacked bar chart for visualization. b, Bootstrap resampling for groups of 10, 50, 100, and 500 samples, alongside the original plot without resampling, to assess regional segregation using $R^2$ by fitting logistic regression models for binary germinal center (GC) vs. non-GC regions and binary T-cell vs. non-T-cell regions on the principal component (PC) scores in the human lymph node. P-values are derived from one-sided Mann-Whitney U tests without adjustment. Dark red indicates p-values where other models outperformed, meeting the statistical significance threshold of 0.05. Each scatter dot represents a different sample.**

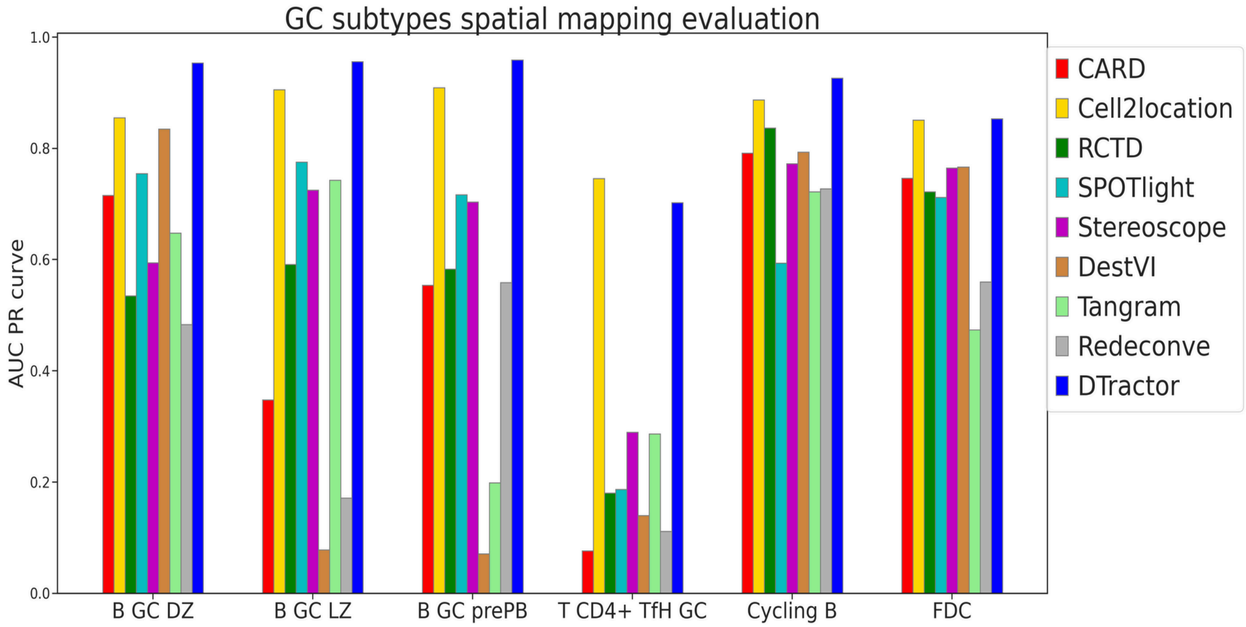


**Fig. S2** **Spatial mapping accuracy for six germinal center (GC) cell types, based on ground gruth annotations in the human lymph node. The height of each bar represents the average precision-recall scores area under the curve (AUC PR)**

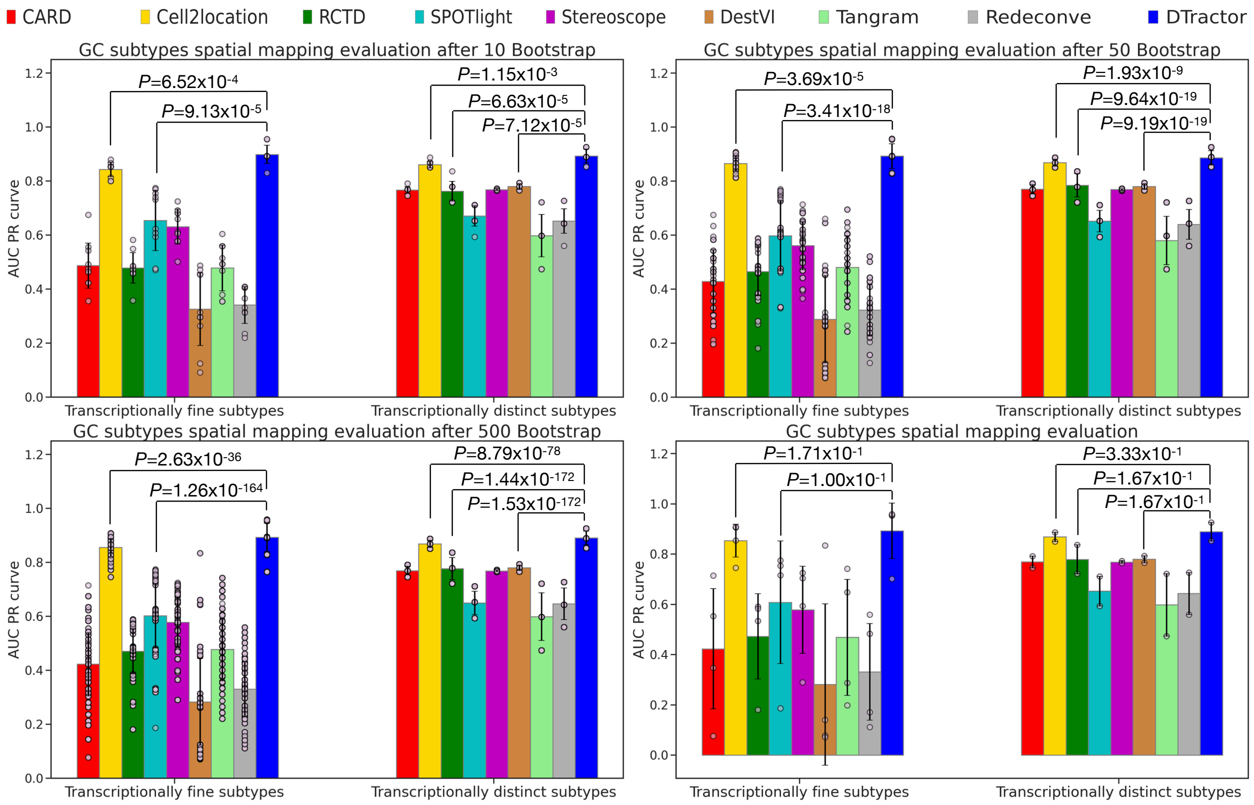


**Fig. S3** **Bootstrap resampling for groups of 10, 50, and 500 samples, alongside the original plot without resampling, to assess spatial mapping accuracy for the six germinal center (GC) expected cell types based on ground truth annotations in the human lymph node. P-values are derived from one-sided Mann-Whitney U tests without adjustment. The height of each bar represents the average precision-recall scores area under the curve (AUC PR), while each scatter dot represents a different sample and the error bars indicate the standard errors. In this context, the classification of six known GC cell types is determined by their transcriptional characteristics, distinguishing between those that are fine-grained and those that are distinct. The transcriptionally fine subtypes are B_GC_DZ, B_GC_LZ, B_GC_prePB, T_CD4+_TfH_GC, while the transcriptionally distinct subtypes are B_cycling, FDC. This classification relies on the Euclidean distance of their expression signatures to the closest other cell types as defined in Cell2location paper[1].**

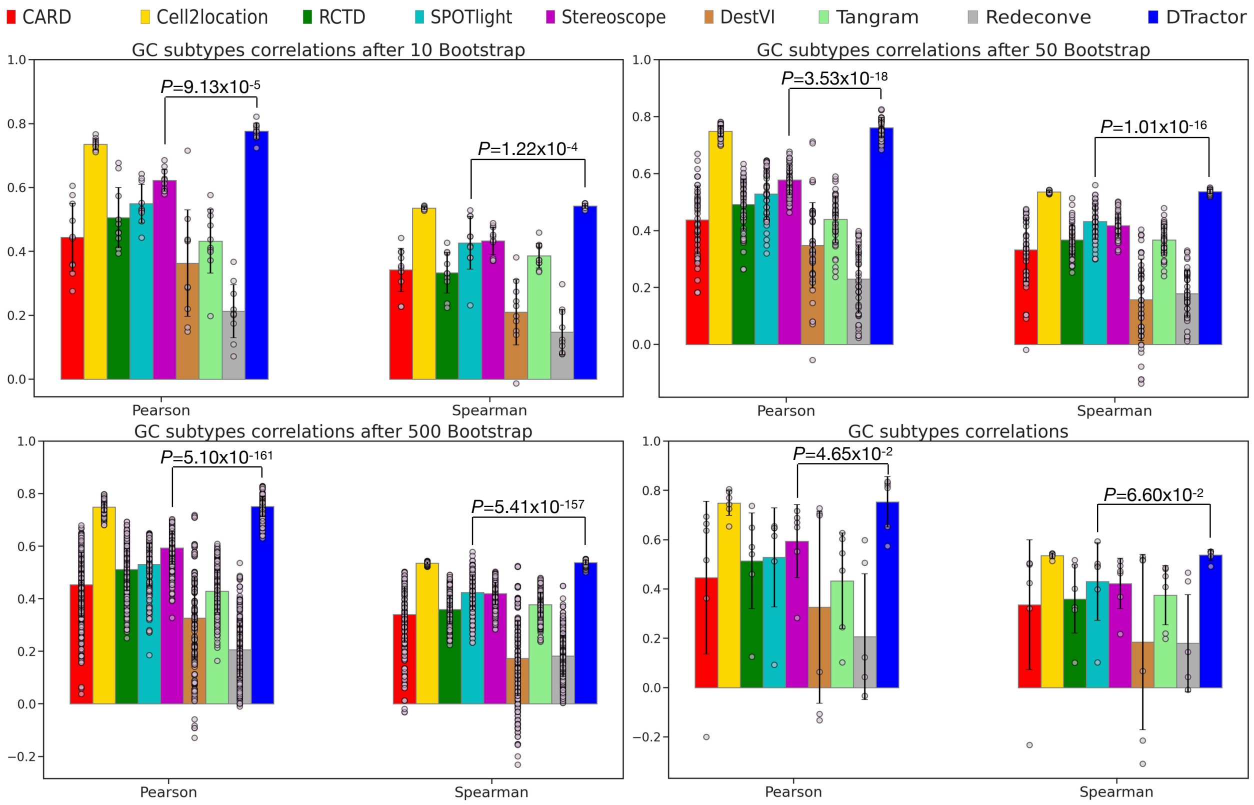


**Fig. S4 Bootstrap resampling for groups of 10, 50, and 500 samples, alongside the original plot without resampling, to assess macro-average correlation coefficients between the estimated cell type composition matrix of six germinal center (GC) expected cell types and the GC ground truth annotation in the human lymph node. P-values are derived from one-sided Mann-Whitney U tests without adjustment. The height of each bar represents the average statistics, while each scatter dot represents a different sample and the error bars indicate the standard errors.**

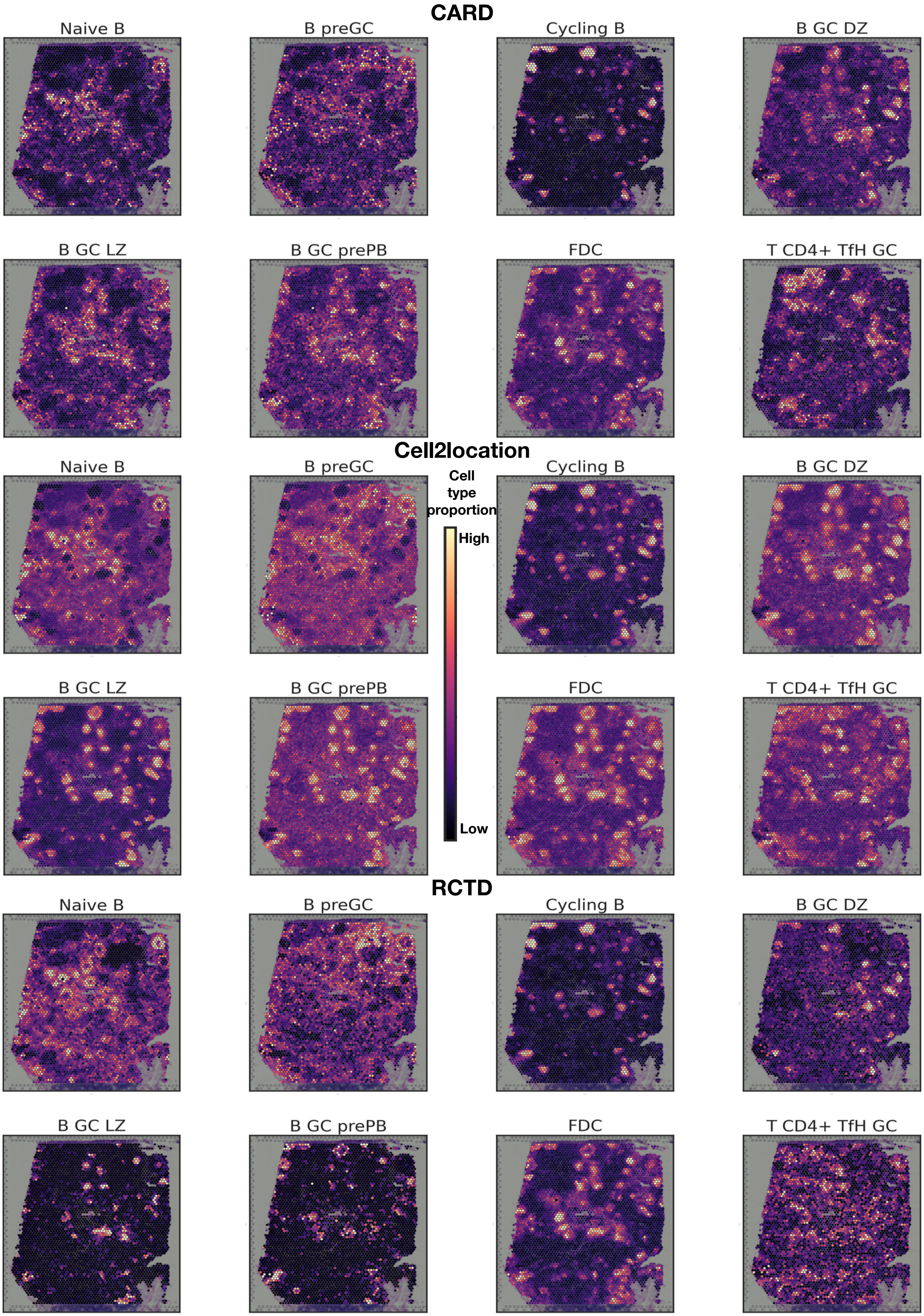
CARD
Naive B
B preGC
Cycling B
B GC DZ
B GC LZ
B GC prePB
FDC
T CD4+ TfH GC
Cell2location
Naive B
B preGC
Cell type proportion
High
Low
Cycling B
B GC DZ
B GC LZ
B GC prePB
FDC
T CD4+ TfH GC
RCTD
Naive B
B preGC
Cycling B
B GC DZ
B GC LZ
B GC prePB
FDC
T CD4+ TfH GC

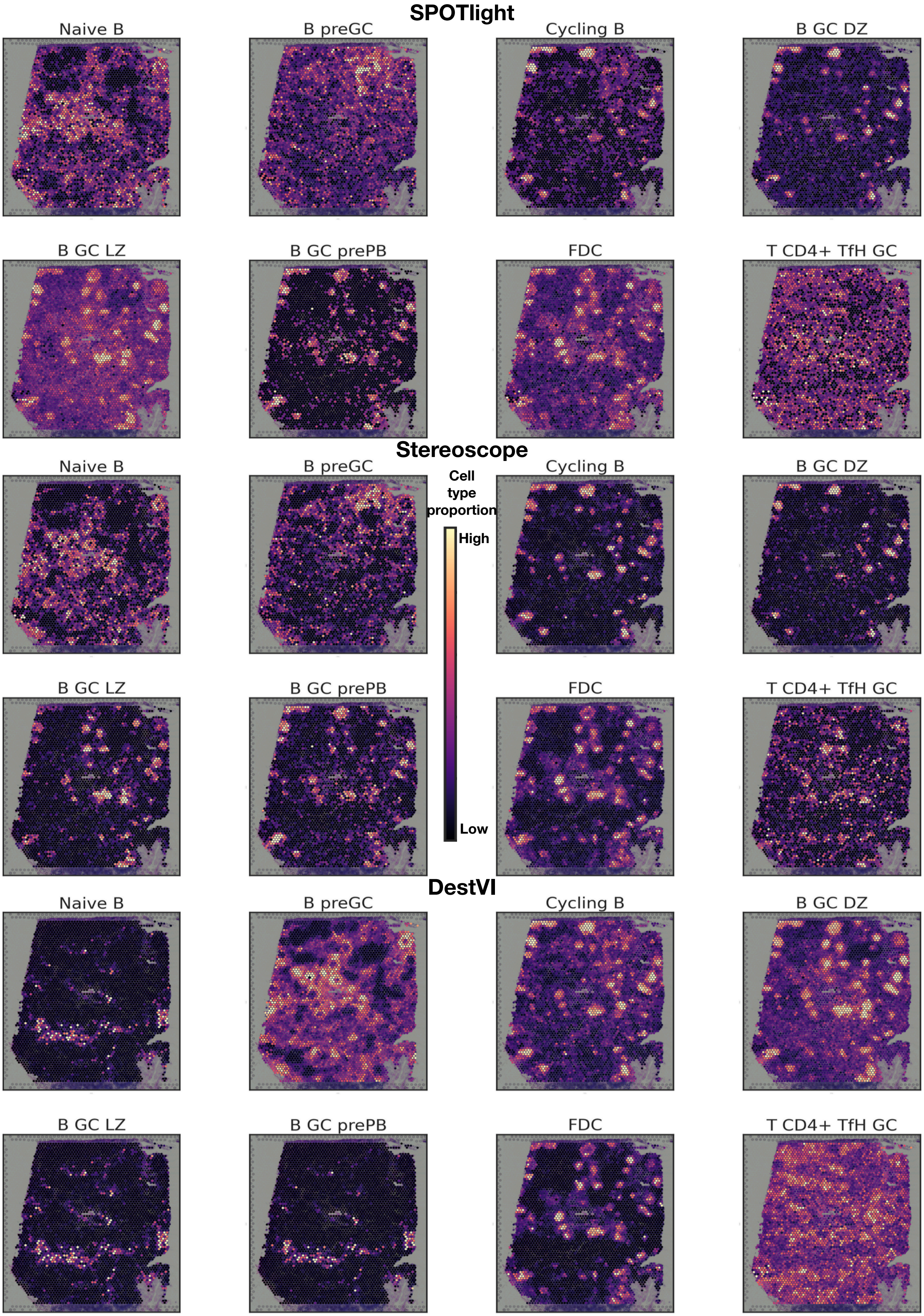
SPOTlight
Naive B
B preGC
Cycling B
B GC DZ
B GC LZ
B GC prePB
FDC
T CD4+ TfH GC
Stereoscope
Naive B
B preGC
Cycling B
B GC DZ
B GC LZ
B GC prePB
FDC
T CD4+ TfH GC
Cell type proportion
High
Low
DestVI
Naive B
B preGC
Cycling B
B GC DZ
B GC LZ
B GC prePB
FDC
T CD4+ TfH GC

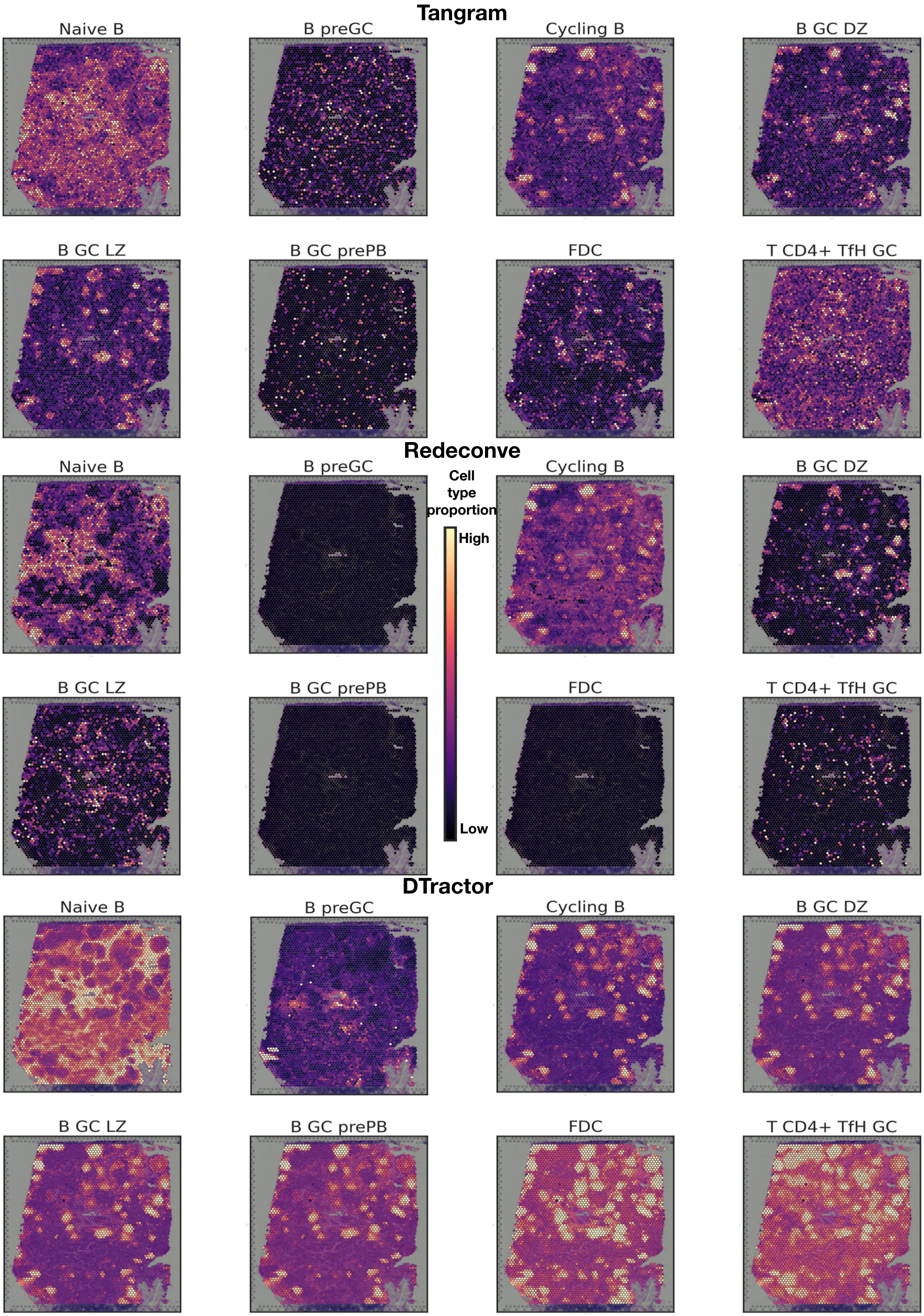




**Fig. S5** Estimated cell abundance of selected six known germinal center (GC) cell types along with Naive B and pre-GC B cell types in the human lymph node.

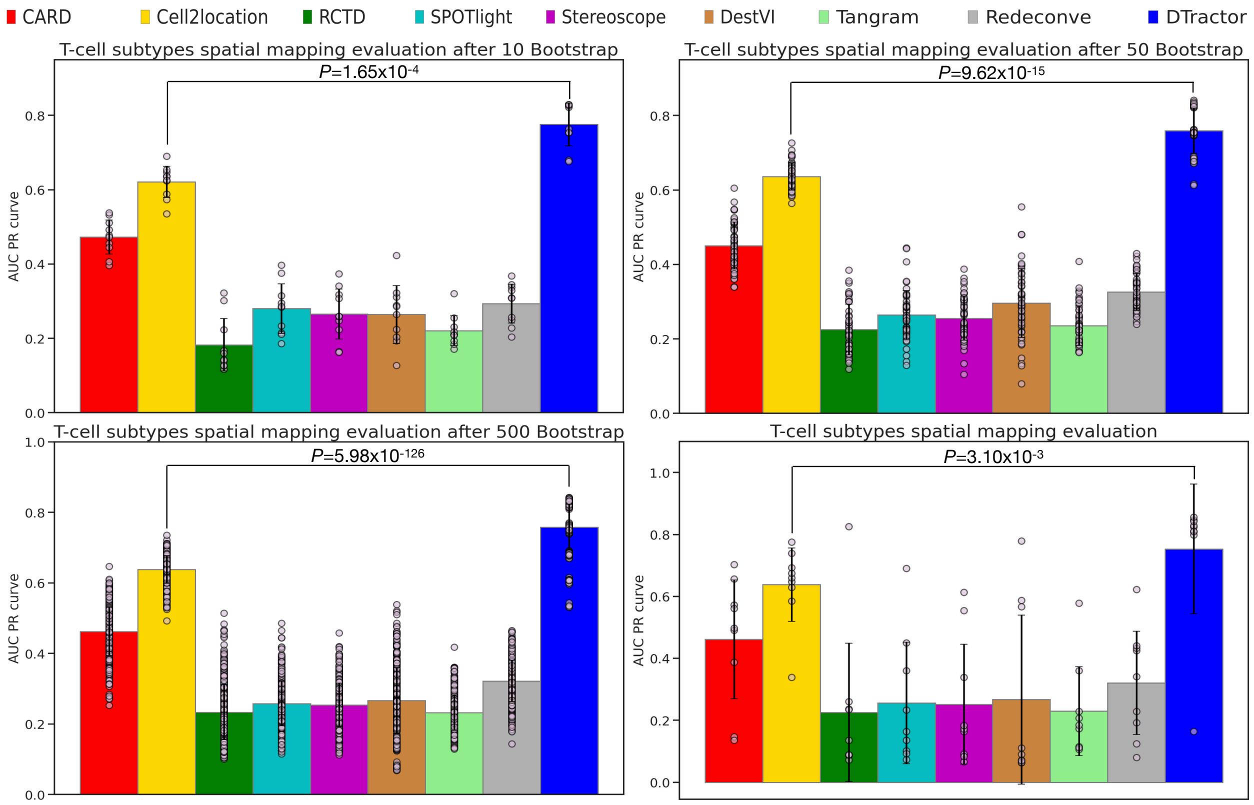


**Fig. S6 Bootstrap resampling for groups of 10, 50, and 500 samples, alongside the original plot without resampling, to assess spatial mapping accuracy for nine T cell types based on ground truth annotations in the human lymph node. P-values are derived from one-sided Mann-Whitney U tests without adjustment. The height of each bar represents the average precision-recall scores area under the curve (AUC PR), while each scatter dot represents a different sample and the error bars indicate the standard errors.**

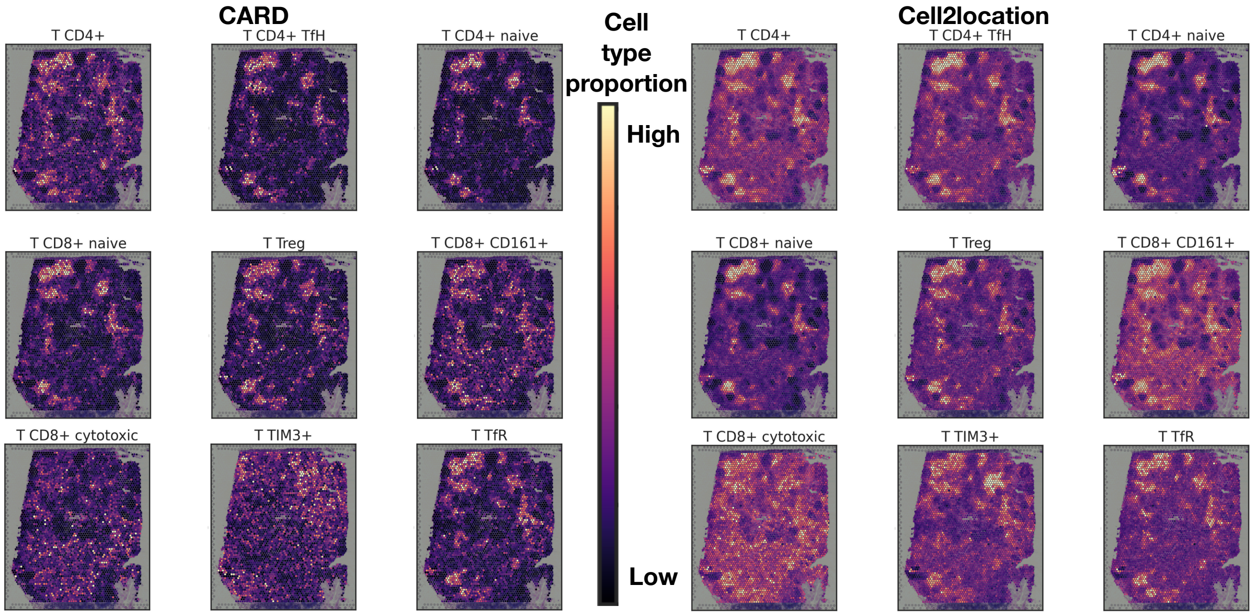

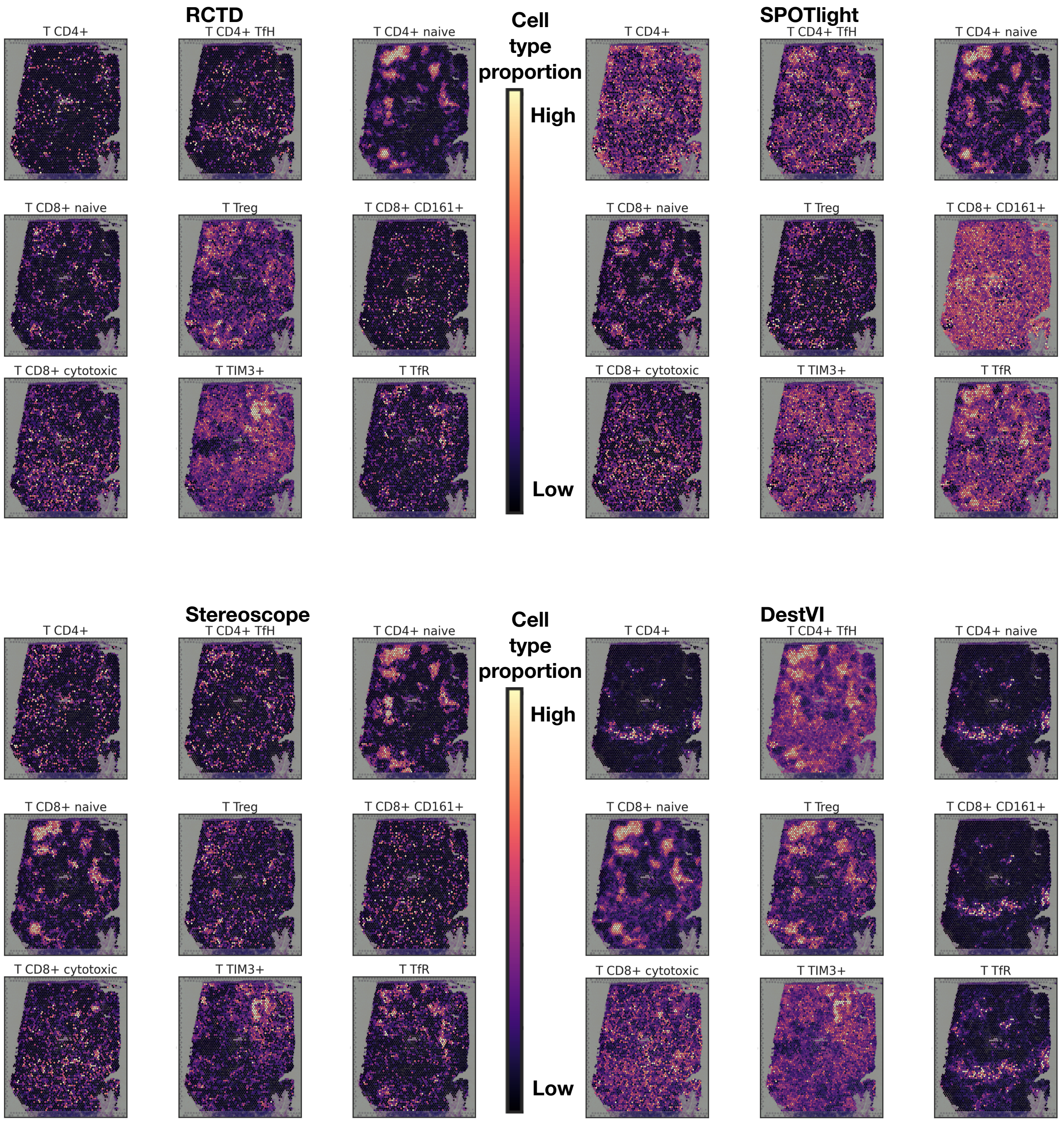
RCTD
T CD4+
T CD4+ TfH
T CD4+ naive
T CD8+ naive
T Treg
T CD8+ CD161+
T CD8+ cytotoxic
T TIM3+
T TfR
Cell type proportion
High
Low
SPOTlight
T CD4+
T CD4+ TfH
T CD4+ naive
T CD8+ naive
T Treg
T CD8+ CD161+
T CD8+ cytotoxic
T TIM3+
T TfR
Stereoscope
T CD4+
T CD4+ TfH
T CD4+ naive
T CD8+ naive
T Treg
T CD8+ CD161+
T CD8+ cytotoxic
T TIM3+
T TfR
Cell type proportion
High
Low
DestVI
T CD4+
T CD4+ TfH
T CD4+ naive
T CD8+ naive
T Treg
T CD8+ CD161+
T CD8+ cytotoxic
T TIM3+
T TfR

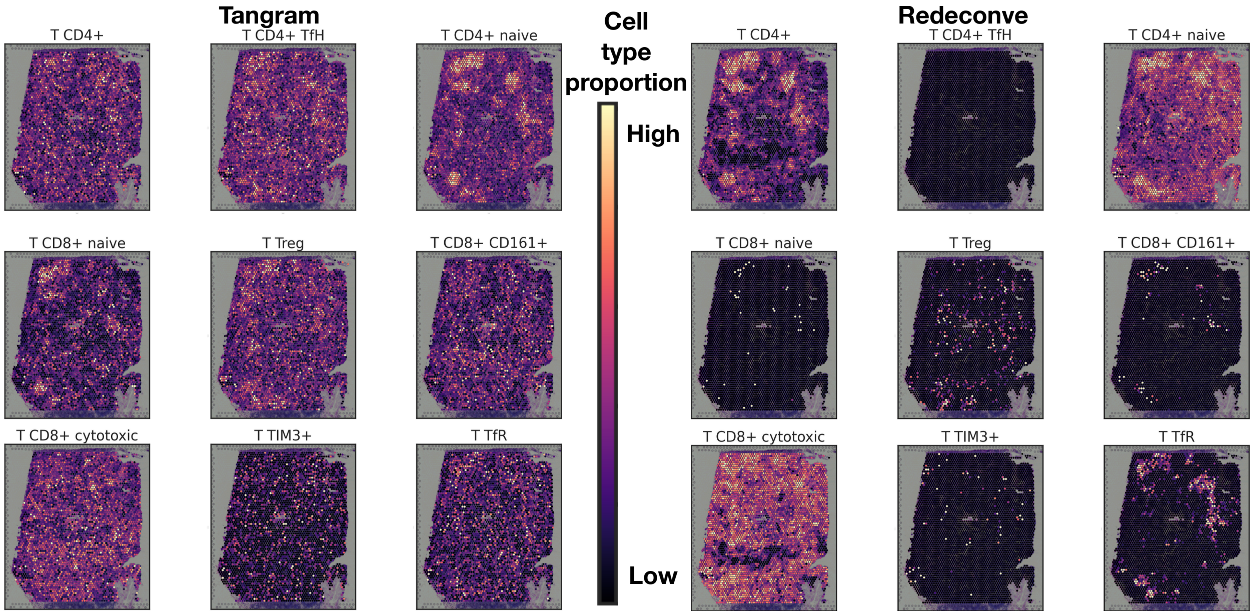


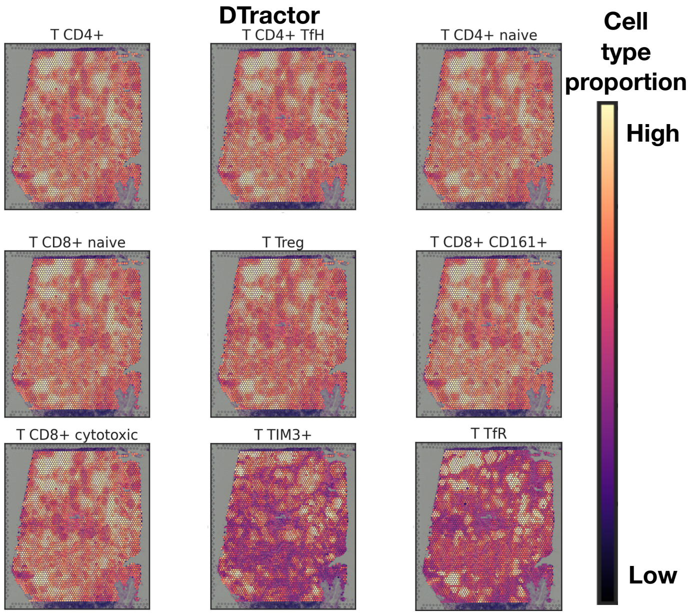


**Fig. S7 Estimated cell abundance of selected T cell types in the human lymph node.**

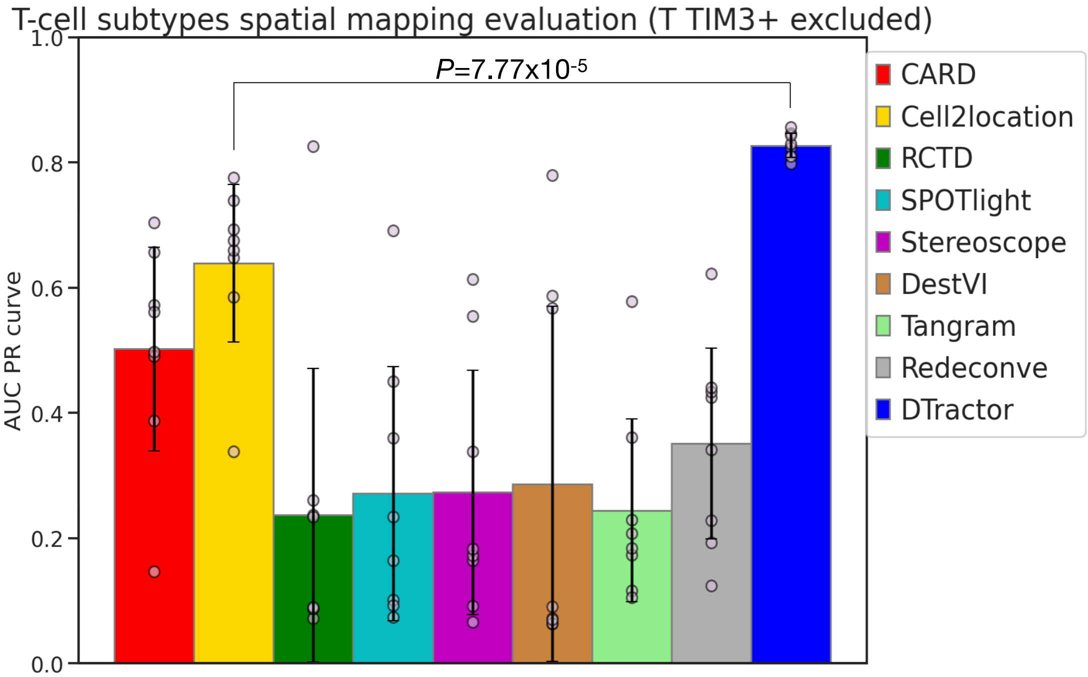


**Fig. S8** Spatial mapping accuracy for eight T cell types, excluding the T TIM3+ subtype, based on ground truth annotations in the human lymph node. P-values are derived from one-sided Mann-Whitney U tests without adjustment. The height of each bar represents the average precision-recall scores area under the curve (AUC PR), while each scatter dot represents a different sample and the error bars indicate the standard errors.

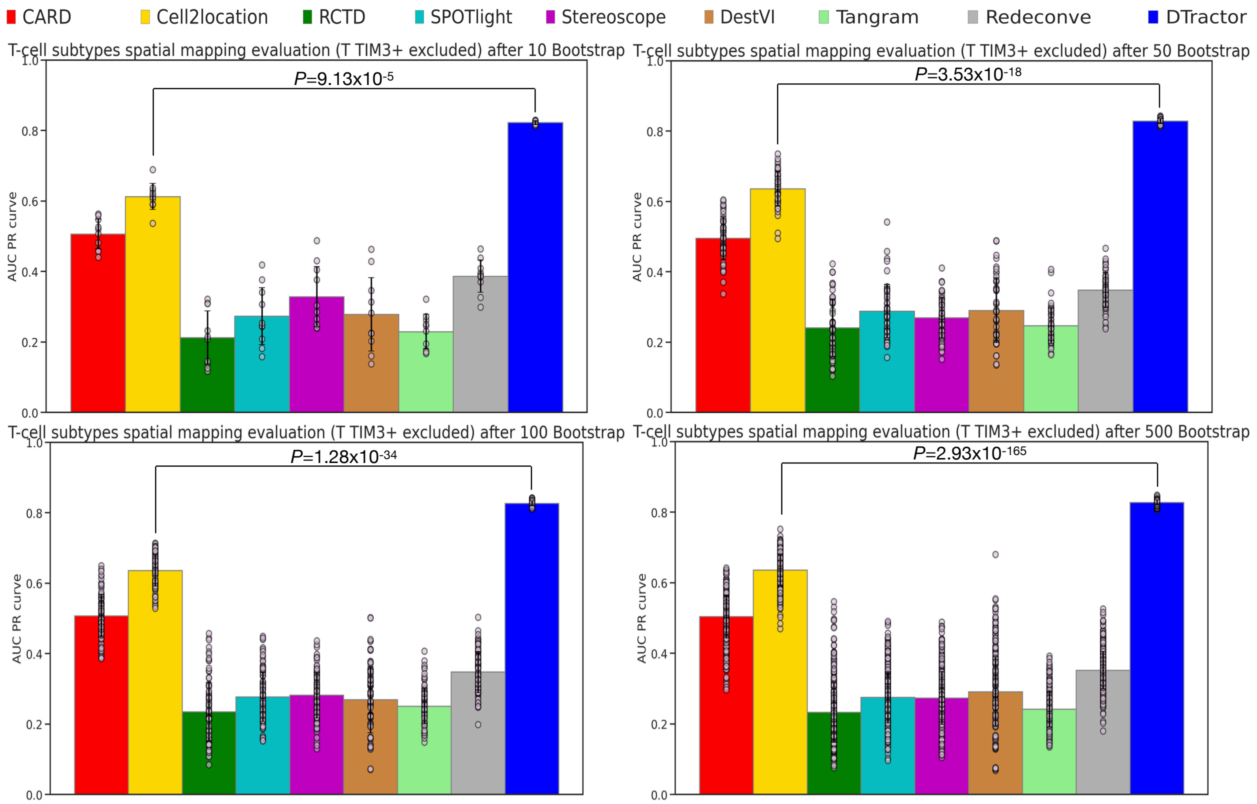


**Fig. S9** **Bootstrap resampling for groups of 10, 50, 100, 500 bootstrap samples to assess spatial mapping accuracy for eight T cell types excluding the T TIM3+ subtype based on ground truth annotations in the human lymph node. P-values are derived from one-sided Mann-Whitney U tests without adjustment. The height of each bar represents the average precision-recall scores area under the curve (AUC PR), while each scatter dot represents a different sample and the error bars indicate the standard errors.**

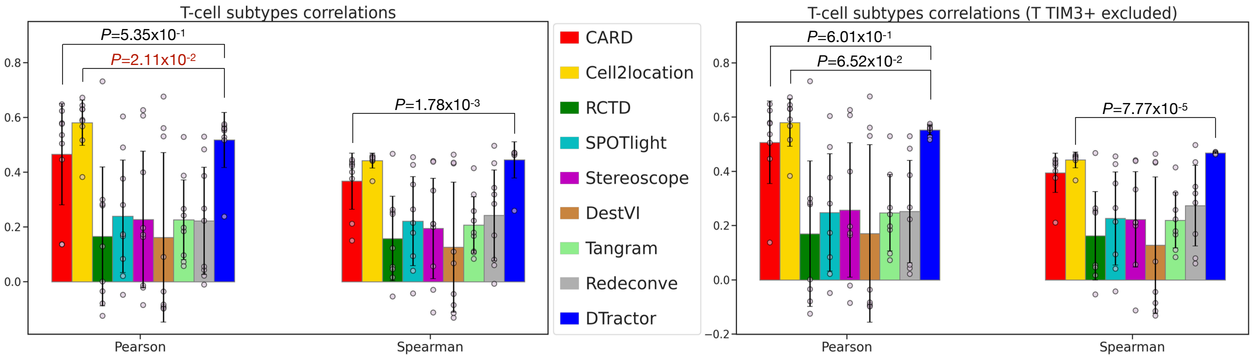


**Fig. S10** **Macro average correlation coefficients for all subtypes in the T-cell zones against ground truth annotations, as well as with the T TIM3+ subtype excluded in the human lymph node. P-values are derived from one-sided Mann-Whitney U tests without adjustment. Dark red indicates p-values where other models outperformed, meeting the statistical significance threshold of 0.05. The height of each bar represents the average statistics, while each scatter dot represents a different sample and the error bars indicate the standard errors.**

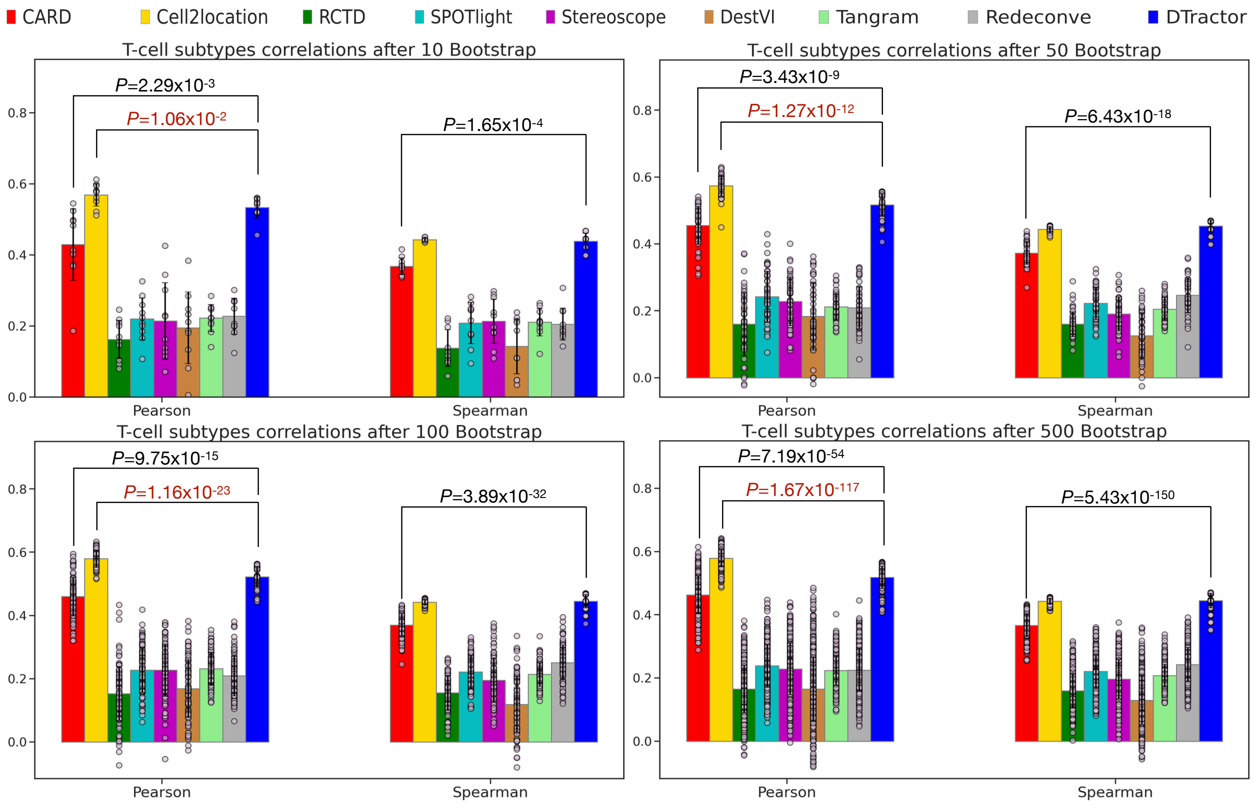


**Fig. S11 Bootstrap resampling for groups of 10, 50, 100, 500 bootstrap samples to assess macro-average correlation coefficients between the estimated cell type composition matrix of all subtypes in the T-cell zone and the T-cell zone ground truth annotation in the human lymph node. P-values are derived from one-sided Mann-Whitney U tests without adjustment. We used dark red font to highlight p-values where other models outperformed in the original plot prior to bootstrap resampling. The height of each bar represents the average statistics, while each scatter dot represents a different sample and the error bars indicate the standard errors.**

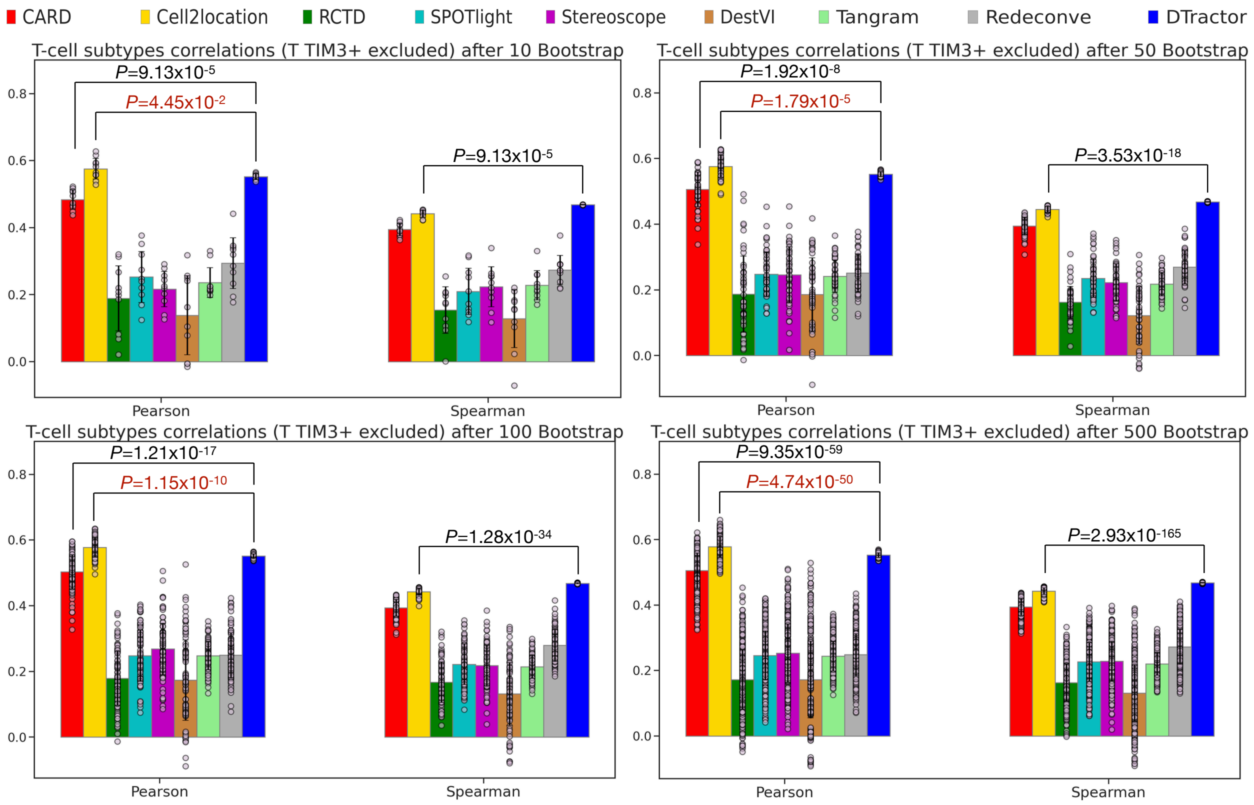


**Fig. S12 Bootstrap resampling for groups of 10, 50, 100, 500 bootstrap samples to assess macro-average correlation coefficients between the estimated cell type composition matrix of eight subtypes in the T-cell zone (excluding the T TIM3+ subtype) and the ground truth annotation of the T-cell zone in the human lymph node. P-values are derived from one-sided Mann-Whitney U tests without adjustment. We used dark red font to highlight p-values where other models outperformed in the original plot prior to bootstrap resampling. The height of each bar represents the average statistics, while each scatter dot represents a different sample and the error bars indicate the standard errors.**

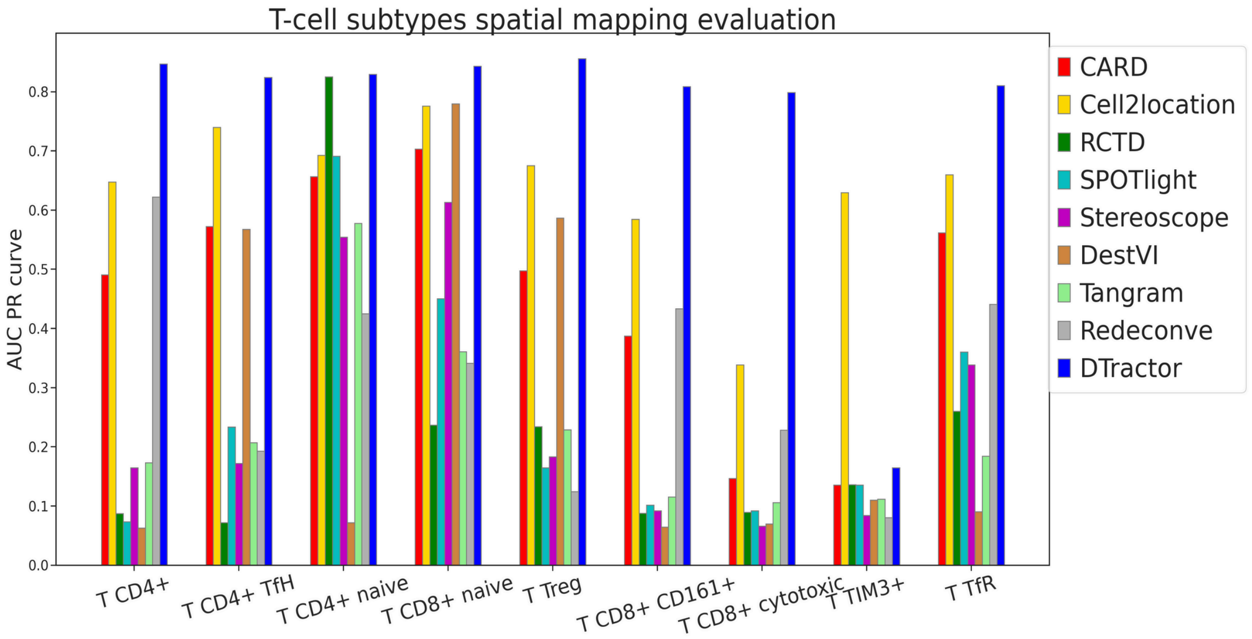


**Fig. S13 Spatial mapping accuracy for nine T cell types based on ground truth annotations in the human lymph node. The height of each bar represents average precision-recall scores area under the curve (AUC PR)**

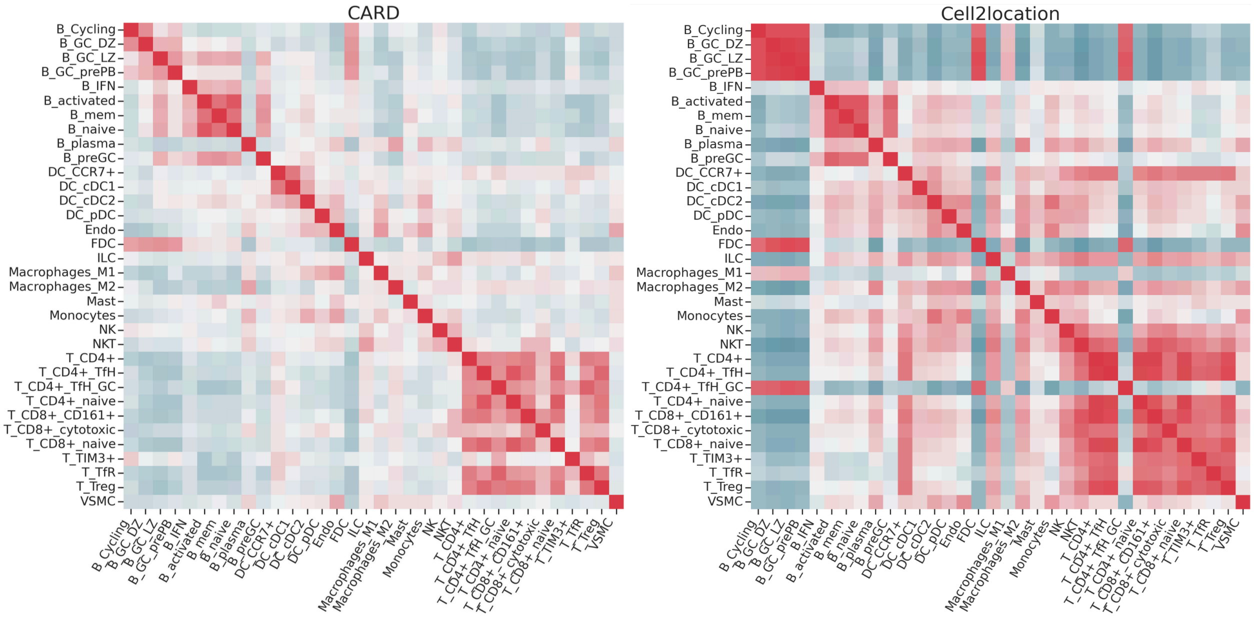

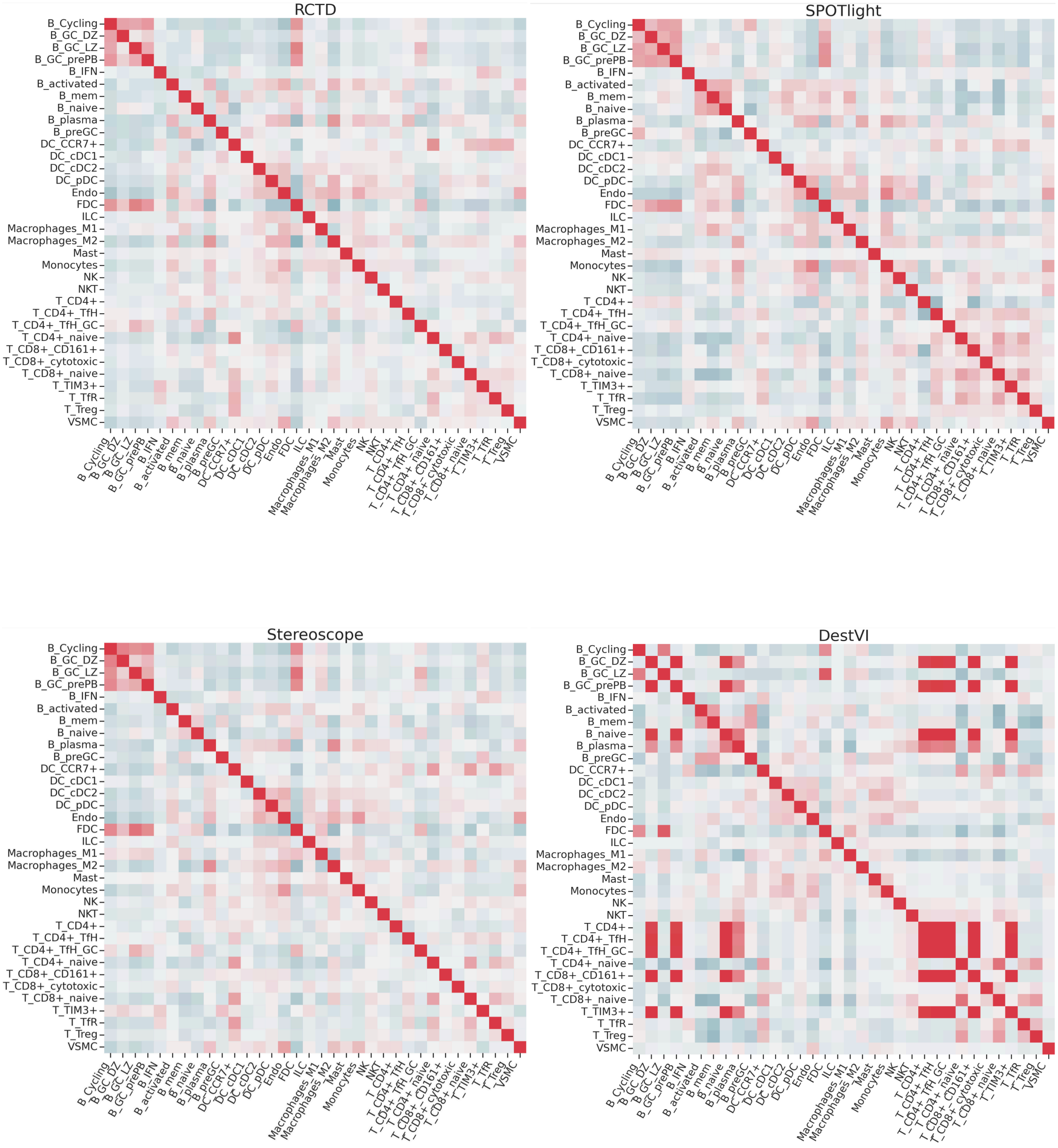
RCTD
SPOTlight
Stereoscope
DestVI
B_Cycling
B_GC_DZ
B_GC_LZ
B_GC_prePB
B_IFN
B_activated
B_mem
B_naive
B_plasma
B_preGC
DC_CCR7+
DC_cDC1
DC_cDC2
DC_pDC
Endo
FDC
ILC
Macrophages_M1
Macrophages_M2
Mast
Monocytes
NK
NKT
T_CD4+
T_CD4+_TfH
T_CD4+_TfH_GC
T_CD4+_naive
T_CD8+_CD161+
T_CD8+_cytotoxic
T_CD8+_naive
T_TIM3+
T_TfR
T_Treg
VSMC

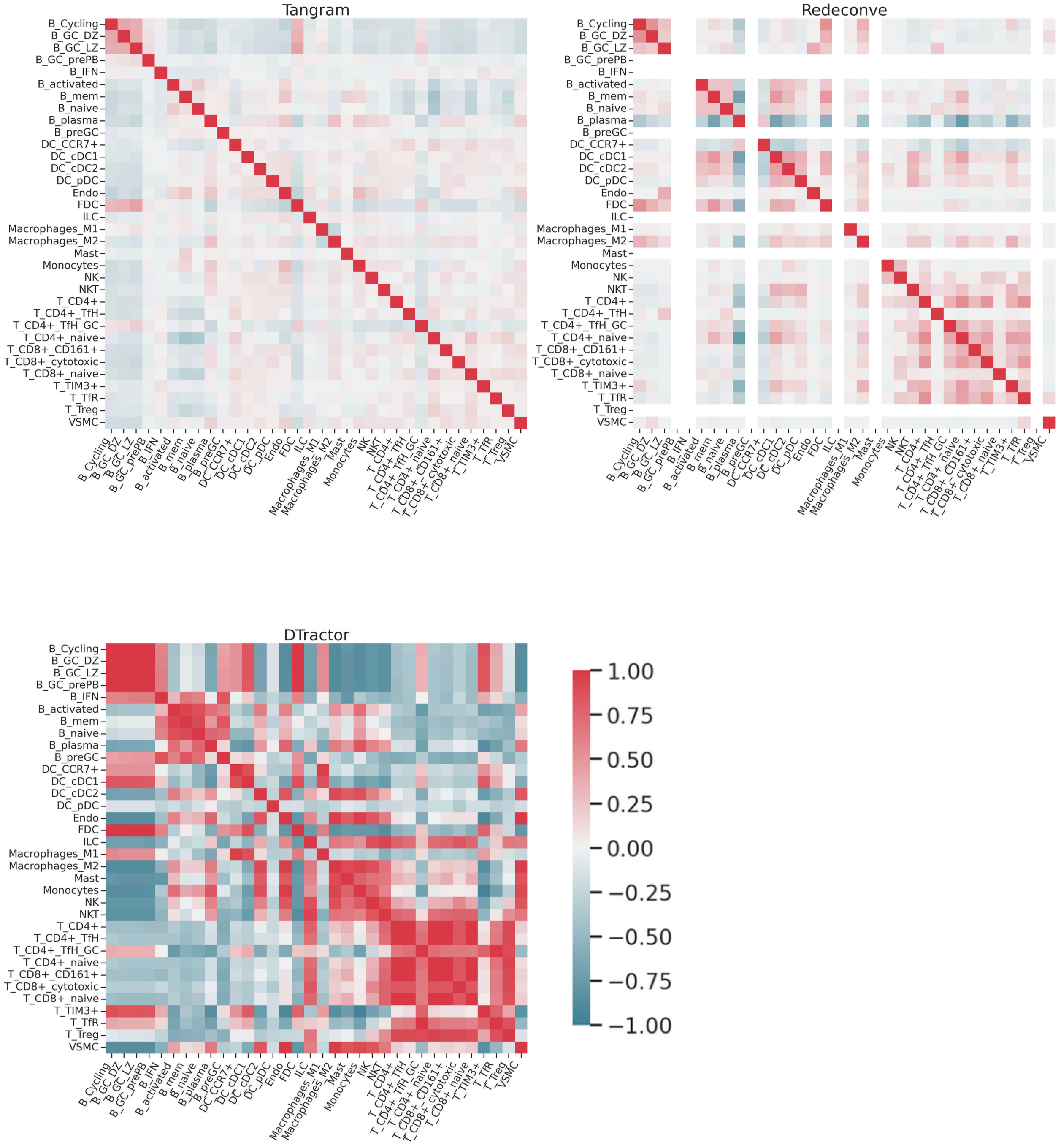


**Fig. S14 Correlations in the estimated cell type composition matrix across spatial locations between pairs of cell types are compared across different algorithms in the human lymph node. The color of each comparison is scaled based on the correlation value.**

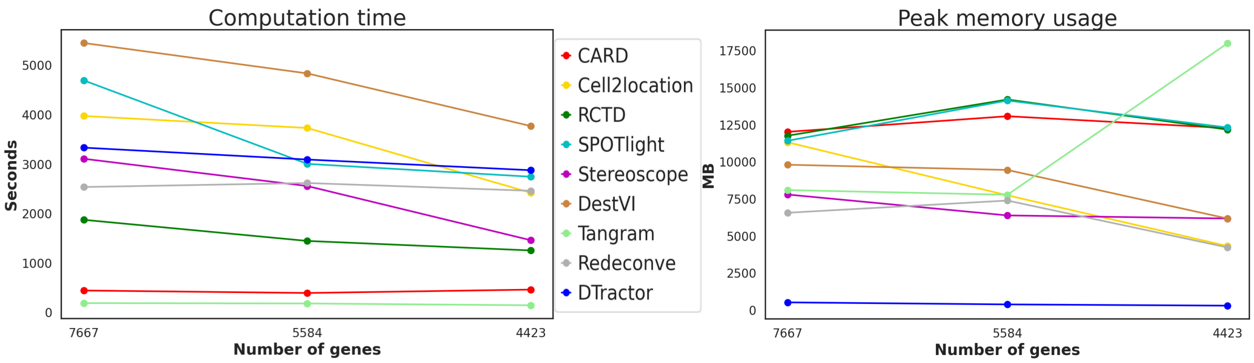


**Fig. S15 Computation time (seconds) and peak memory usage (MB) for each deconvolution method for the human lymph node. To ensure fairness and consistency, we adhered to the default settings for all methods when measuring time and memory usage, as outlined in the "Parameter setting" section under "Benchmarking" in the Methods.**

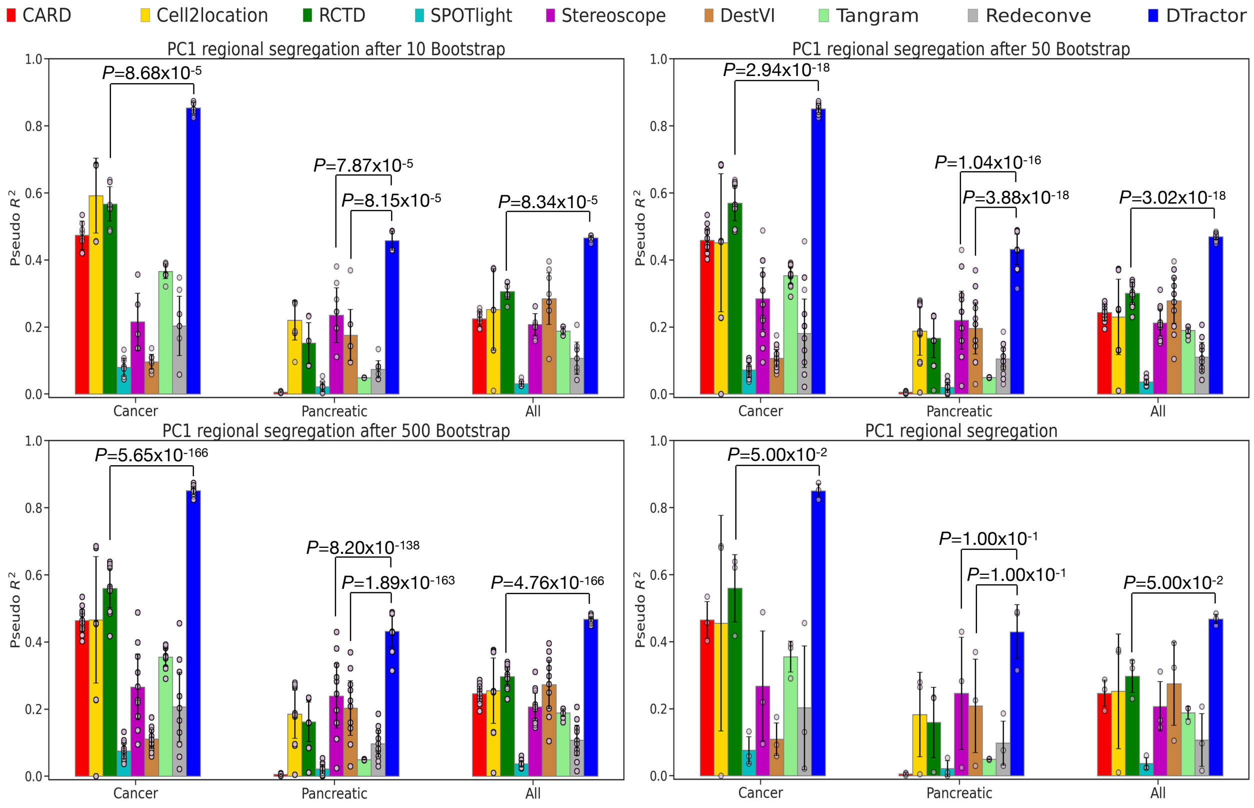


**Fig. S16 Bootstrap resampling for groups of 10, 50, and 500 samples, alongside the original plot without resampling, to assess regional segregation of cancer, pancreatic, and all four regions using $R^2$ by fitting logistic regression models for binary cancer vs. non-cancer regions, binary pancreatic vs. non-pancreatic regions, and multiclass classification for all four regions on the principal component (PC) scores in PCAC. P-values are derived from one-sided Mann-Whitney U tests without adjustment. The height of each bar represents the average statistics, while each scatter dot represents a different sample and the error bars indicate the standard errors.**

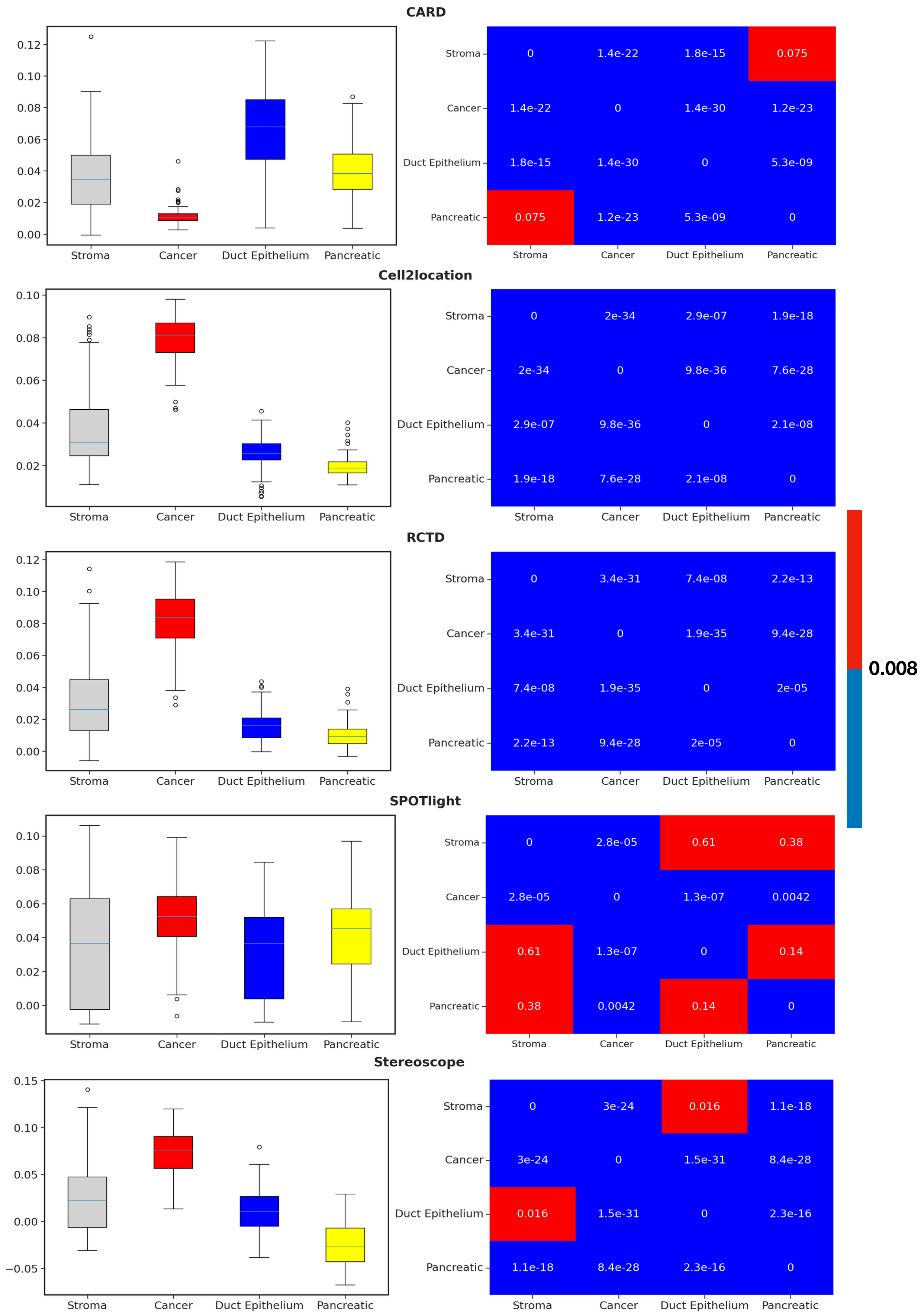

CARD
Stroma
Cancer
Duct Epithelium
Pancreatic
0 1.4e-22 1.8e-15 0.075
1.4e-22 0 1.4e-30 1.2e-23
1.8e-15 1.4e-30 0 5.3e-09
0.075 1.2e-23 5.3e-09 0
Cell2location
0 2e-34 2.9e-07 1.9e-18
2e-34 0 9.8e-36 7.6e-28
2.9e-07 9.8e-36 0 2.1e-08
1.9e-18 7.6e-28 2.1e-08 0
RCTD
0 3.4e-31 7.4e-08 2.2e-13
3.4e-31 0 1.9e-35 9.4e-28
7.4e-08 1.9e-35 0 2e-05
2.2e-13 9.4e-28 2e-05 0
0.008
SPOTlight
0 2.8e-05 0.61 0.38
2.8e-05 0 1.3e-07 0.0042
0.61 1.3e-07 0 0.14
0.38 0.0042 0.14 0
Stereoscope
0 3e-24 0.016 1.1e-18
3e-24 0 1.5e-31 8.4e-28
0.016 1.5e-31 0 2.3e-16
1.1e-18 8.4e-28 2.3e-16 0

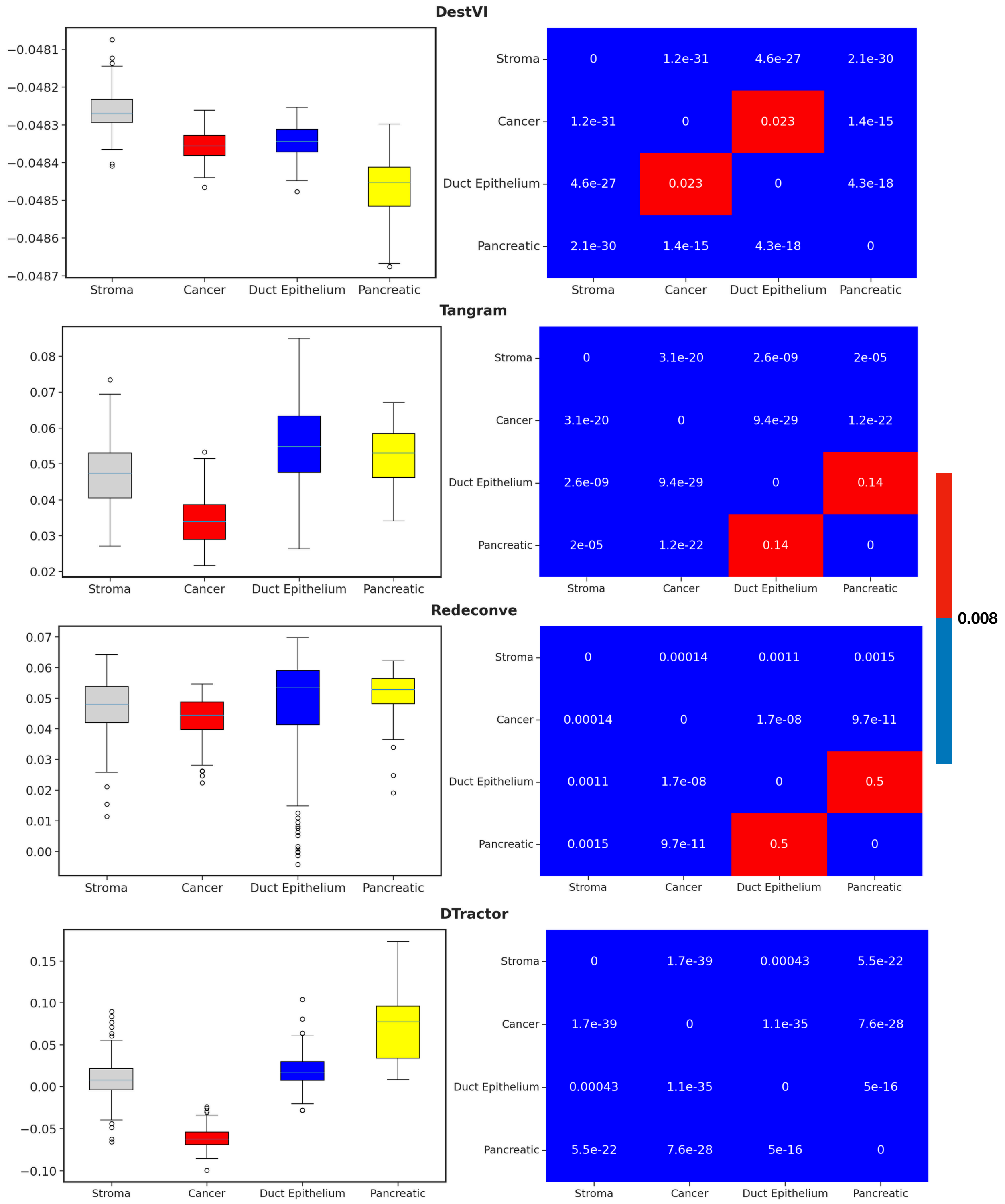


**Fig. S17** Boxplot and heatmap of pairwise two-sided Wilcoxon-rank sum test using the first principal component (PC1) of the estimated cell type composition matrix to discern how the PC1 differentiates between the four tissue regions, aligning with the ground truth annotations for PDAC (13402 shared genes). After applying the Bonferroni correction (p-value threshold of $0.05/6 = 0.008$), p-values less than 0.008 are considered significant and are highlighted blue. DTractor clearly distinguishes between each region in all four. Furthermore, these boxplot demonstrates that the PC1 scores effectively capture the relative regional distances.

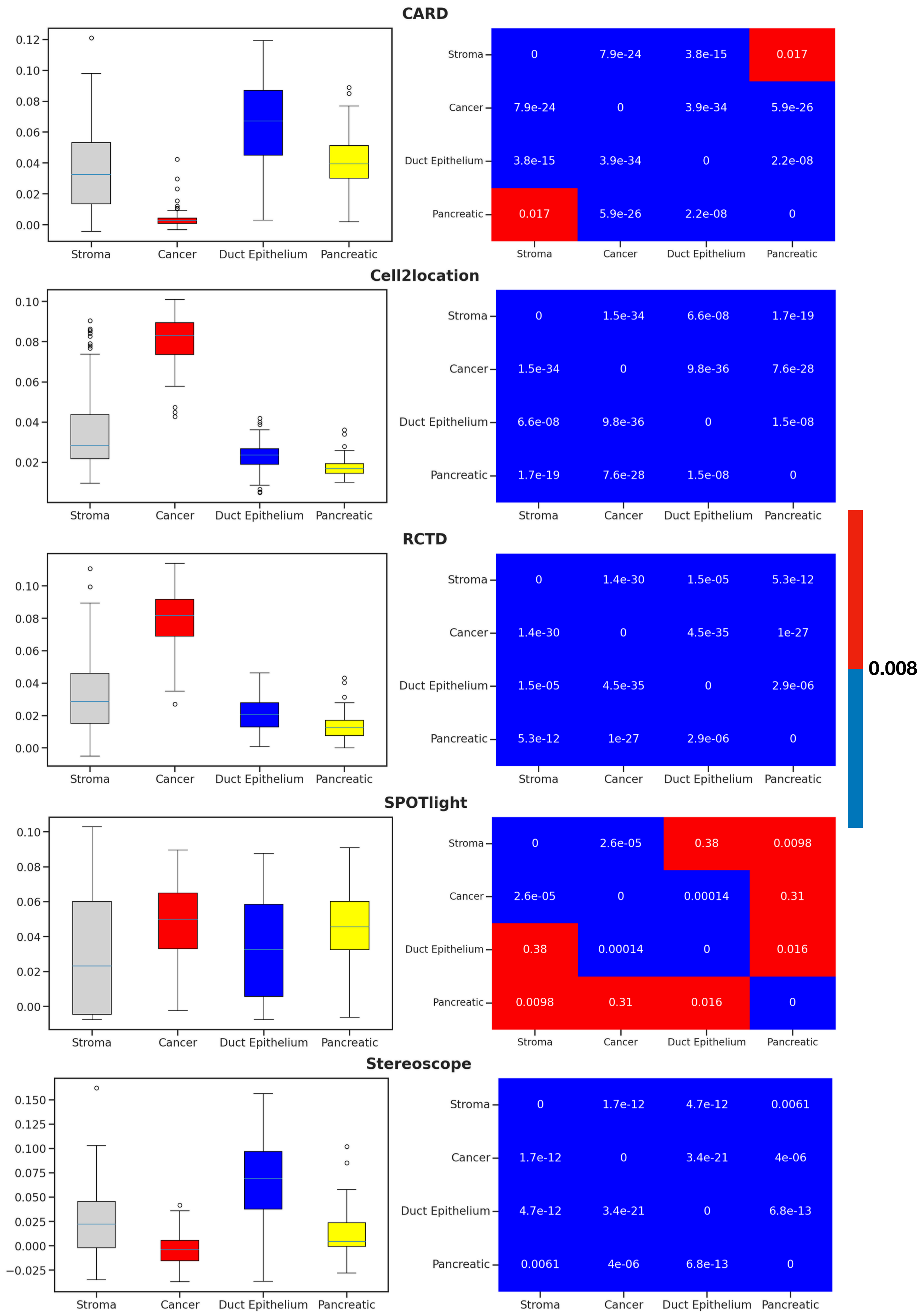
CARD
0.12
0.10
0.08
0.06
0.04
0.02
0.00
Stroma
Cancer
Duct Epithelium
Pancreatic
0 | 7.9e-24 | 3.8e-15 | 0.017
7.9e-24 | 0 | 3.9e-34 | 5.9e-26
3.8e-15 | 3.9e-34 | 0 | 2.2e-08
0.017 | 5.9e-26 | 2.2e-08 | 0
Cell2location
0.10
0.08
0.06
0.04
0.02
0 | 1.5e-34 | 6.6e-08 | 1.7e-19
1.5e-34 | 0 | 9.8e-36 | 7.6e-28
6.6e-08 | 9.8e-36 | 0 | 1.5e-08
1.7e-19 | 7.6e-28 | 1.5e-08 | 0
RCTD
0 | 1.4e-30 | 1.5e-05 | 5.3e-12
1.4e-30 | 0 | 4.5e-35 | 1e-27
1.5e-05 | 4.5e-35 | 0 | 2.9e-06
5.3e-12 | 1e-27 | 2.9e-06 | 0
0.008
SPOTlight
0 | 2.6e-05 | 0.38 | 0.0098
2.6e-05 | 0 | 0.00014 | 0.31
0.38 | 0.00014 | 0 | 0.016
0.0098 | 0.31 | 0.016 | 0
Stereoscope
0.150
0.125
0.100
0.075
0.050
0.025
0.000
−0.025
0 | 1.7e-12 | 4.7e-12 | 0.0061
1.7e-12 | 0 | 3.4e-21 | 4e-06
4.7e-12 | 3.4e-21 | 0 | 6.8e-13
0.0061 | 4e-06 | 6.8e-13 | 0

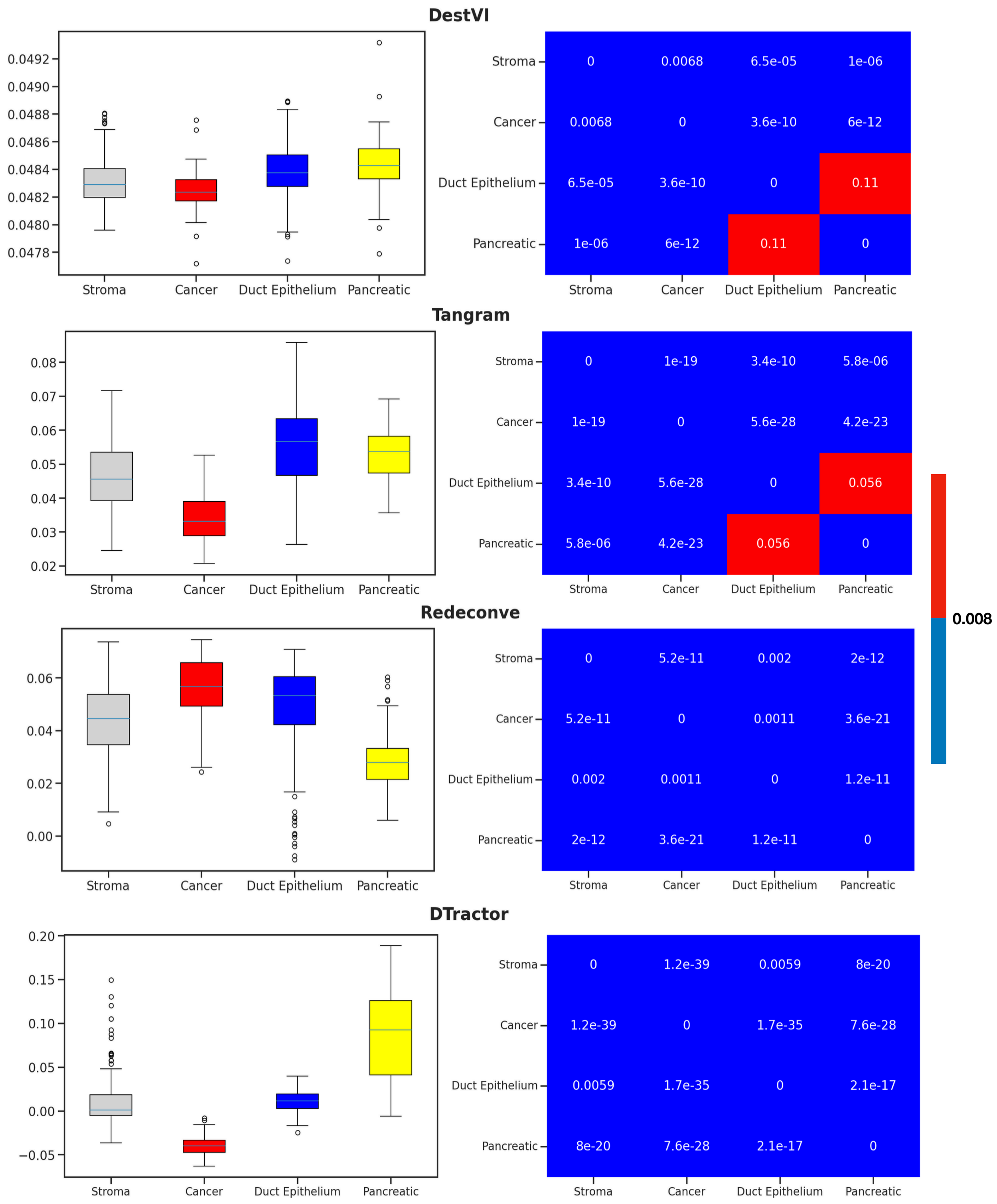


**Fig. S18** Boxplot and heatmap of pairwise two-sided Wilcoxon-rank sum test using the first principal component (PC1) of the estimated cell type composition matrix to discern how the PC1 differentiates between the four tissue regions, aligning with the ground truth annotations for PDAC (9818 shared genes). After applying the Bonferroni correction (p-value threshold of $0.05/6 = 0.008$), p-values less than 0.008 are considered significant and are highlighted blue. DTractor clearly distinguishes between each region in all four. Furthermore, these boxplot demonstrates that the PC1 scores effectively capture the relative regional distances.

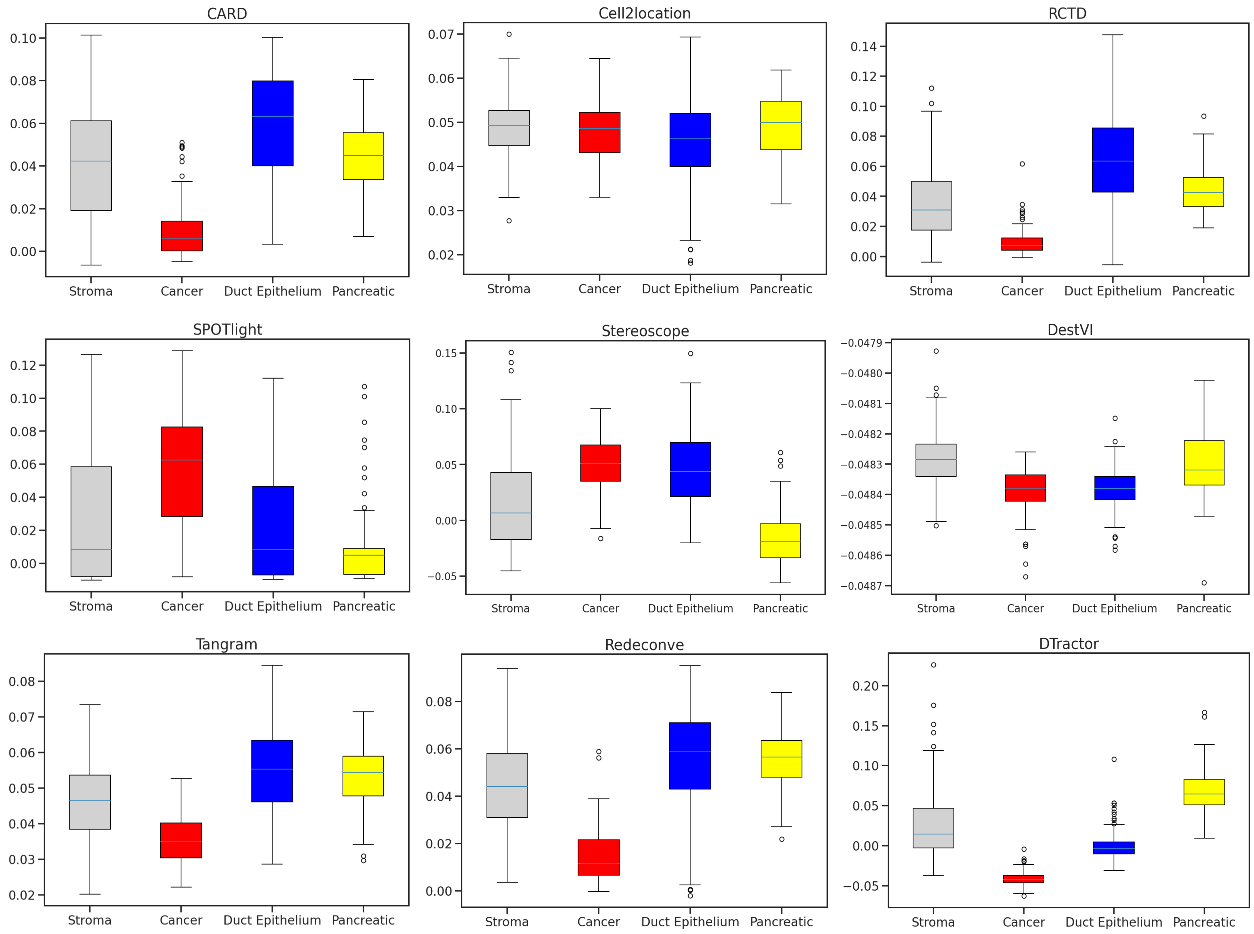


**Fig. S19** **Boxplot of the first principal component (PC1) scores of the estimated cell type composition matrix to discern how the PC1 differentiates between the four tissue regions, aligning with the ground truth annotations for PDAC (8729 shared genes). DTractor successfully distinguished four different types of tissue regions consistent with histological analysis annotations. Furthermore, these boxplot demonstrates that the PC1 scores effectively capture the relative regional distances.**

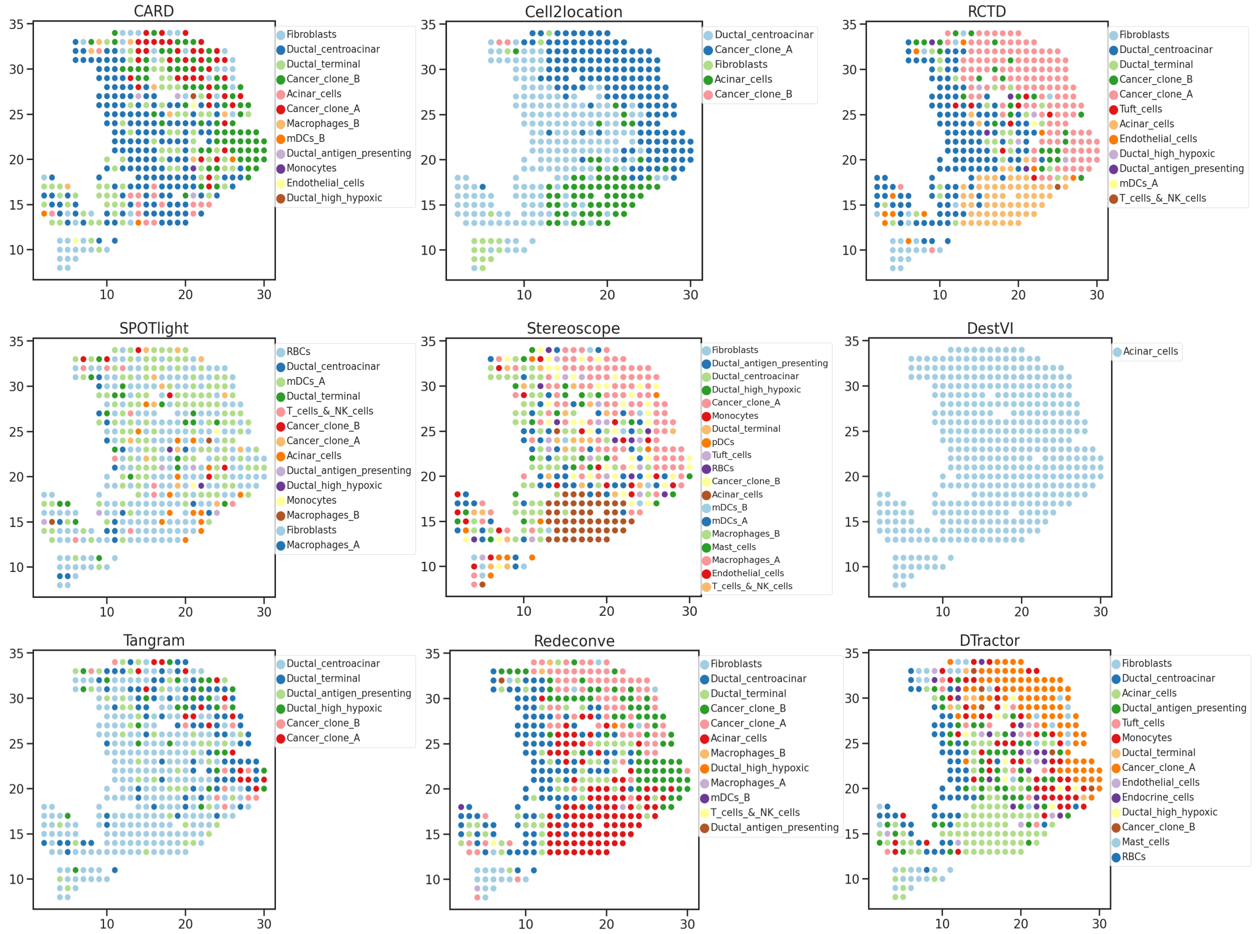


**Fig. S20 Inferred dominant cell type of the estimated cell type composition for each location in PDAC (13402 shared genes).**

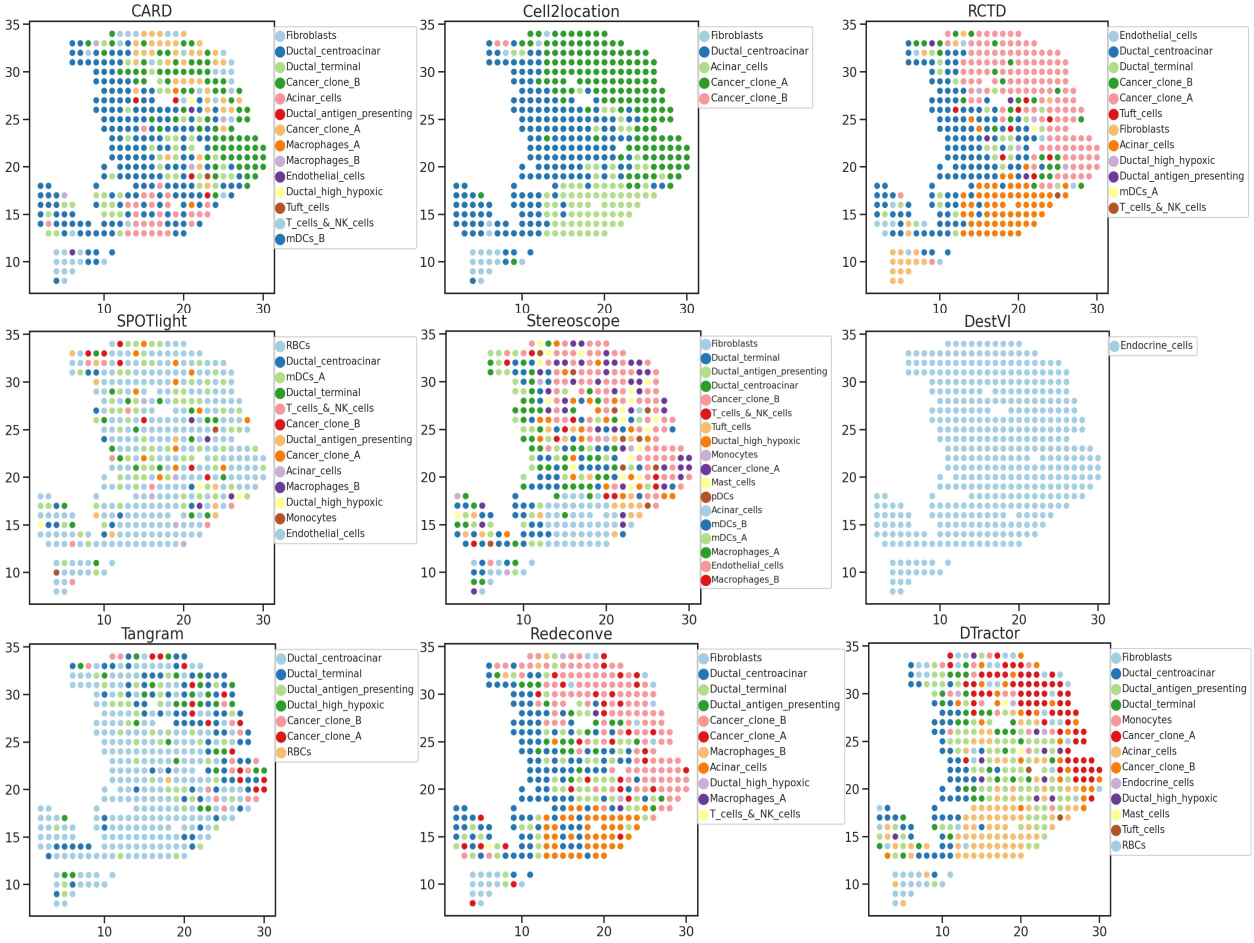


**Fig. S21 Inferred dominant cell type of the estimated cell type composition for each location in PDAC (9818 shared genes).**

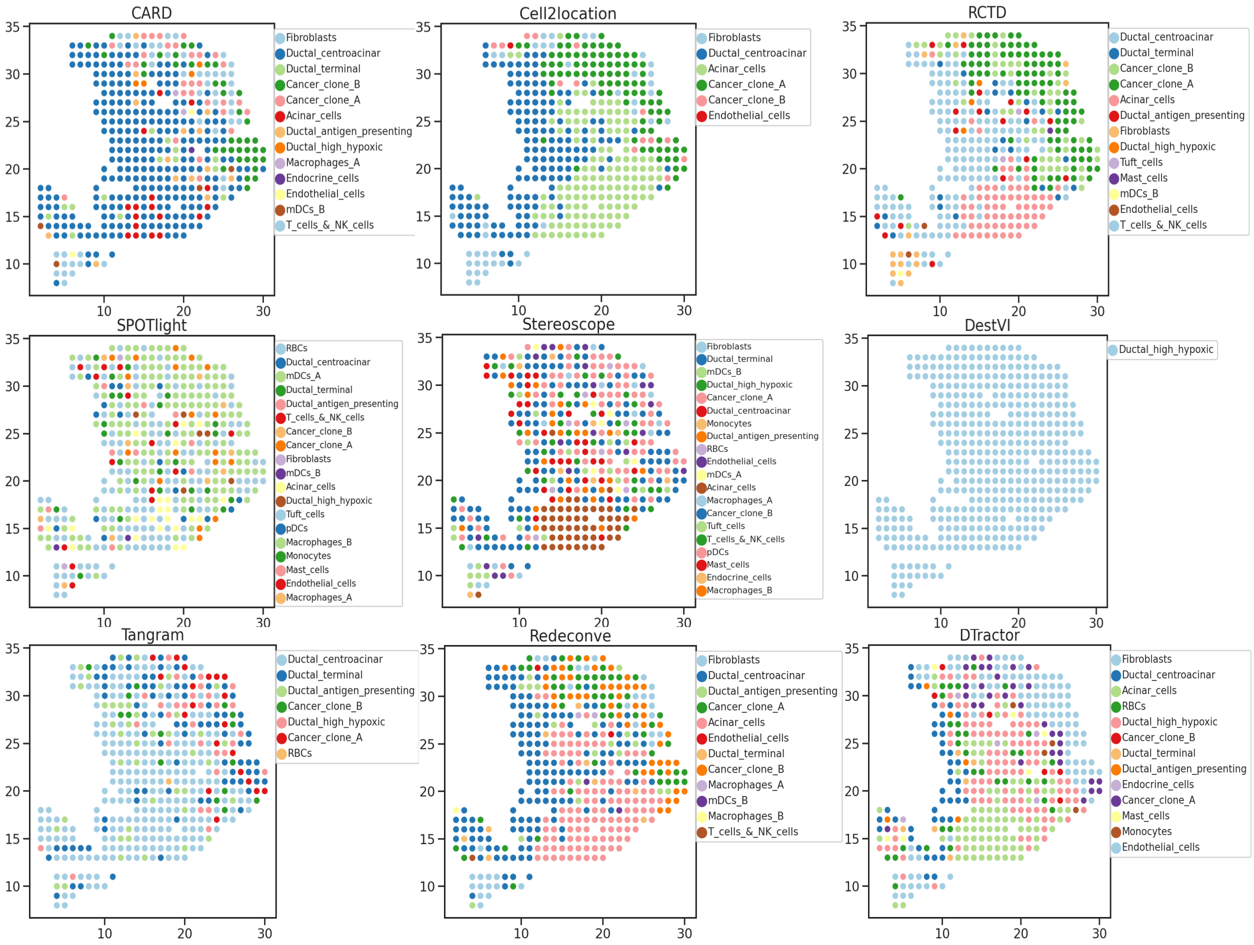


**Fig. S22 Inferred dominant cell type of the estimated cell type composition for each location in PDAC (8729 shared genes).**

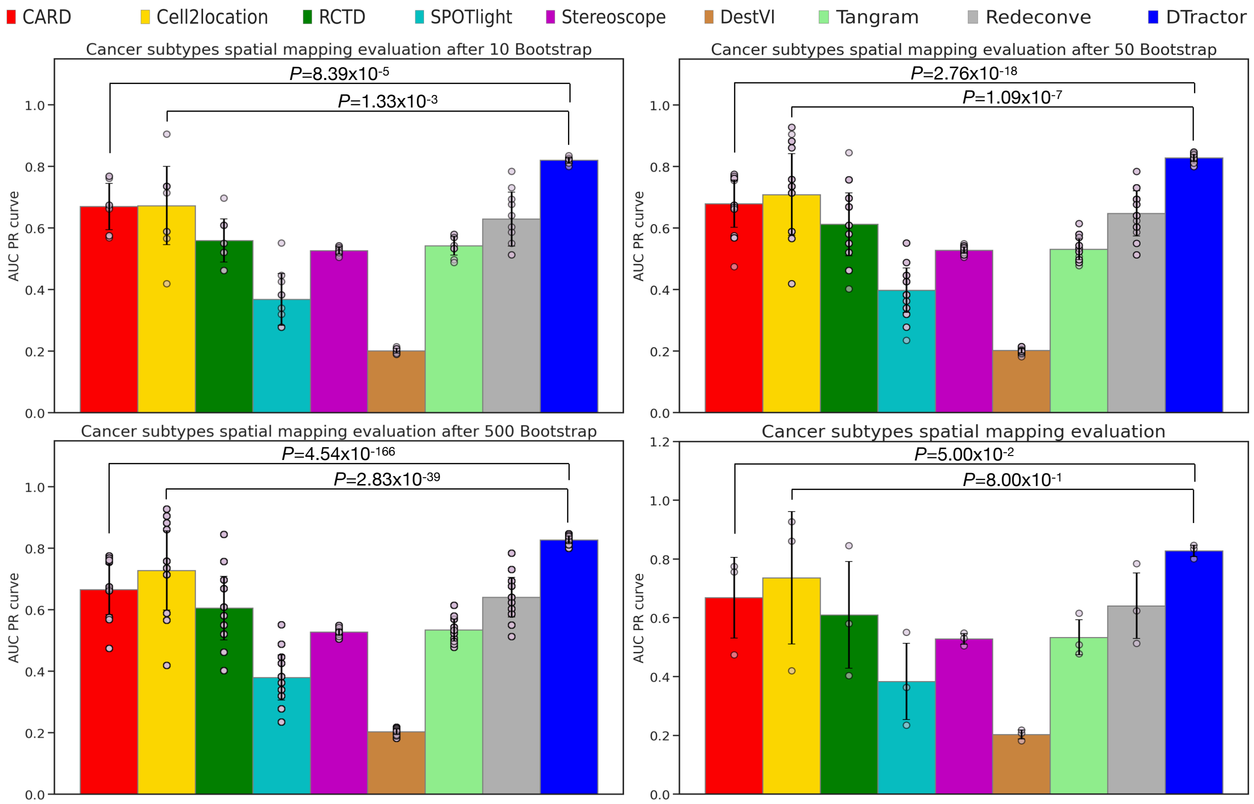


**Fig. S23 Bootstrap resampling for groups of 10, 50, and 500 samples, alongside the original plot without resampling, to assess the spatial mapping accuracy of cancer subtypes based on ground truth annotations in PDAC. P-values are derived from one-sided Mann-Whitney U tests without adjustment. The height of each bar represents the average precision-recall scores area under the curve (AUC PR), while each scatter dot represents a different sample and the error bars indicate the standard errors.**

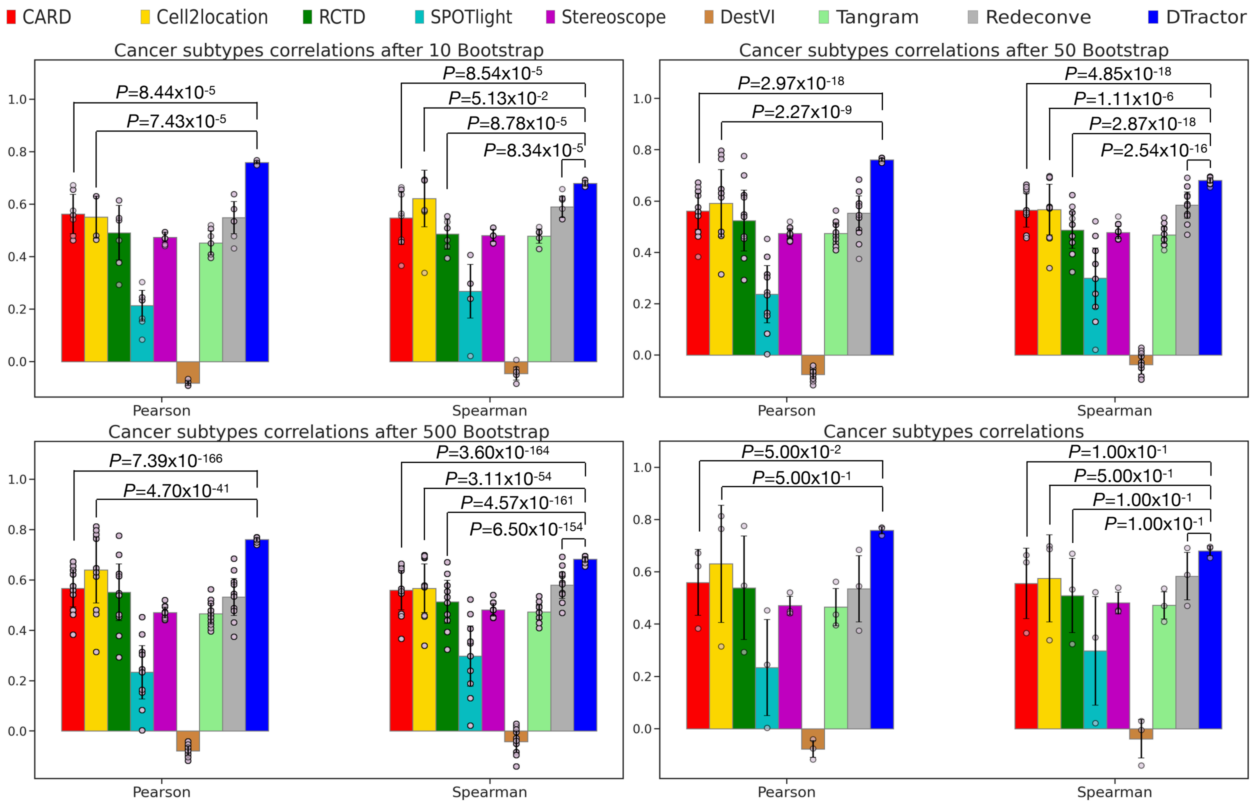


**Fig. S24 Bootstrap resampling for groups of 10, 50, and 500 samples, alongside the original plot without resampling, to assess macro-average correlation coefficients between the estimated cell type composition matrix of cancer-associated cell types and the cancer ground truth annotation in PDAC. P-values are derived from one-sided Mann-Whitney U tests without adjustment. The height of each bar represents the average statistics, while each scatter dot represents a different sample and the error bars indicate the standard errors.**

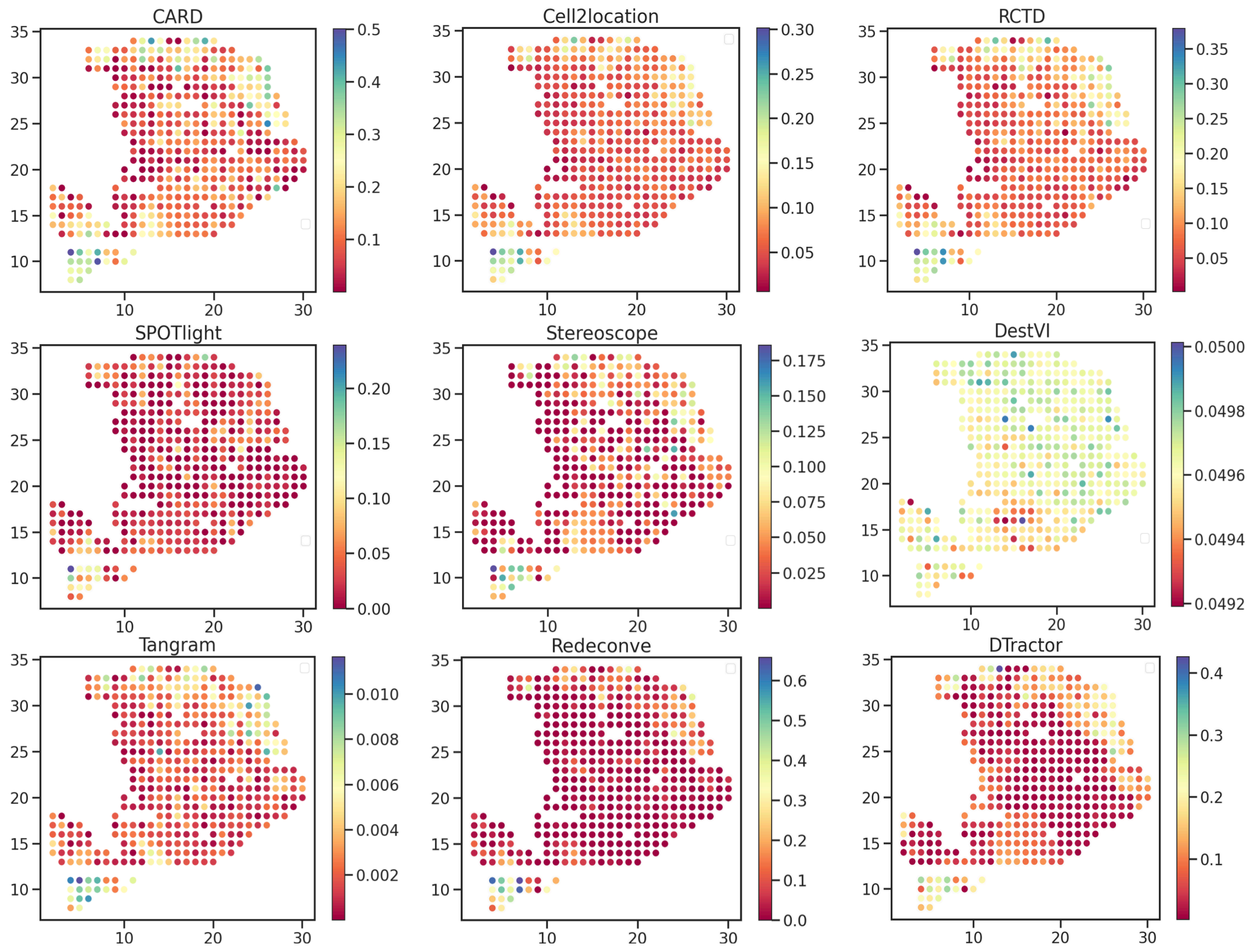


**Fig. S25 Estimated fibroblasts cell abundance (color intensity) in PDAC (13402 shared genes).**

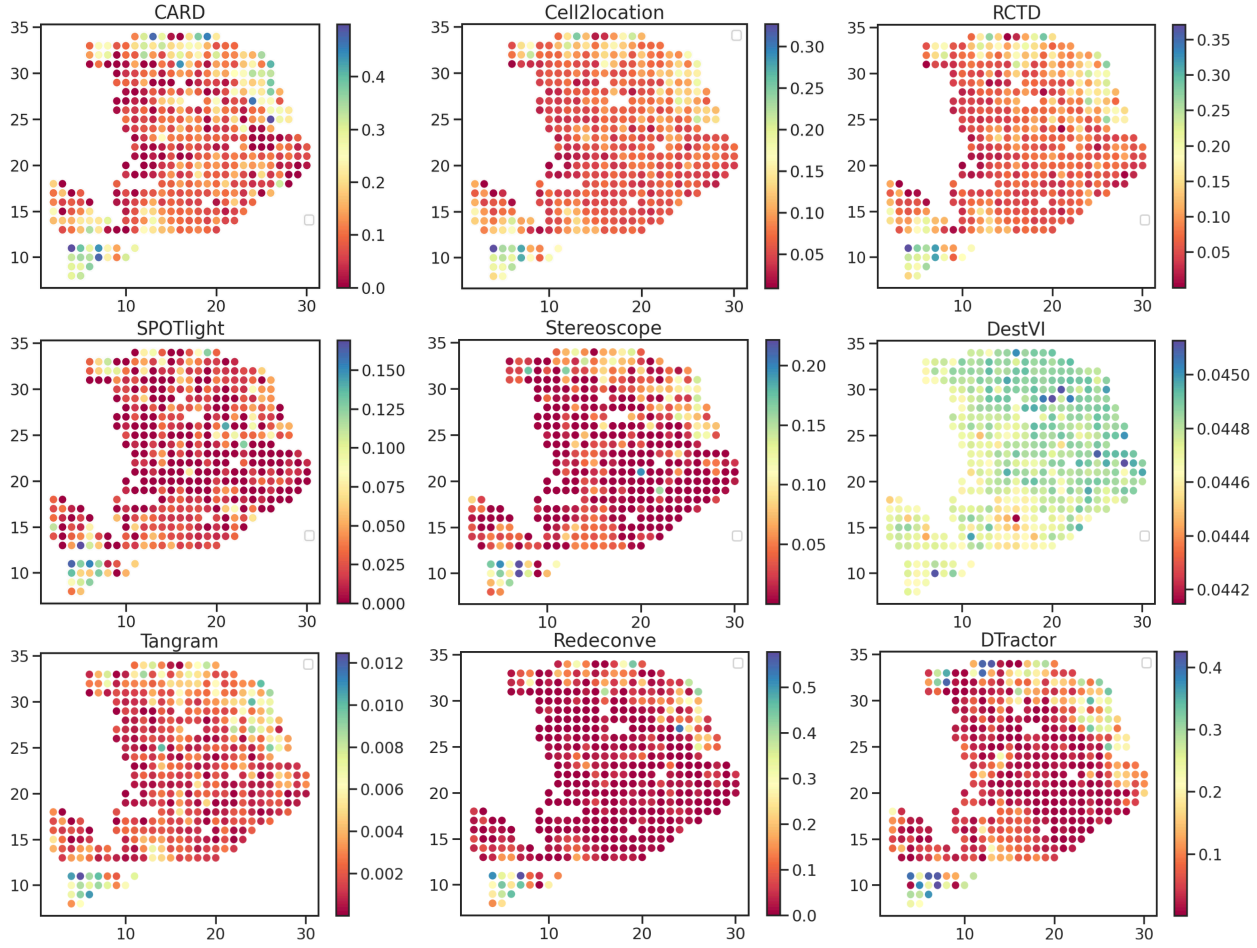


**Fig. S26 Estimated fibroblasts cell abundance (color intensity) in PDAC (9818 shared genes).**

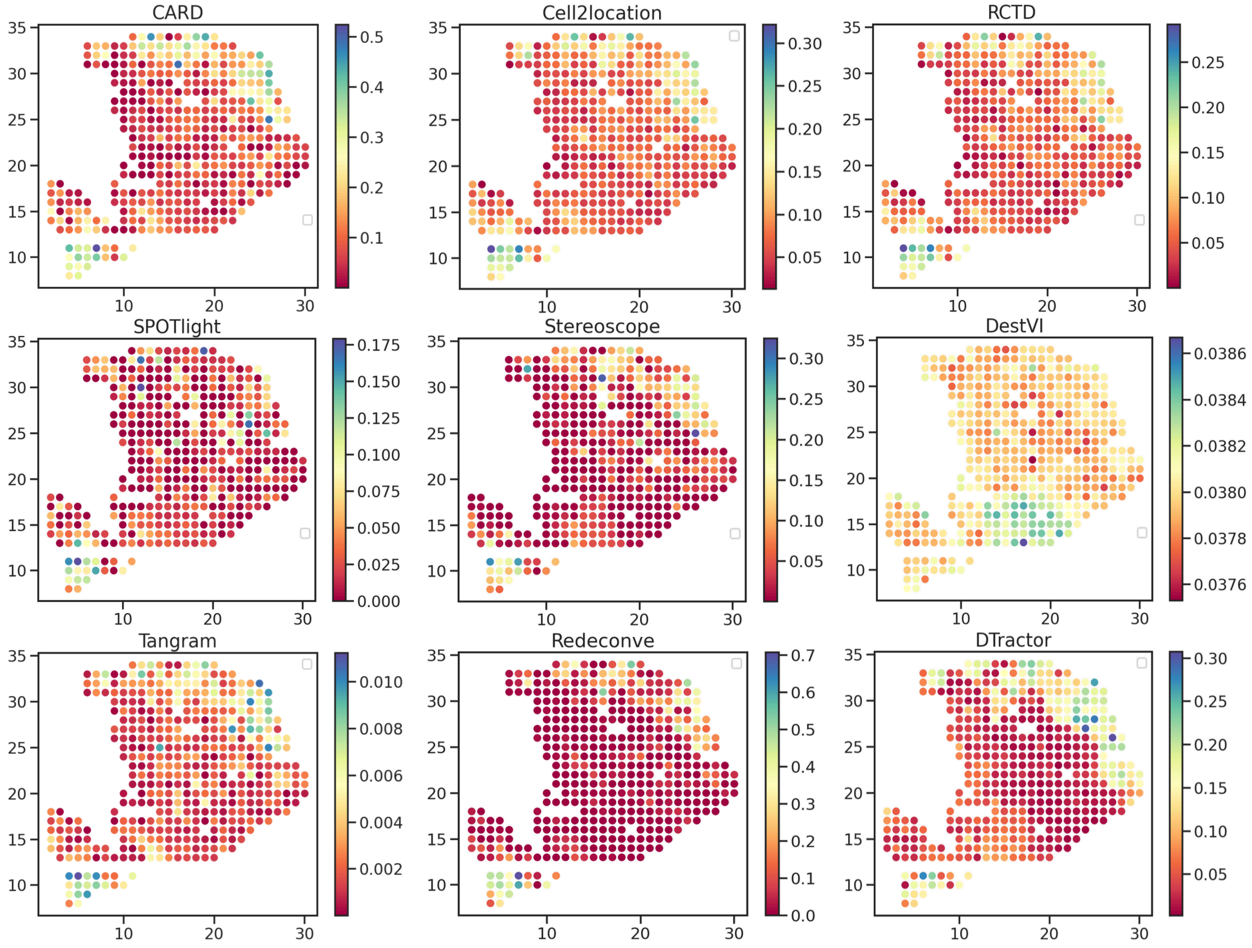


**Fig. S27 Estimated fibroblasts cell abundance (color intensity) in PDAC (8729 shared genes).**

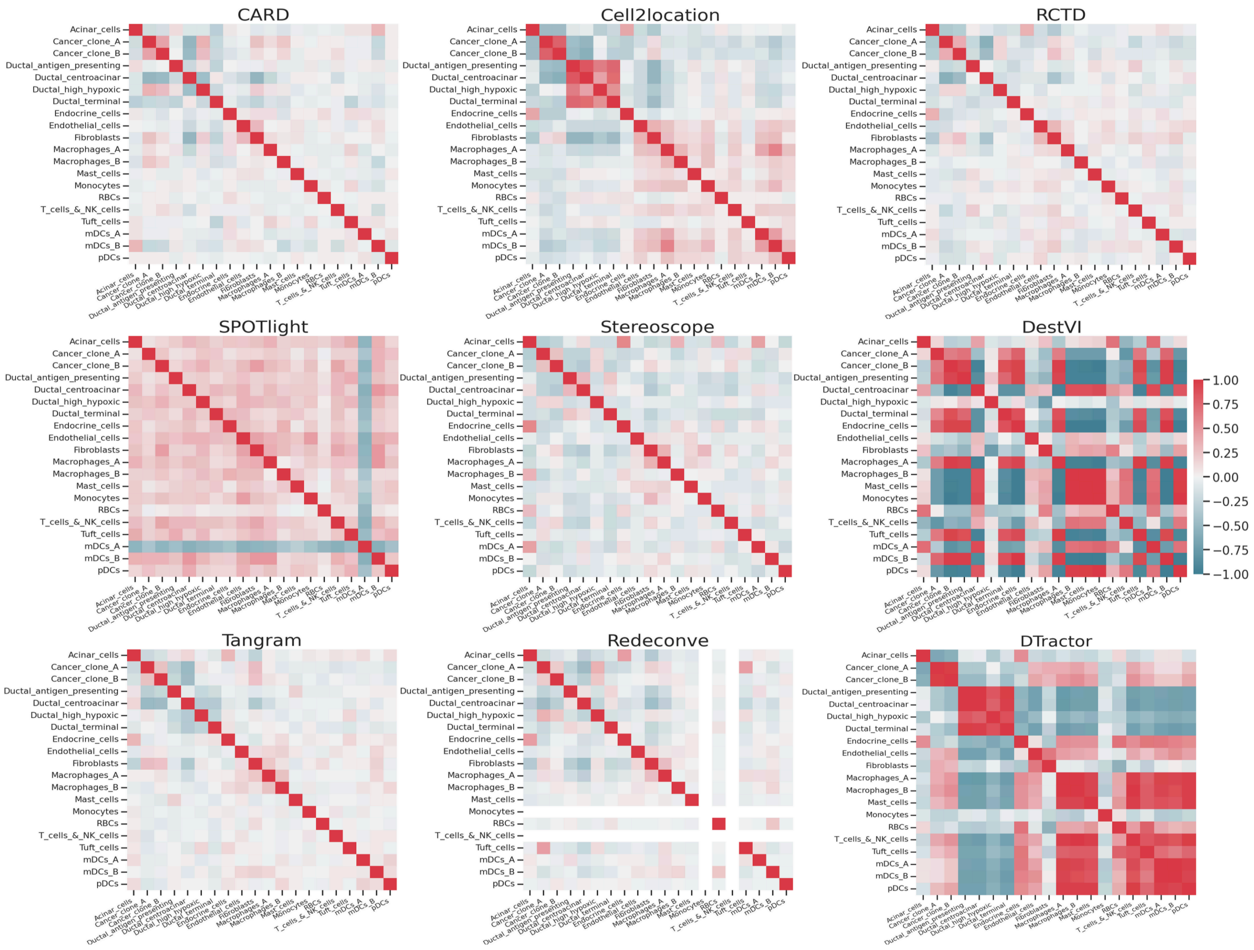


**Fig. S28** **Correlations in the estimated cell type composition matrix across spatial locations between pairs of cell types are compared across different algorithms in PDAC. The color of each comparison is scaled based on the correlation value.**

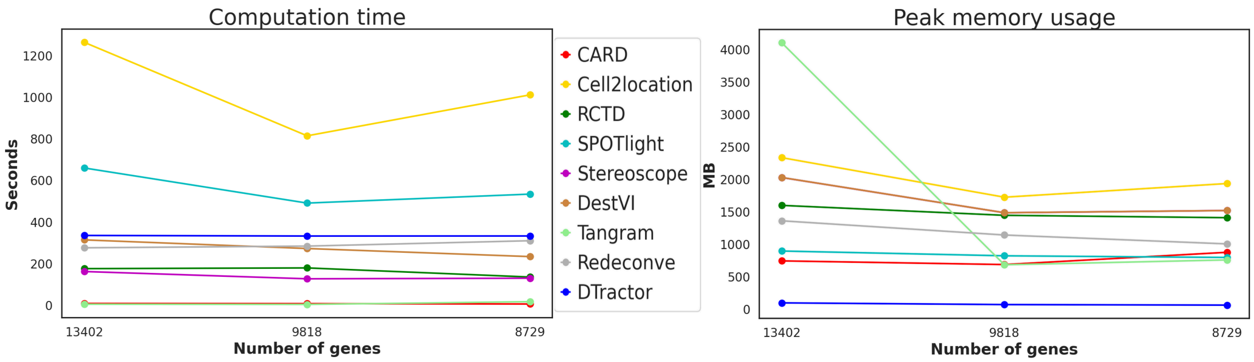


**Fig. S29** **Computation time (seconds) and peak memory usage (MB) for each deconvolution method for PDAC. To ensure fairness and consistency, we adhered to the default settings for all methods when measuring time and memory usage, as outlined in the "Parameter setting" section under "Benchmarking" in the Methods.**

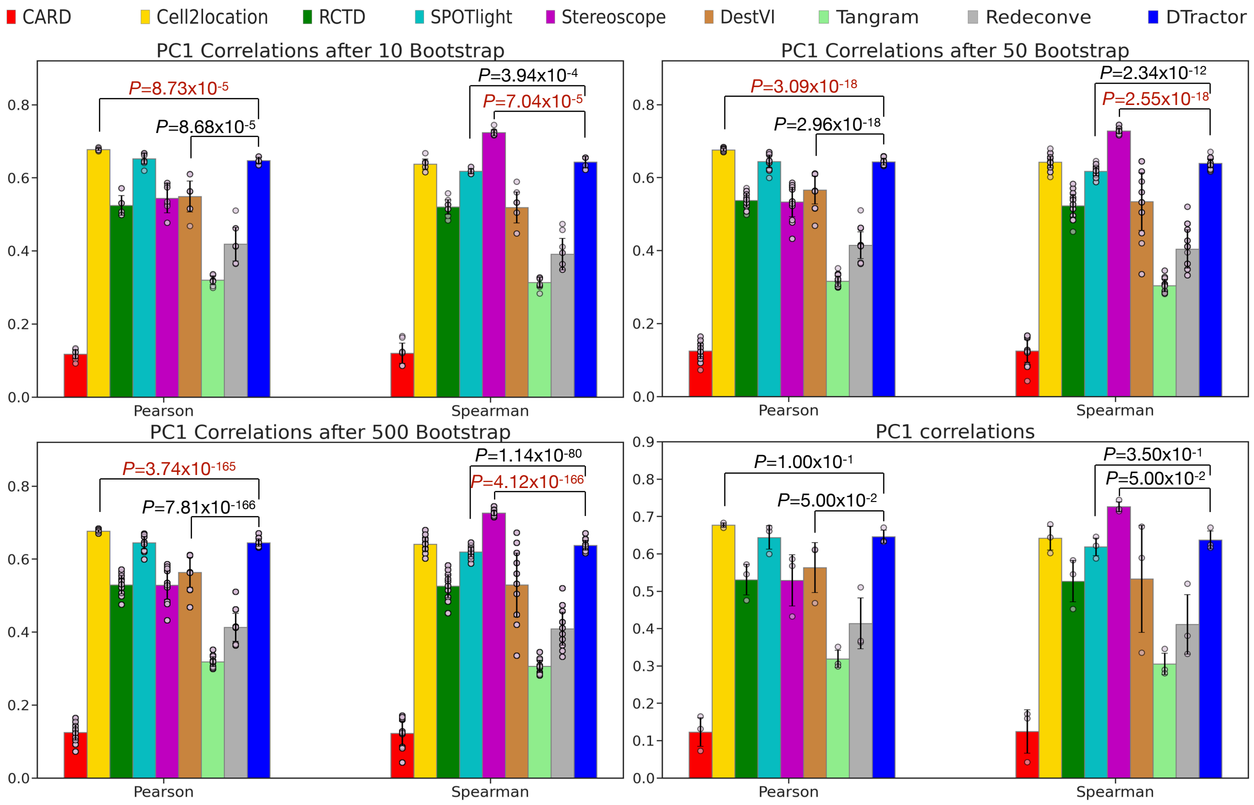


**Fig. S30 Bootstrap resampling for groups of 10, 50, and 500 samples, alongside the original plot without resampling, to assess the correlations between first principal component (PC1) scores and the entire tissue regions in the mouse olfactory bulb. P-values are derived from one-sided Mann-Whitney U tests without adjustment. Dark red indicates p-values where other models outperformed, meeting the statistical significance threshold of 0.05. The height of each bar represents the average statistics, while each scatter dot represents a different sample and the error bars indicate the standard errors.**

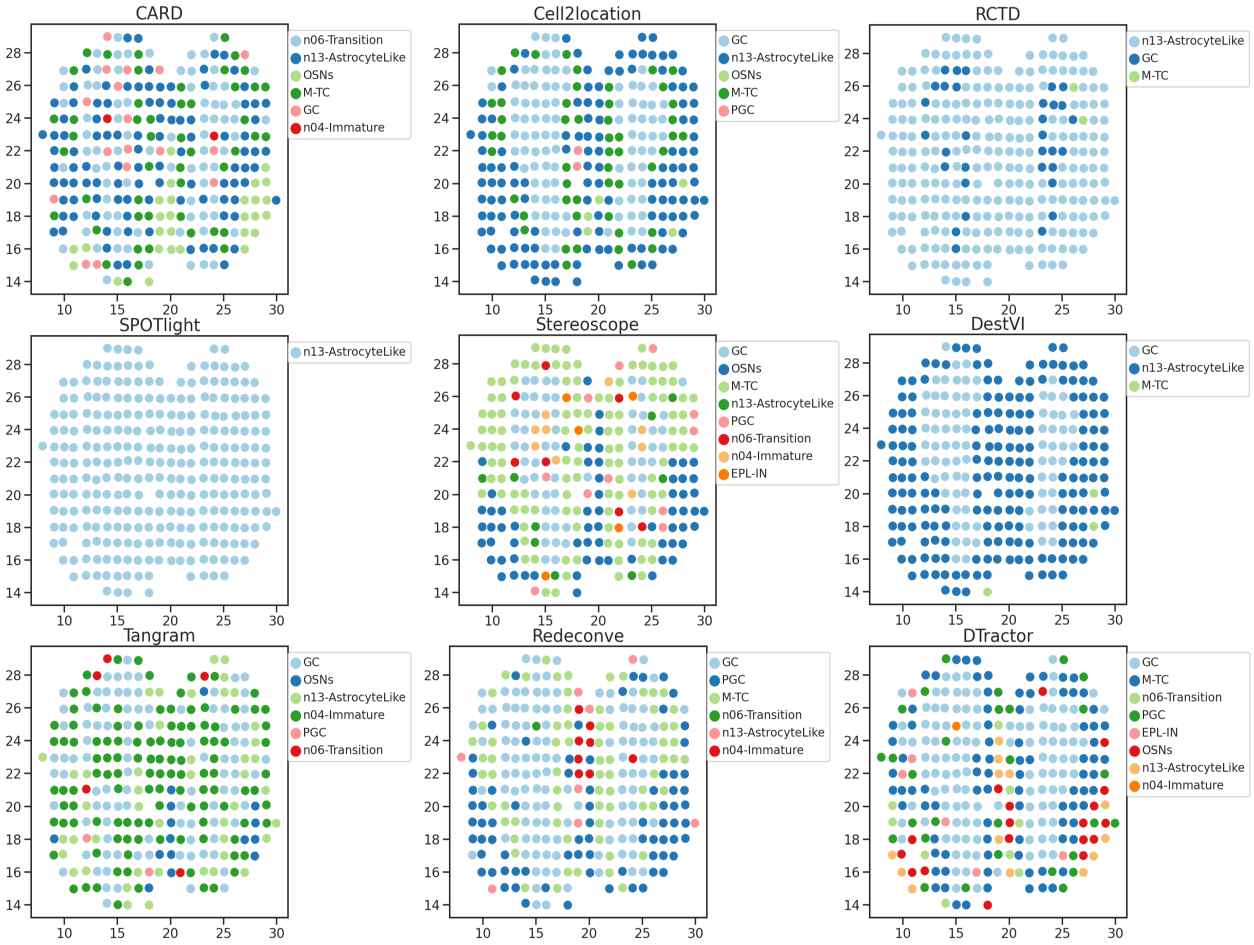


**Fig. S31** **Inferred dominant simplified cell type (granule cells (GC), periglomerular cells (PGC), olfactory sensory neurons (OSNs), mitral and tufted cells (M-TC), external plexiform layer interneurons (EPL-IN), immature cells, astrocyte-like cells, and transitional cells) of the estimated cell type composition for each location in the mouse olfactory bulb.**

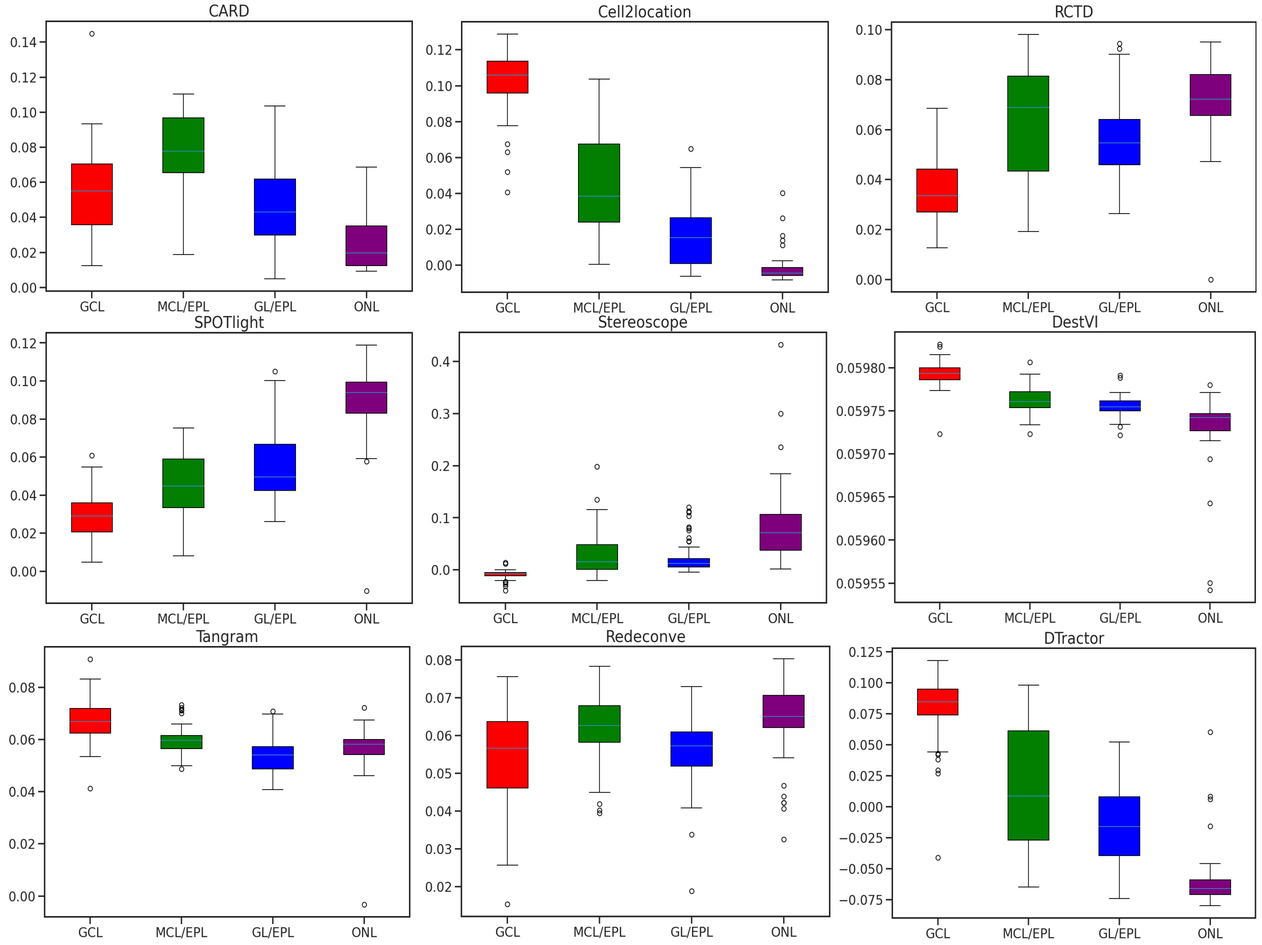


**Fig. S32** **Boxplot of the first principal component (PC1) scores of the estimated cell type composition matrix to discern how the PC1 differentiates between the four tissue regions, aligning with the ground truth annotations for the mouse olfactory bulb. DTractor successfully distinguished four different types of tissue regions consistent with histological analysis annotations. Furthermore, these boxplot demonstrates that the PC1 scores effectively capture the relative regional distances.**

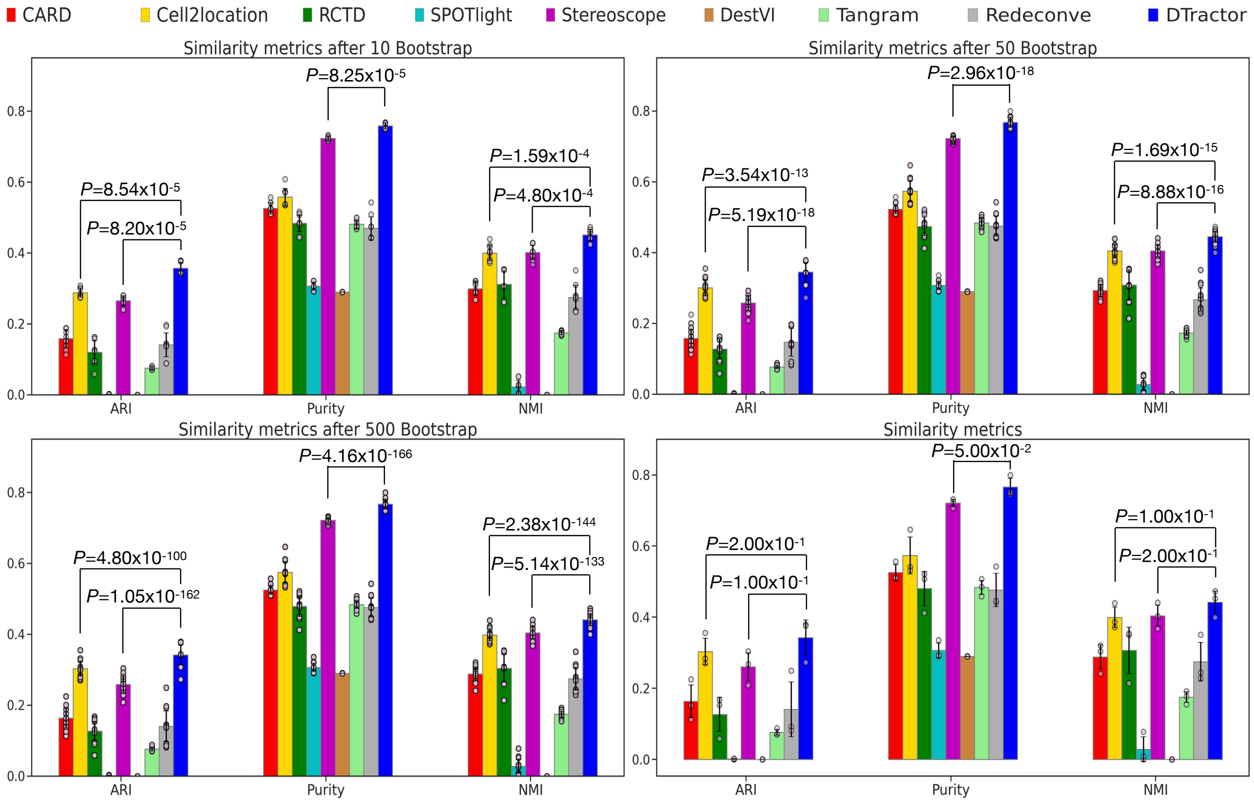


**Fig. S33** **Bootstrap resampling for groups of 10, 50, and 500 samples, alongside the original plot without resampling, to assess the similarity metrics based on ground truth annotations in the mouse olfactory bulb. Pairwise accuracy between the estimated dominant cell types and the ground truth annotation using adjusted rand index (ARI), normalized mutual information (NMI), and purity score. P-values are derived from one-sided Mann-Whitney U tests without adjustment. We used dark red font to highlight p-values where other models outperformed in the original plot prior to bootstrap resampling. The height of each bar represents the average statistics, while each scatter dot represents a different sample and the error bars indicate the standard errors.**

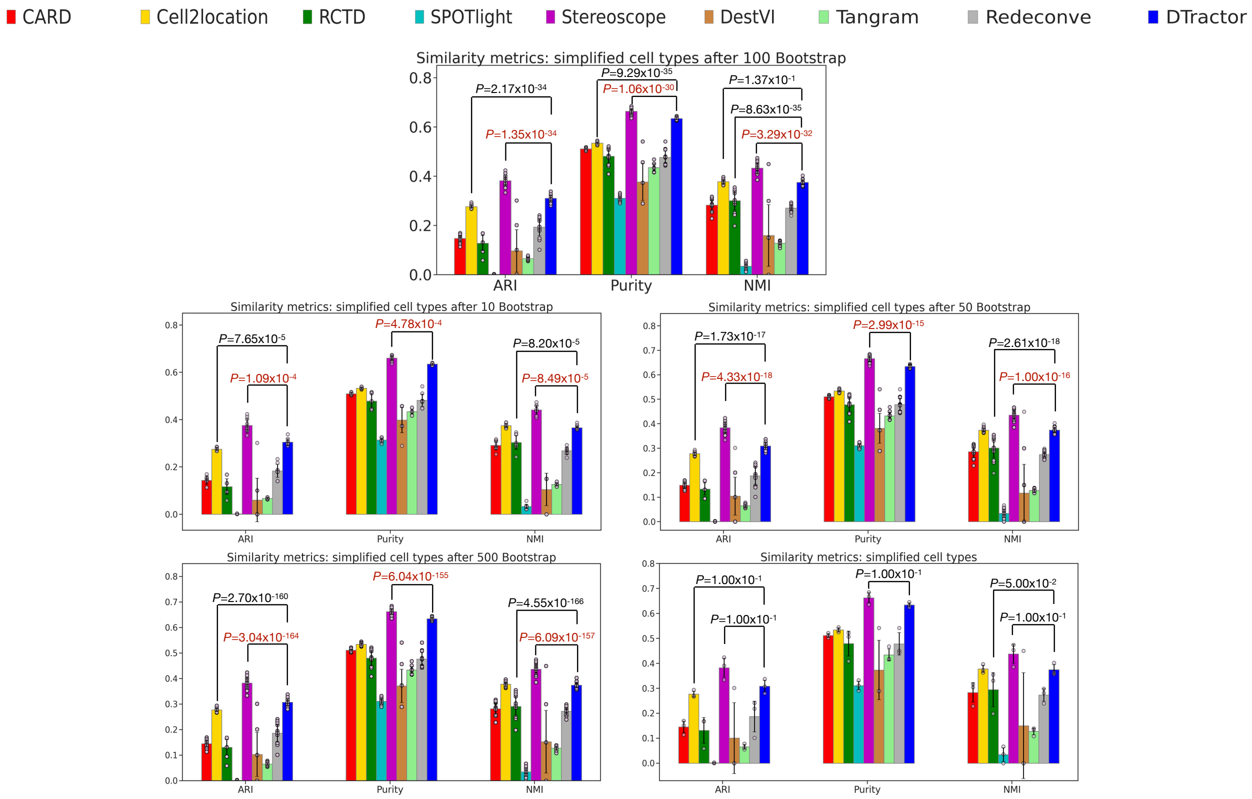


**Fig. S34 Bootstrap resampling for groups of 10, 50, 100, and 500 samples, alongside the original plot without resampling, to assess the similarity metrics for the simplified cell types (granule cells (GC), periglomerular cells (PGC), olfactory sensory neurons (OSNs), mitral and tufted cells (M-TC), external plexiform layer interneurons (EPL-IN), immature cells, astrocyte-like cells, and transitional cells) based on ground truth annotation in the mouse olfactory bulb. Pairwise accuracy between the estimated dominant cell types and the ground truth annotation using adjusted rand index (ARI), normalized mutual information (NMI), and purity score. The same procedure as in Fig. S45, but it combines each cell subtype into a larger category (e.g., seven GC cell subtypes into the GC cell type), thus obtaining similarity scores between eight major cell type categories and the ground truth. P-values are derived from one-sided Mann-Whitney U tests without adjustment. Dark red indicates p-values where other models outperformed, meeting the statistical significance threshold of 0.05. The height of each bar represents the average statistics, while each scatter dot represents a different sample and the error bars indicate the standard errors.**

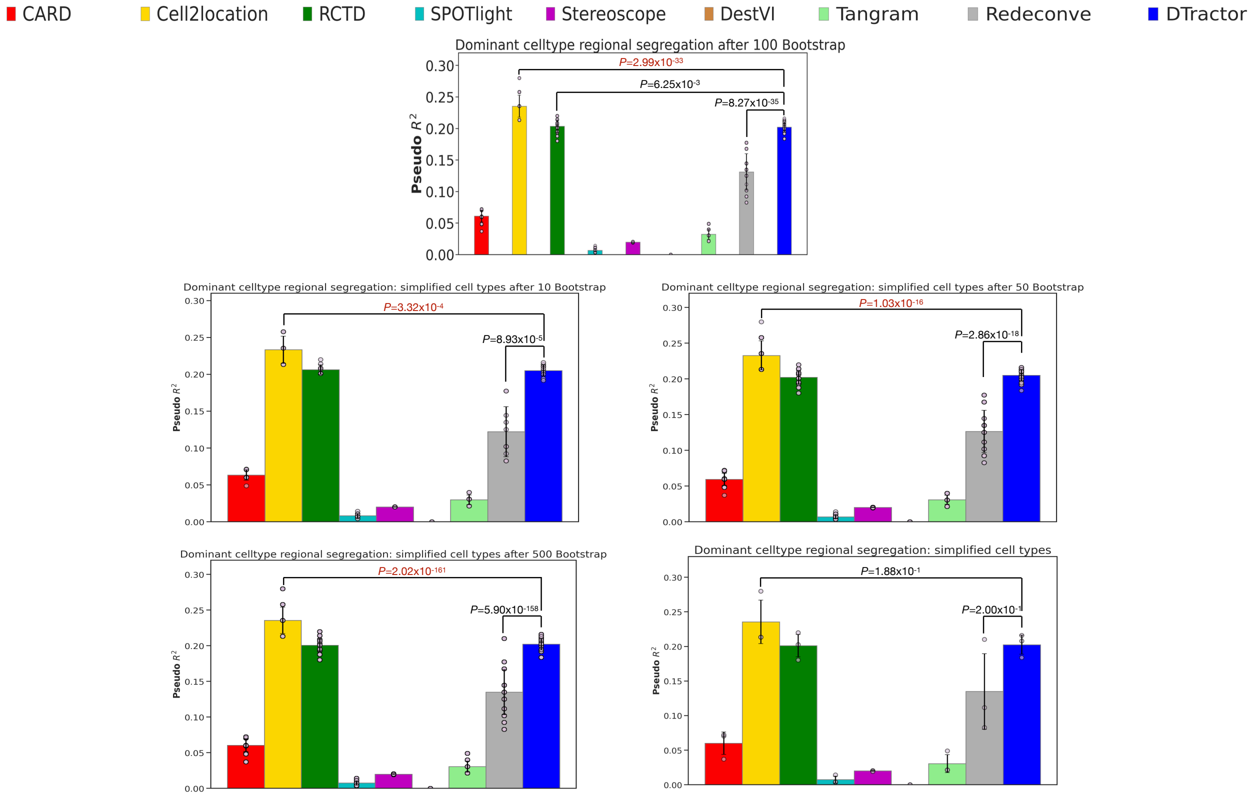


**Fig. S35 Bootstrap resampling for groups of 10, 50, 100, and 500 samples, alongside the original plot without resampling, to assess the regional segregation of the inferred dominant cell types of the estimated cell type composition matrix using the simplified cell types (granule cells (GC), periglomerular cells (PGC), olfactory sensory neurons (OSNs), mitral and tufted cells (M-TC), external plexiform layer interneurons (EPL-IN), immature cells, astrocyte-like cells, and transitional cells) in the mouse olfactory bulb. We calculated $R^2$ by fitting logistic regression models for multiclass classification across all four regions based on the inferred dominant cell types using the eight simplified cell types in the mouse olfactory bulb. P-values are derived from one-sided Mann-Whitney U tests without adjustment. Dark red indicates p-values where other models outperformed, meeting the statistical significance threshold of 0.05. The height of each bar represents the average statistics, while each scatter dot represents a different sample and the error bars indicate the standard errors.**

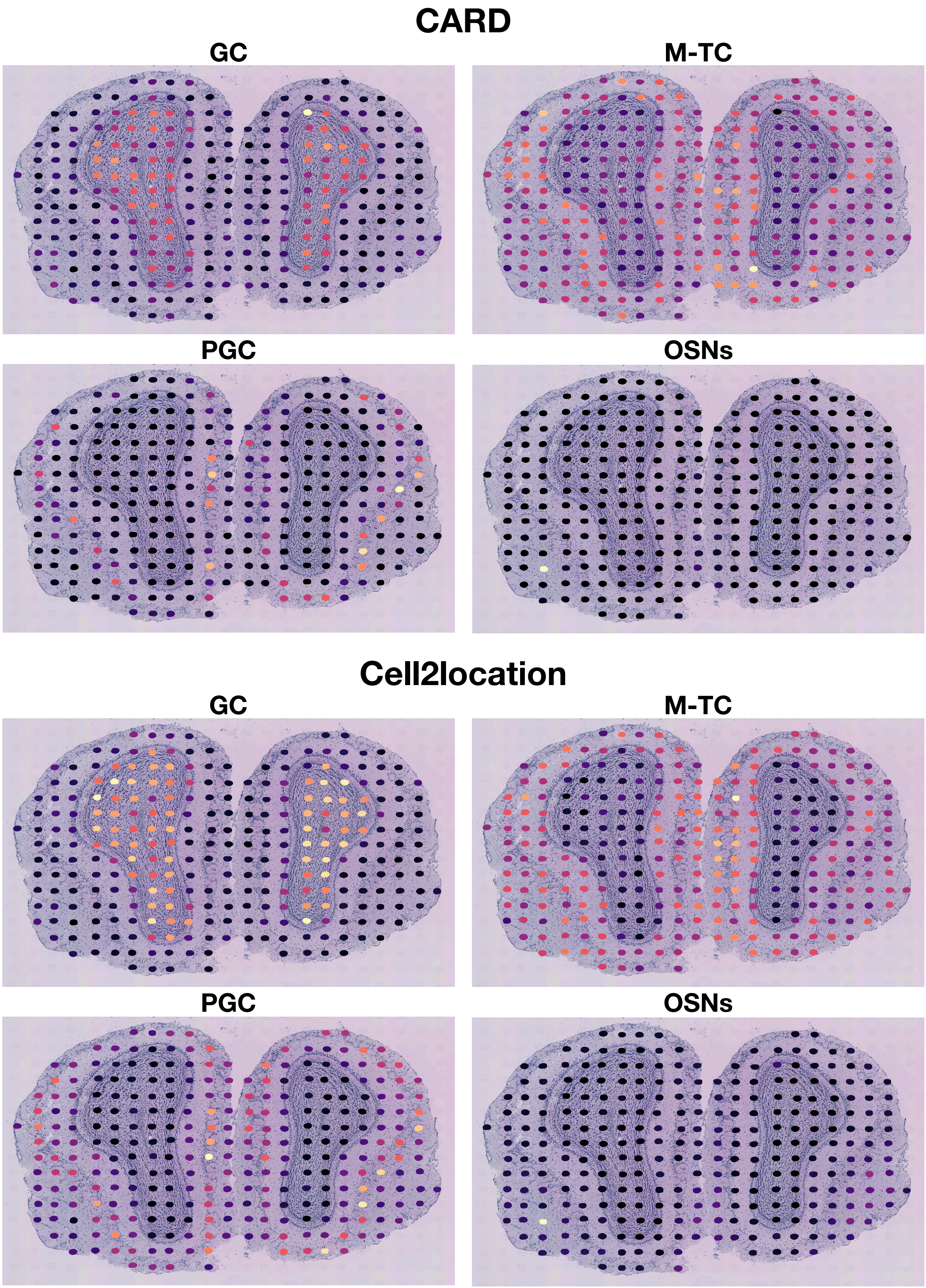
CARD
GC
M-TC
PGC
OSNs
Cell2location
GC
M-TC
PGC
OSNs

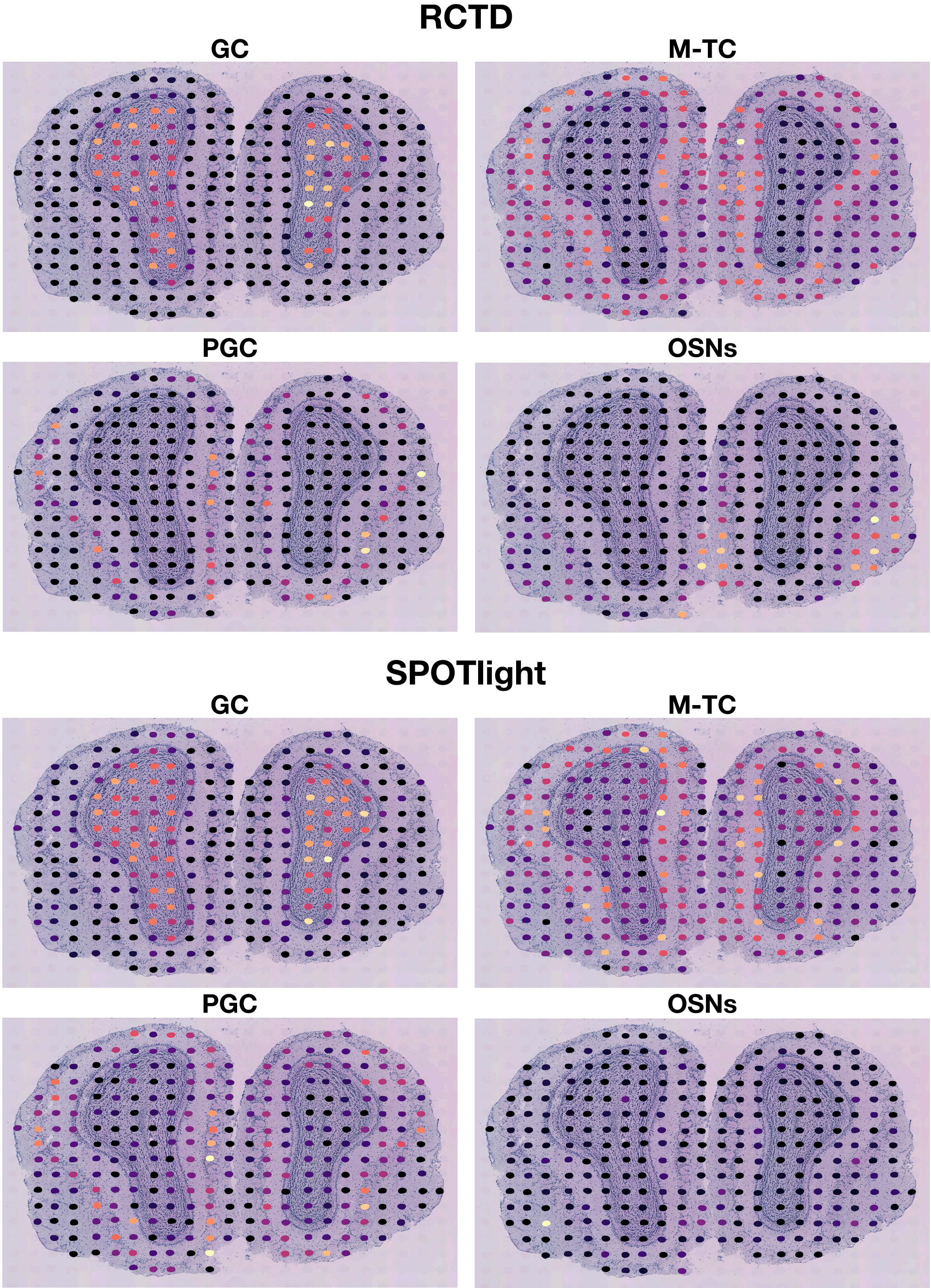
RCTD
GC
M-TC
PGC
OSNs
SPOTlight
GC
M-TC
PGC
OSNs

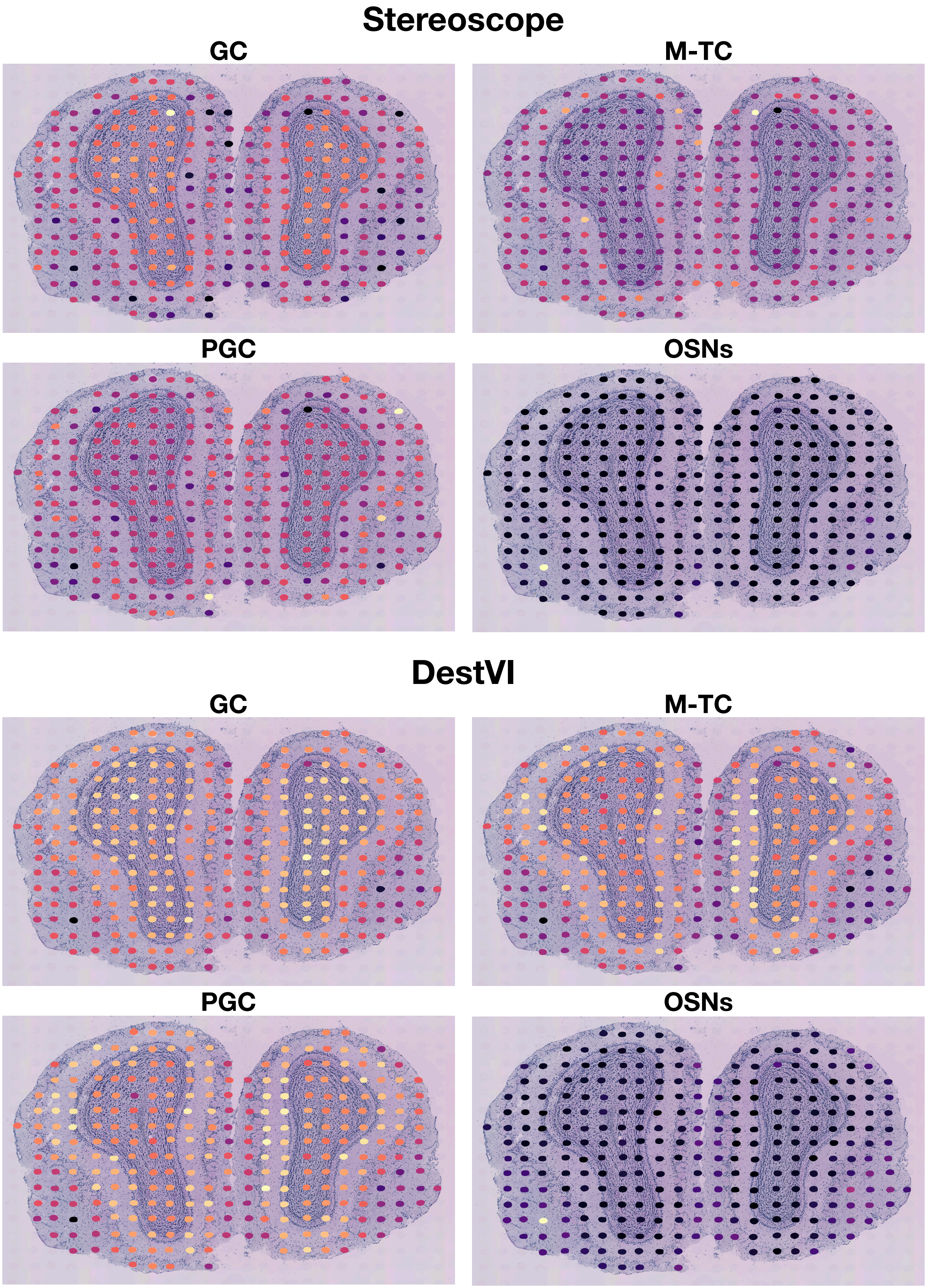
Stereoscope
GC
M-TC
PGC
OSNs
DestVI
GC
M-TC
PGC
OSNs

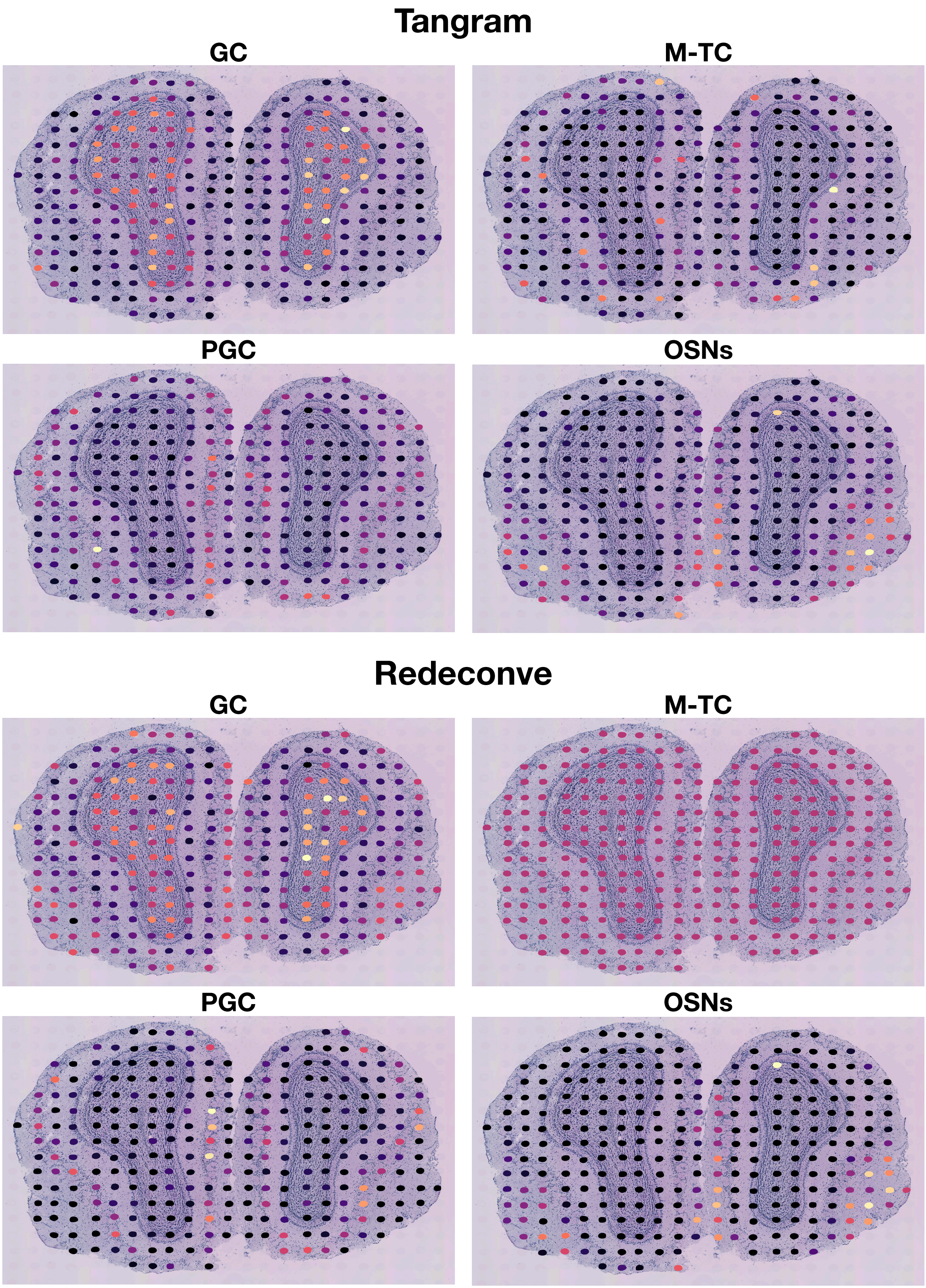
Tangram
GC
M-TC
PGC
OSNs
Redeconve
GC
M-TC
PGC
OSNs

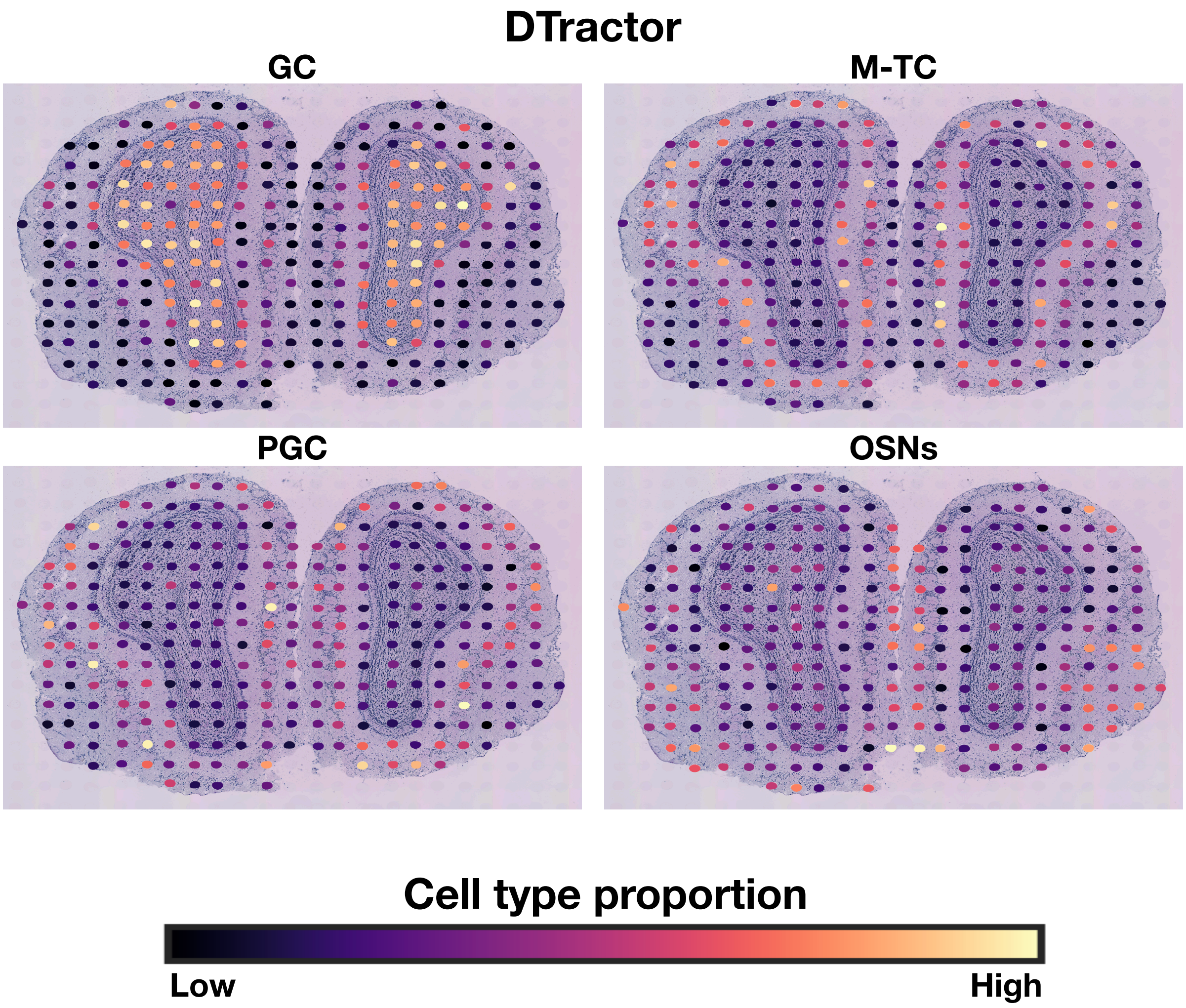


**Fig. S36** The proportion of one randomly selected example from each of the four cell types (granule cells (GC), mitral and tufted cells (M-TC), periglomerular cells (PGC), and olfactory sensory neurons (OSNs)) of the mouse olfactory bulb, utilizing the minimum number of shared genes (4,700).

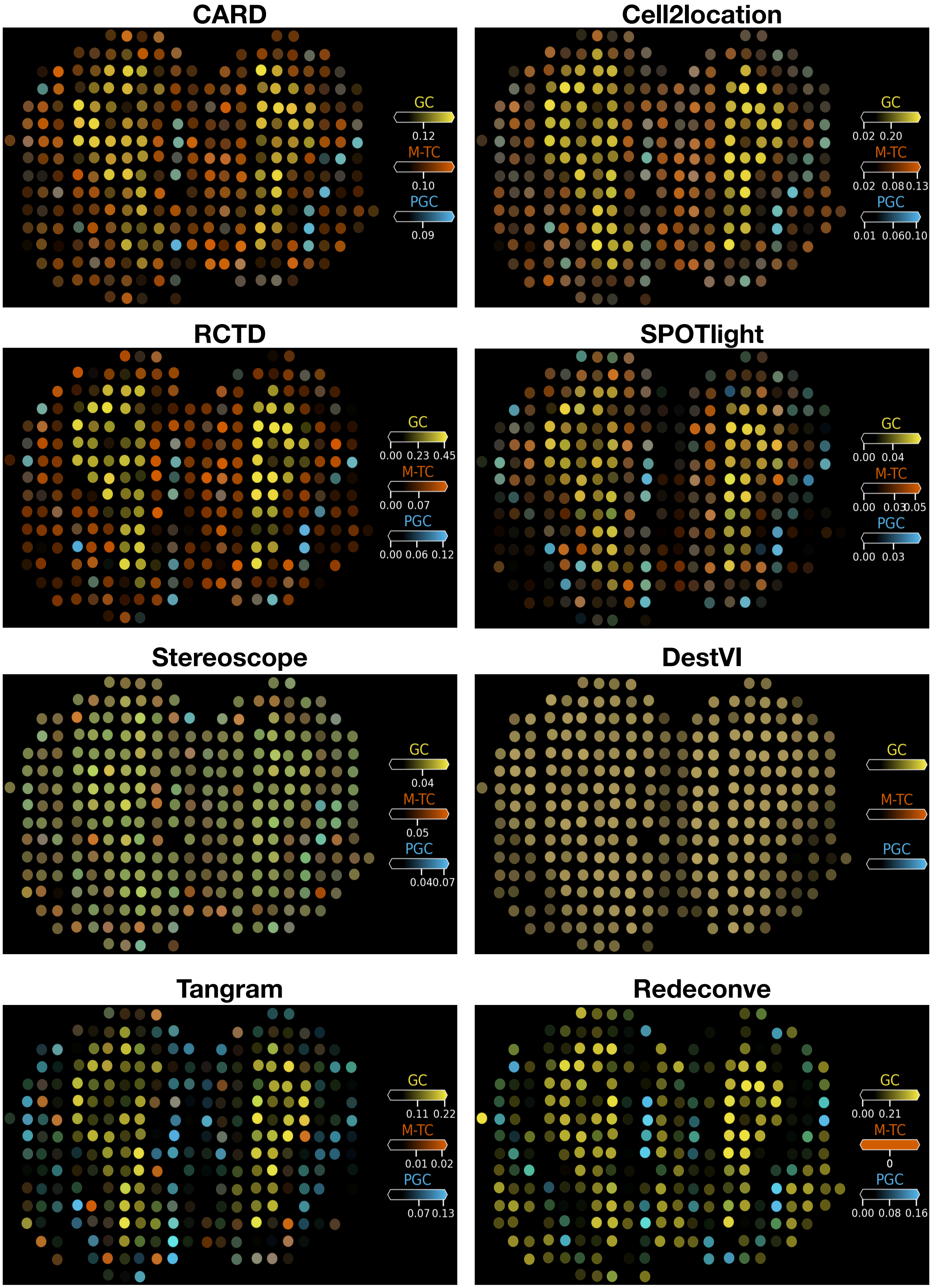
CARD
GC
0.12
M-TC
0.10
PGC
0.09
Cell2location
GC
0.02 0.20
M-TC
0.02 0.08 0.13
PGC
0.01 0.06 0.10
RCTD
GC
0.00 0.23 0.45
M-TC
0.00 0.07
PGC
0.00 0.06 0.12
SPOTlight
GC
0.00 0.04
M-TC
0.00 0.03 0.05
PGC
0.00 0.03
Stereoscope
GC
0.04
M-TC
0.05
PGC
0.04 0.07
DestVI
GC
M-TC
PGC
Tangram
GC
0.11 0.22
M-TC
0.01 0.02
PGC
0.07 0.13
Redeconve
GC
0.00 0.21
M-TC
0
PGC
0.00 0.08 0.16

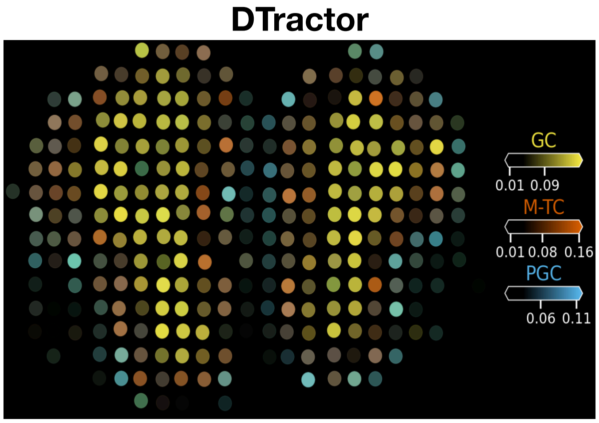


**Fig. S37 Estimated cell abundance (indicated by color intensity) of major cell types from the key tissue zones of the mouse olfactory bulb, including one randomly selected example from each of the GC, M/TC, and PGC subtypes, visualized using the Cell2location[1] plotting function, utilizing the minimum number of shared genes (4,700).**

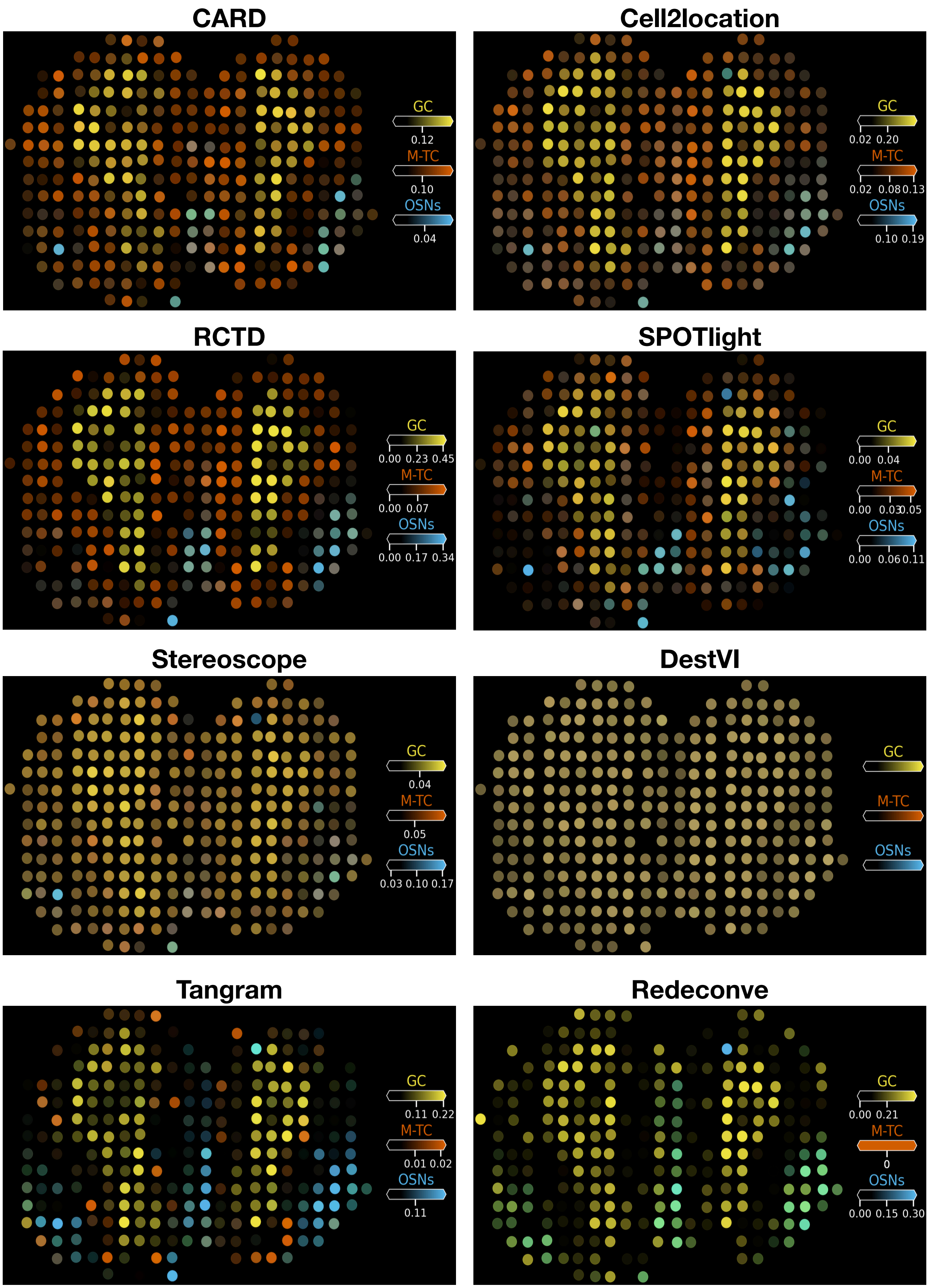
CARD
GC
0.12
M-TC
0.10
OSNs
0.04
Cell2location
GC
0.02 0.20
M-TC
0.02 0.08 0.13
OSNs
0.10 0.19
RCTD
GC
0.00 0.23 0.45
M-TC
0.00 0.07
OSNs
0.00 0.17 0.34
SPOTlight
GC
0.00 0.04
M-TC
0.00 0.03 0.05
OSNs
0.00 0.06 0.11
Stereoscope
GC
0.04
M-TC
0.05
OSNs
0.03 0.10 0.17
DestVI
GC
M-TC
OSNs
Tangram
GC
0.11 0.22
M-TC
0.01 0.02
OSNs
0.11
Redeconve
GC
0.00 0.21
M-TC
0
OSNs
0.00 0.15 0.30

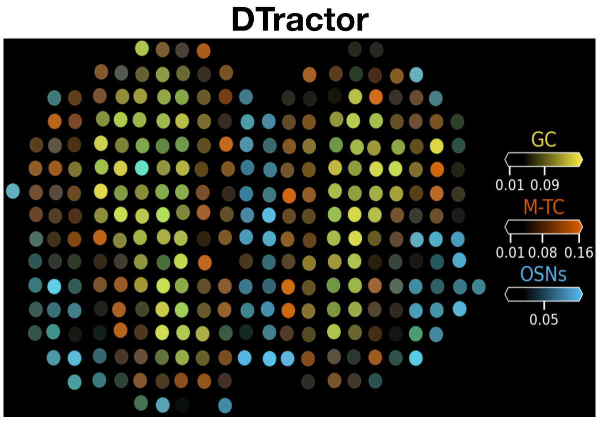


**Fig. S38** **Estimated cell abundance (indicated by color intensity) of major cell types from the key tissue zones of the mouse olfactory bulb, including one randomly selected example from each of the GC, M/TC, and OSNs subtypes, visualized using the Cell2location[1] plotting function, utilizing the minimum number of shared genes (4,700).**

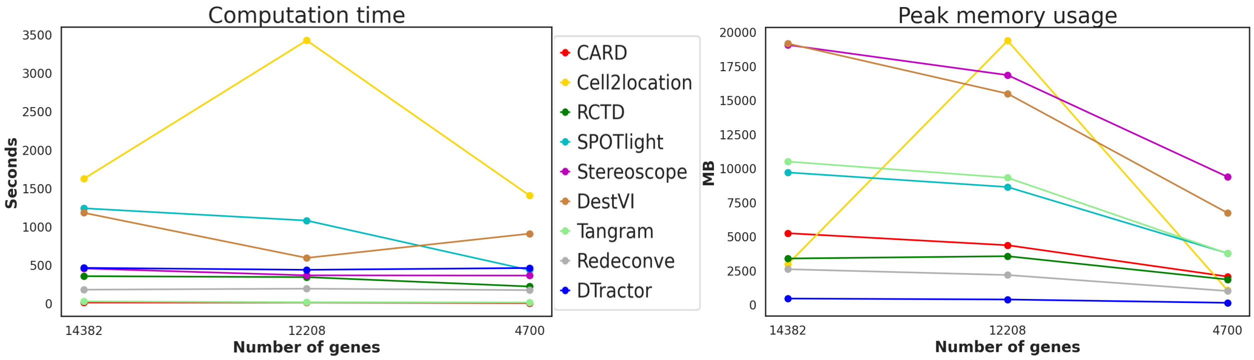


**Fig. S39** **Computation time (seconds) and peak memory usage (MB) for each deconvolution method for the mouse olfactory bulb. To ensure fairness and consistency, we adhered to the default settings for all methods when measuring time and memory usage, as outlined in the "Parameter setting" section under "Benchmarking" in the Methods.**

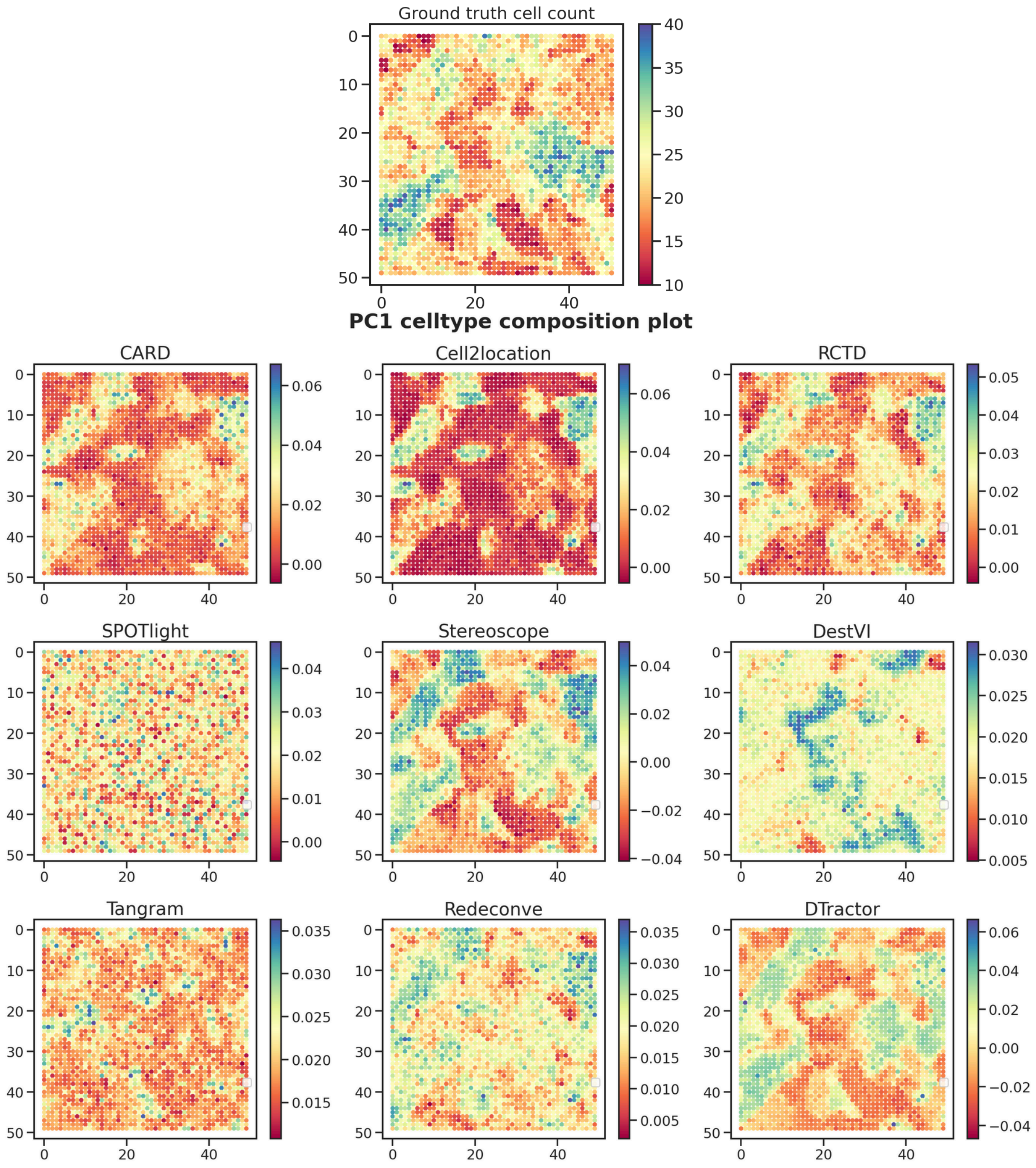


**Fig. S40** **The ground truth total cell counts and the first principal component (PC1) of the estimated cell type composition matrix for an alternative seed in the mouse brain simulation are provided to assess the robustness of each model.**

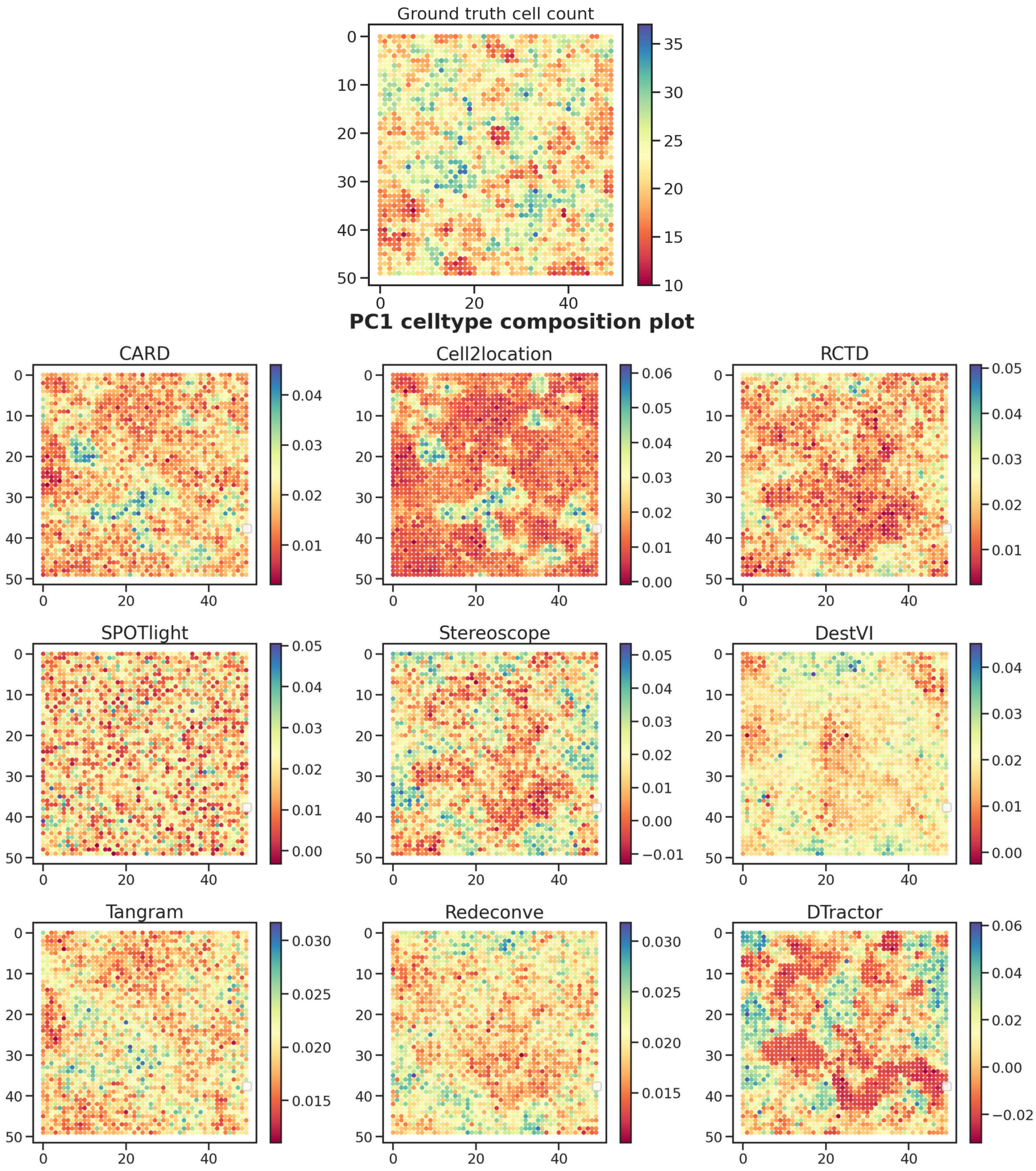


**Fig. S41** **The ground truth total cell counts and the first principal component (PC1) of the estimated cell type composition matrix for an alternative seed in the mouse brain simulation are provided to assess the robustness of each model.**

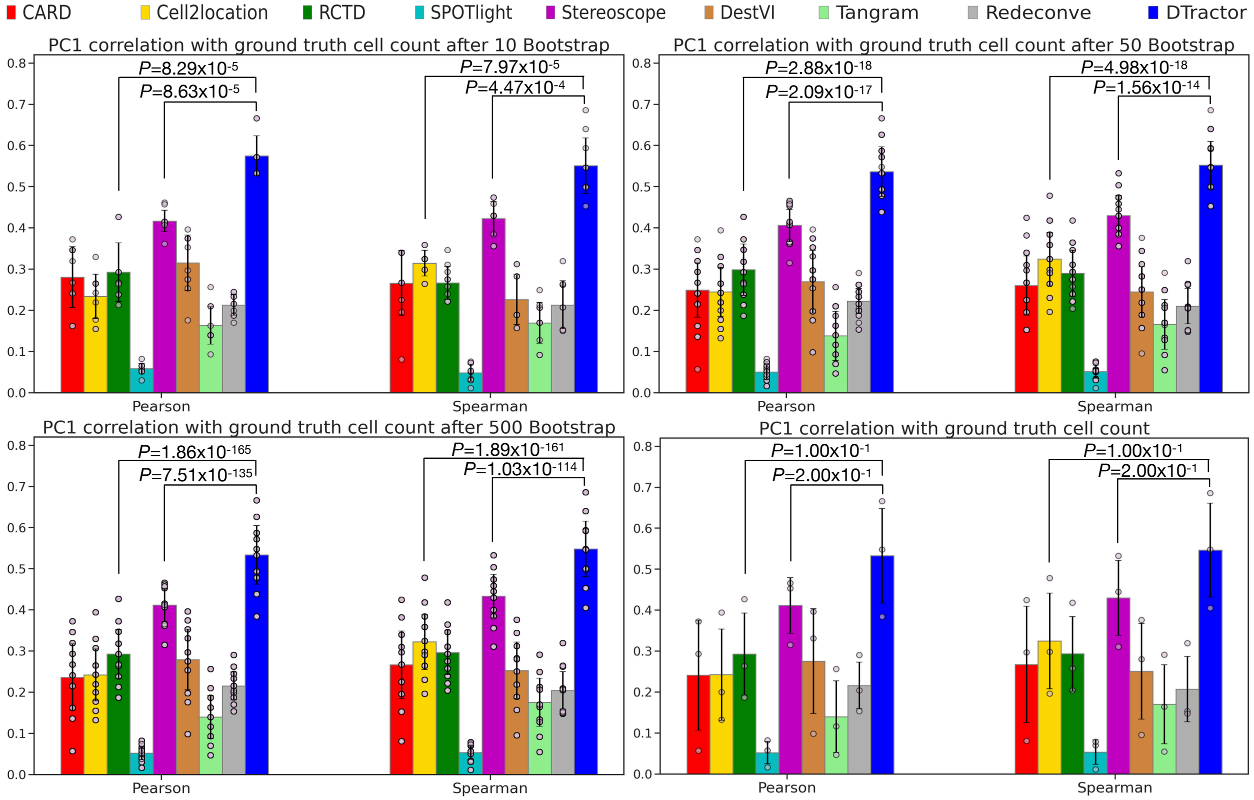


**Fig. S42 Bootstrap resampling for groups of 10, 50, and 500 samples, alongside the original plot without resampling, to evaluate the correlation between first principal component (PC1) scores and the ground truth cell count in the mouse brain simulation. P-values are derived from one-sided Mann-Whitney U tests without adjustment. The height of each bar represents the average statistics, while each scatter dot represents a different sample and the error bars indicate the standard errors.**

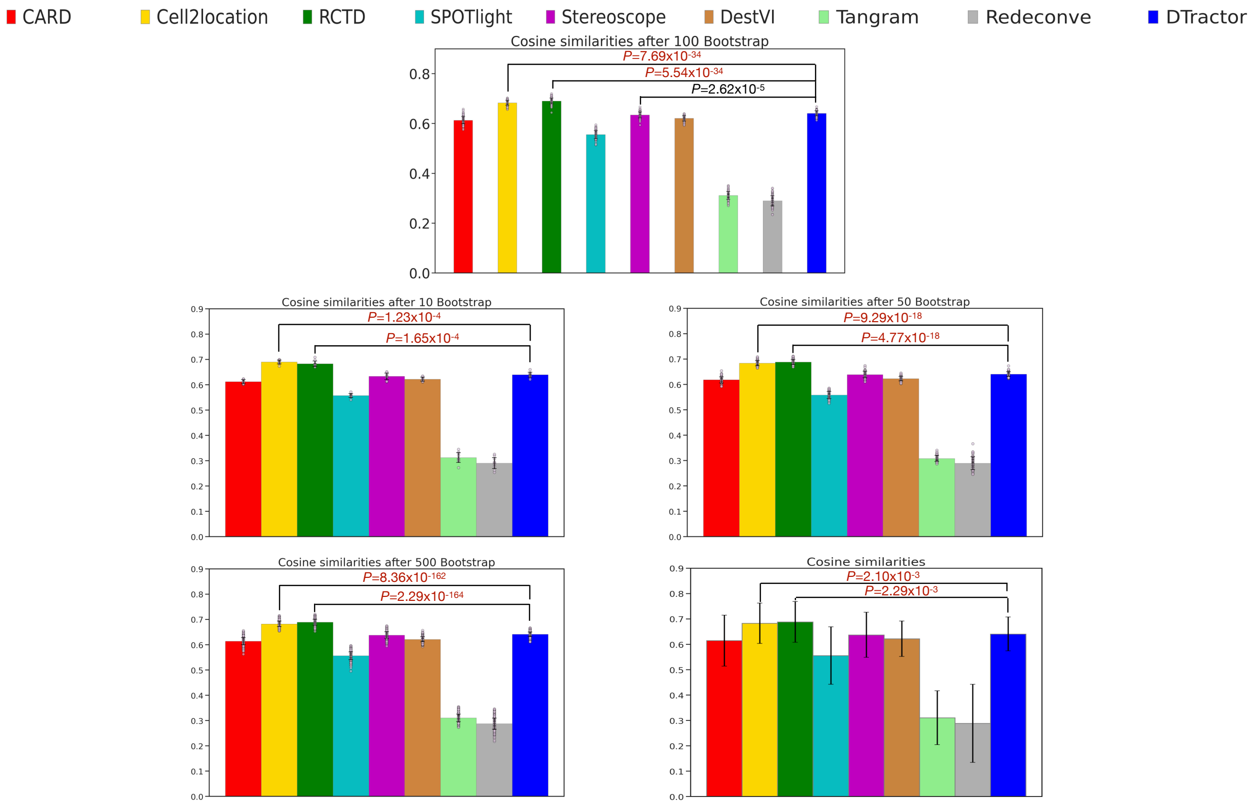


**Fig. S43 Bootstrap resampling for groups of 10, 50, 100, and 500 samples, alongside the original plot without resampling, to assess the cosine similarities between the simulated cell counts and the estimated cell type composition for each of the nine methods across individual locations for all 49 cell types in the mouse brain simulation. P-values are derived from one-sided Mann-Whitney U tests without adjustment. Dark red indicates p-values where other models outperformed, meeting the statistical significance threshold of 0.05. The height of each bar represents the average statistics, while each scatter dot represents a different sample and the error bars indicate the standard errors.**

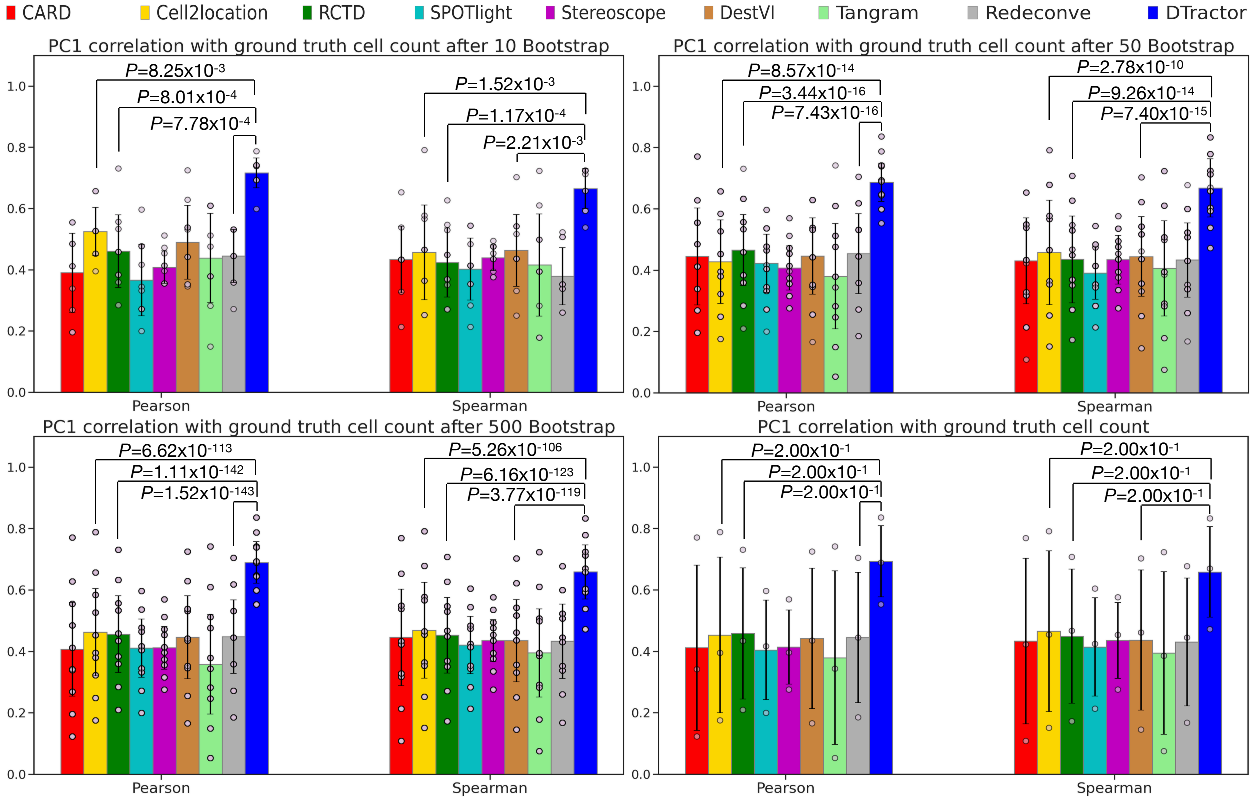


**Fig. S44 Bootstrap resampling for groups of 10, 50, and 500 samples, alongside the original plot without resampling, to assess the correlation between first principal component (PC1) scores and the ground truth cell count in the murine lymph node simulation. P-values are derived from one-sided Mann-Whitney U tests without adjustment. The height of each bar represents the average statistics, while each scatter dot represents a different sample and the error bars indicate the standard errors.**

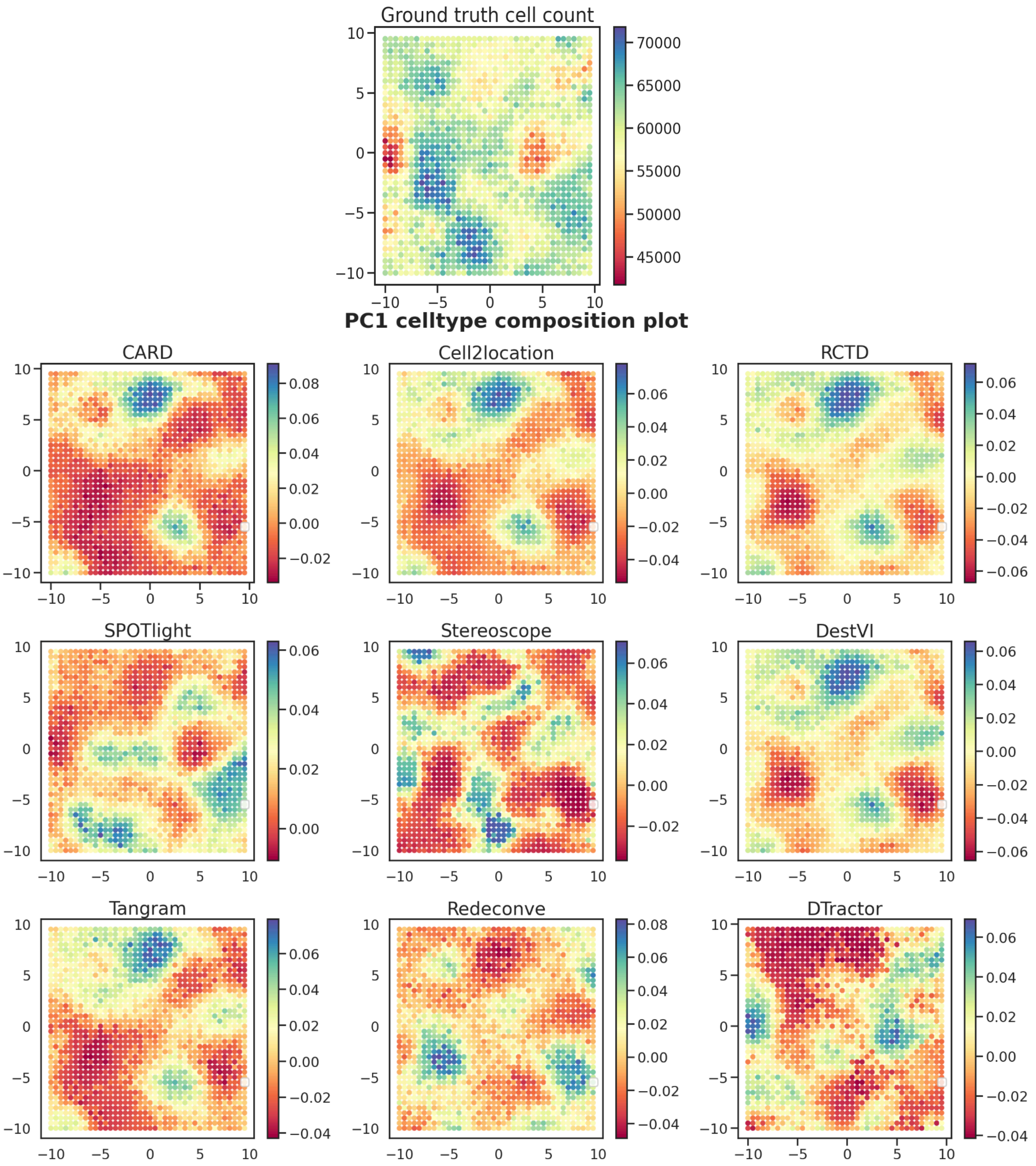


**Fig. S45** **The ground truth total cell counts and the first principal component (PC1) of the estimated cell type composition matrix for an alternative seed in the murine lymph node simulation are provided to assess the robustness of each model.**

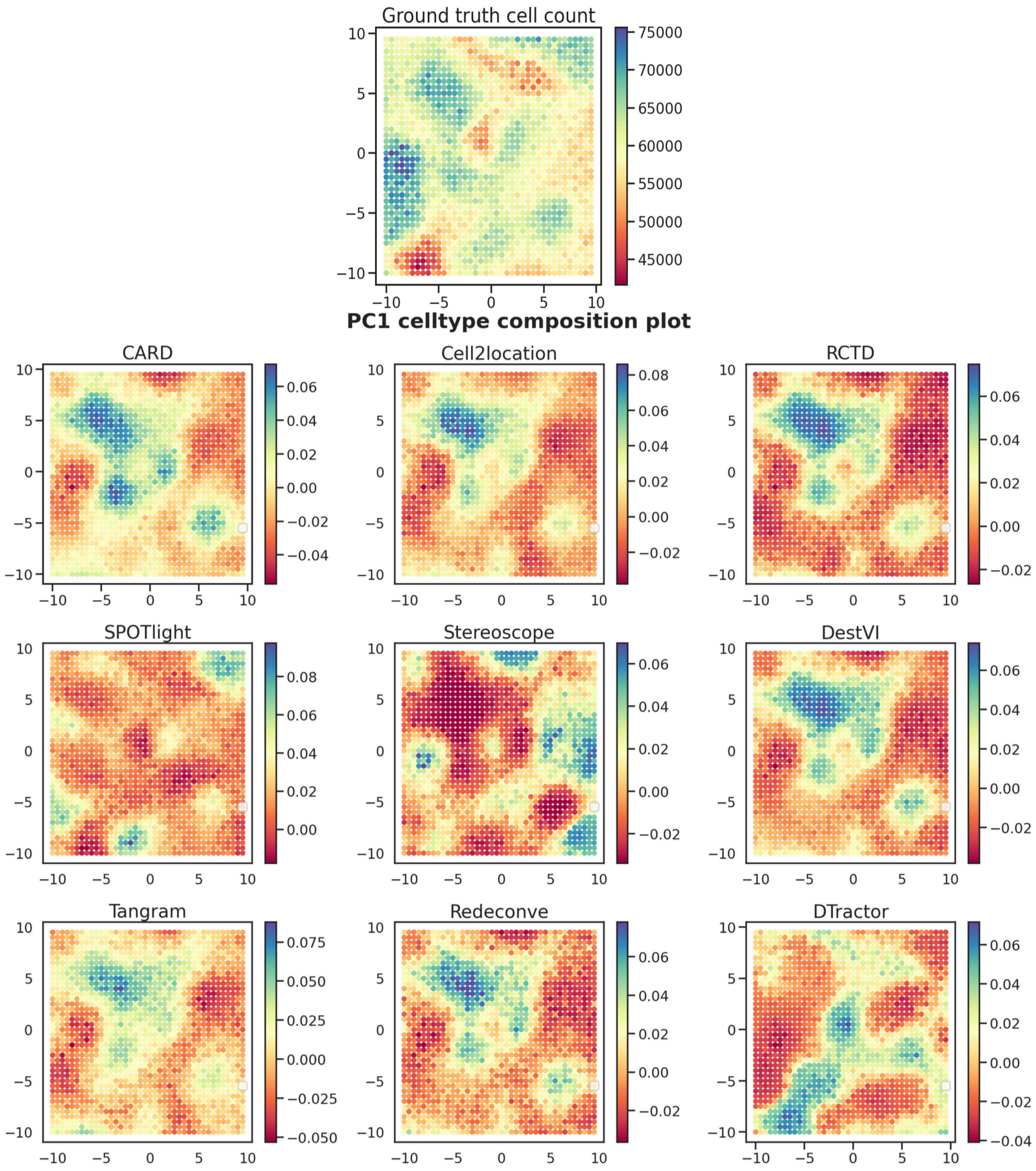


**Fig. S46** **The ground truth total cell counts and the first principal component (PC1) of the estimated cell type composition matrix for an alternative seed in the murine lymph node simulation are provided to assess the robustness of each model.**

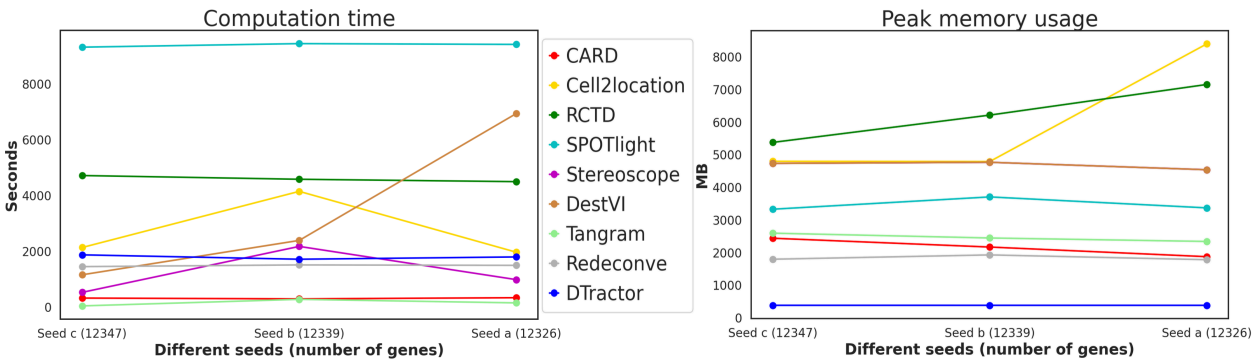


**Fig. S47** Computation time (seconds) and peak memory usage (MB) for each deconvolution method for the mouse brain simulation. To ensure fairness and consistency, we adhered to the default settings for all methods when measuring time and memory usage, as outlined in the "Parameter setting" section under "Benchmarking" in the Methods.

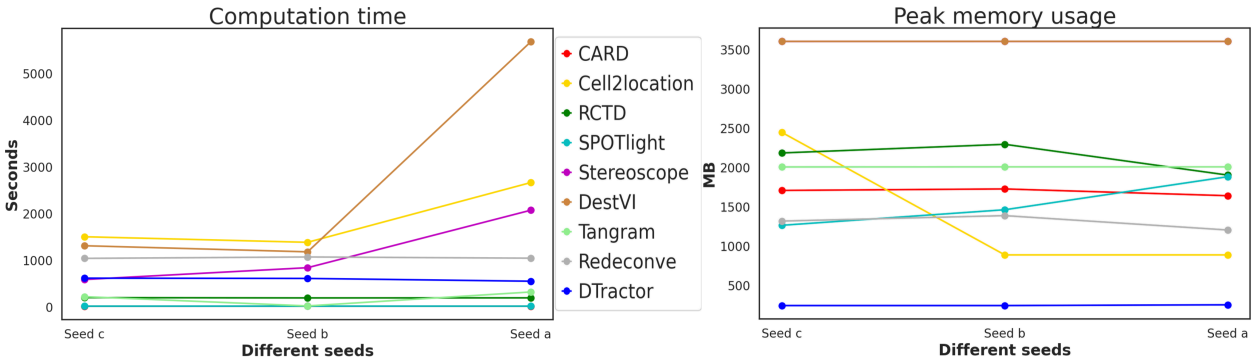


**Fig. S48** Computation time (seconds) and peak memory usage (MB) for each deconvolution method for the murine lymph node simulation. To ensure fairness and consistency, we adhered to the default settings for all methods when measuring time and memory usage, as outlined in the "Parameter setting" section under "Benchmarking" in the Methods.

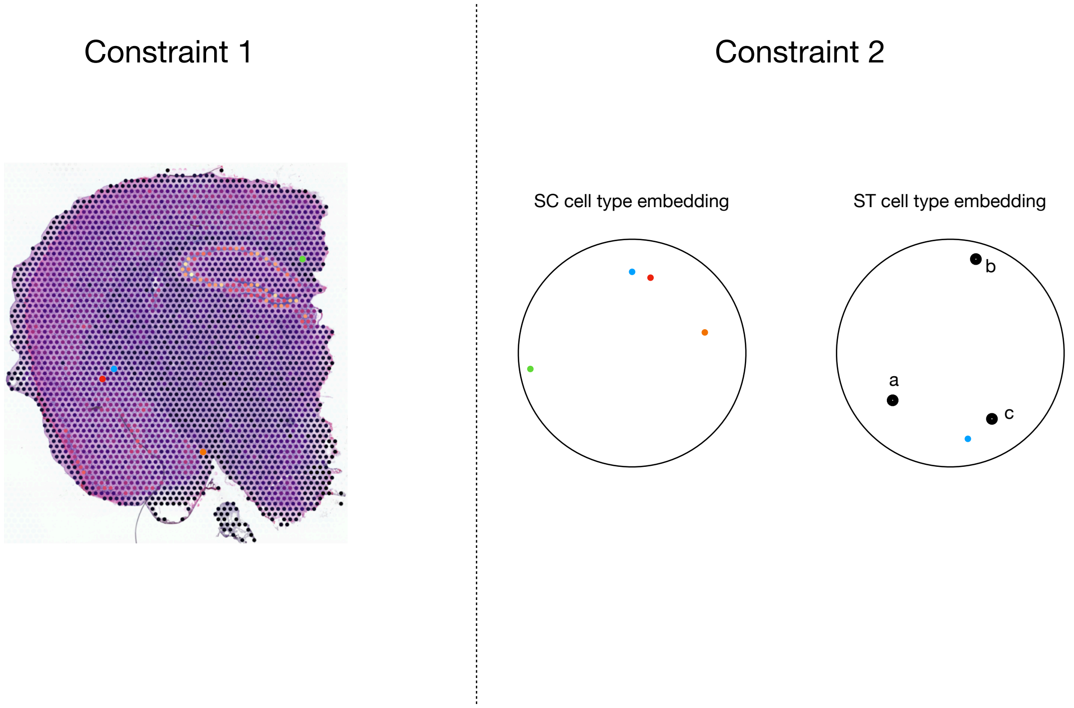


**Fig. S49 Two optional regularizations in spatial transcriptomics (ST) decomposition that can be used to extend the loss function. Two additional informative losses are introduced based on two concepts: similarity in cell-type proportions among neighboring spots and similarity between single-cell (SC) and ST cell-type embeddings**

### Supplementary Tables

| Dataset | Number of cells |
|---|---|
| Cell2location simulation (49 cell types) | Oligodendrocytes 2 / Oligo_2 (n=2011), Excitatory neurons cortical layer L23 / Ext_L23 (n=358), Excitatory neurons thalamus 1 / Ext_Thal_1 (n=273), Inhibitory neurons 4 / Inh_4 (n=269), Ext_L25 (n=260), Ext_L56 (n=257), Microglia / Micro (n=253), Excitatory neurons piriform cortex / Ext_Pir (n=245), Inh_1 (n=239), Excitatory neurons hippocampal regions dentate gyrus 2 / Ext_Hpc_DG2 (n=238), Ext_L5_1 (n=216), Unknown 1 / Unk_1 (n=188), Oligodendrocyte precursor cells 1 / OPC_1 (n=186), Parvalbumin-positive inhibitory neurons / Inh_Pvalb (n=181), Excitatory neurons hippocampal regions CA1 / Ext_Hpc_CA1 (n=171), Inhibitory neurons expression Meis2 / Inh_Meis2_3 (n=162), Ext_Thal_2 (n=158), Excitatory neurons amygdala 2 / Ext_Amy_2 (n=157), Inh_3 (n=144), Somatostatin-positive inhibitory neurons / Inh_Sst (n=125), Inh_6 (n=122), Inh_Meis2_2 (n=121), Ext_L6 (n=120), Vasoactive intestinal peptide-expressing inhibitory neurons / Inh_Vip (n=108), Astrocytes cortex / Astro_CTX (n=104), Ext_Hpc_DG1 (n=101), Inh_Meis2_1 (n=91), Oligo_1 (n=89), Astrocytes hypothalamus / Astro_HYPO (n=89), Astrocytes amygdala cortex / Astro_AMY_CTX (n=84), Inh_2 (n=75), Astrocytes medial thalamus / Astro_THAL_med (n=74), Inhibitory neurons expressing LAMP5 / Inh_Lamp5 (n=67), Astro_AMY (n=62), Astrocytes lateral thalamus / Astro_THAL_lat (n=61), Ext_Hpc_CA3 (n=60), OPC_2 (n=60), Ext_L6B (n=59), Low-quality cells 2 / LowQ_2 (n=59), Ext_Amy_1 (n=53), Neuroblast 1 / Nb_1 (n=52), Ext_L5_2 (n=52), Astrocytes hippocampus / Astro_HPC (n=50), Excitatory neurons claustrum or pyramidal neurons / Ext_ClauPyr (n=45), Inh_Meis2_4 (n=44), Excitatory neurons unknown 3 / Ext_Unk_3 (n=35), Inh_5 (n=30), Nb_2 (n=29), Astrocytes white matter / Astro_WM (n=24) |
| DestVI simulation (5 cell types) | B cells, CD4 T cells, CD8 T cells, Migratory dendritic cells / Migratory DCs, Regulatory T cells / Tregs |
| Human lymph node (34 cell types) | Memory B cells / B_mem (n=13476), naive B cells / B_naive (n=8924), naive CD4+ T cells / T_CD4+_naive (n=6012), B_Cycling (n=4765), T follicular helper cells / T_CD4+_TfH (n=4690), T_CD8+_cytotoxic (n=3890), T_CD4+_TfH_GC (n=3653), B_activated (n=3575), B cells in the light zone of the germinal center / B_GC_LZ (n=3298), T_CD4+ (n=3059), regulatory T cells / T_Treg (n=2958), B cells in the dark zone of the germinal center / B_GC_DZ (n=2500), CD8+ T cells expressing CD161 / T_CD8+_CD161+ (n=2294), T_CD8+_naive (n=2253), natural killer cells / NK (n=1372), B_plasma (n=1094), follicular regulatory T cells / T_TfR (n=1065), natural killer T cells / NKT (n=896), endothelial cells / Endo (n=622), innate lymphoid cells / ILC (n=617), pre-germinal center B cells / B_preGC (n=404), T cells expressing TIM3 / T_TIM3+ (n=357), Monocytes (n=306), plasmacytoid dendritic cells / DC_pDC (n=226), interferon-producing B cells / B_IFN (n=199), conventional dendritic cells type-2 / DC_cDC2 (n=173), pro-inflammatory macrophages / Macrophages_M1 (n=121), anti-inflammatory macrophages / Macrophages_M2 (n=110), conventional dendritic cells type-1 / DC_cDC1 (n=101), follicular dendritic cells/ FDC (n=76), pre-plasma blast B cells in the germinal center / B_GC_prePB (n=74), dendritic cells expressing CCR7 / DC_CCR7+ (n=42), vascular smooth muscle cells / VSMC (n=40), Mast (n=18) |

| Dataset | Number of cells |
|---|---|
| PDAC (20 cell types) | Ductal centroacinar (n=529), Ductal terminal (n=350), antigen-presenting ductal cells expressing major histocompatibility complex class II genes and complement pathway components / Ductal antigen presenting (n=287), Ductal expressing APOL1 and hypoxia-response-related genes / Ductal high hypoxic (n=215), Cancer clone B (n=170), Cancer clone A (n=126), T & natural killer / T cells & NK cells (n=40), myeloid dendritic B cells / mDCs B (n=33), Tuft cells (n=32), Macrophages M2 / A (n=21), Macrophages M1 / B (n=19), Monocytes (n=18), red blood cells / RBCs (n=15), Mast cells (n=14), Acinar cells (n=13), plasmacytoid dendritic cells / pDCs (n=13), myeloid dendritic A cells / mDCs A (n=12), Endothelial cells (n=11), Fibroblasts (n=5), Endocrine cells (n=3) |
| Mouse olfactory bulb (18 cell types) | Immature inhibitory neurons / Immature (n=3910), granule cell-2 / GC-2 (n=3608), AstrocyteLike (n=2950), Transition (n=2085), GC-1 (n=1914), GC-5 (n=1669), olfactory sensory neurons / OSNs (n=1200), periglomerular cell-1 / PGC-1 (n=1037), Mitral and tufted cell-3 / M/TC-3 (n=927), GC-7 (n=679), PGC-2 (n=377), PGC-3 (n=279), GC-3 (n=273), GC-6 (n=237), GC-4 (n=234), external plexiform layer interneuron / EPL-IN (n=161), M/TC-2 (n=116), M/TC-1 (n=90) |

**Table S1**: Detailed cell type information in the single-cell/nucleus datasets.

| Dataset | Number of spots | Number of genes in spatial | Number of cells | Number of genes in single-cell |
|---|---|---|---|---|
| Mouse brain simulation | 2500 | 31053 | 8111 | 31053 |
| Murine lymph node simulation | 1600 | 2000 | 32000 | 2000 |
| Human lymph node | 4035 | 36601 | 73260 | 10237 |
| PDAC | 428 | 25753 | 1926 | 19736 |
| Mouse olfactory bulb (replicate 12) | 282 | 16034 | 21746 | 18560 |

**Table S2** List of spatial transcriptomics datasets and single-cell/nucleus datasets we used in our analysis.

| Dataset | Number of spots |
|---|---|
| Human lymph node | Other (3208), GC (n=475), T-cell zone (n=352) |
| PDAC | Stroma (n=152), Duct Epithelium (n=109), Cancer (n=100), Pancreatic (n=67) |
| Mouse olfactory bulb | GL/EPL (n=83), MCL/EPL (n=74), GCL (n=69), ONL (n=56) |

**Table S3** Detailed tissue region information in the spatial datasets.
